# Pubertal sex hormones control transcriptional trajectories in the medial preoptic area

**DOI:** 10.1101/2021.09.02.458782

**Authors:** Koichi Hashikawa, Yoshiko Hashikawa, Yuejia Liu, Mark A. Rossi, Marcus L. Basiri, Jane Y. Chen, Omar R. Ahmad, Rishi V. Mukundan, Nathan L. Johnston, Jenna A. McHenry, Richard D. Palmiter, David R. Rubinow, Larry S. Zweifel, Garret D. Stuber

**Affiliations:** Center for the Neurobiology of Addiction, Pain, and Emotion, Department of Anesthesiology and Pain Medicine, Department of Pharmacology, University of Washington, Seattle, WA 98195; University of North Carolina, Chapel Hill, NC 27599; Department of Biochemistry, University of Washington, Seattle, WA 98195; Department of Psychology & Neuroscience, Duke University, Durham, NC 27708; Howard Hughes Medical Institute, University of Washington, Seattle, WA 98195; Department of Psychiatry, University of North Carolina at Chapel Hill, Chapel Hill, NC 27599; Department of Psychiatry and Behavioral Sciences, University of Washington, Seattle, WA 98195; Department of Pharmacology, University of Washington, Seattle, WA 98195

## Abstract

Pubertal maturation aids development of emotion, cognition, and reproduction. We investigated transcriptional dynamics in the medial preoptic area (MPOA), a hypothalamic center for reproductive behaviors, in male and female mice at single-cell resolution (scRNAseq) during puberty. Defined subsets of neurons expressing *Slc32a1* and *Esr1* (Vgat^+^ Esr1^+^) were the most transcriptionally dynamic compared to other cell types throughout puberty. These cell type specific transcriptional progressions towards adulthood were bidirectionally controlled by the levels of circulating testosterone and estradiol. Selective deletion of *Esr1* in *Slc32a1*-expressing cells in the MPOA prior to puberty arrested transcriptional progression and revealed a sexually dimorphic gene-regulatory network governed by Esr1. Deletion of *Esr1* in Vgat^+^ cells prevented the development of mating behavior in both sexes. These analyses reveal both sexually common and dimorphic transcriptional progressions during puberty as well as their regulatory mechanisms, which have important implications towards understanding adaptative and maladaptive processes governing adolescent brain development.

## Introduction

Puberty is a critical period when juvenile animals transition to adults via physiological maturation of the body and nervous system, which includes establishment of reproductive capabilities (Blakemore et al., 2010; Romeo et al., 2002; Schulz and Sisk, 2006). Puberty onset is initiated by activation of the HPG (hypothalamus-pituitary-gonad) axis, triggered by the excitation of gonadotropin releasing hormone (GnRH) neurons in the hypothalamus and pulsatile secretion of GnRH into the median eminence (Han et al., 2005; Herbison, 2016; Sisk and Foster, 2004). GnRH signaling in the pituitary gland facilitates the secretion of LH (luteinizing hormone) and FSH (follicle stimulating hormone), which stimulates gonadal cells in the testes and ovaries to initiate secretion of testosterone and estrogen. Reduction in negative-feedback regulation of GnRH neurons via steroids during puberty further increases the activity of GnRH neurons, ultimately leading to a gradual increase in circulating sex steroids in the periphery and the brain (Herbison, 2016; Sisk and Foster, 2004). The increase in circulating sex hormones during puberty is essential for physical, behavioral and mental maturation; however, a detailed molecular understanding of these processes remains elusive.

Extensive investigations into neuronal circuitry underlying reproductive behaviors have consistently shown that the medial preoptic area (MPOA) in the anterior hypothalamus, is a conserved circuit node in both males and females (Clemens and Yang, 2000; Fang et al., 2018; Heimer and Larsson, 1967; Karigo et al., 2020; Kohl et al., 2018; McHenry et al., 2017; Oomura et al., 1988; Rosenblatt et al., 1996; Rosenblatt and Ceus, 1998; Wei et al., 2021, 2018; Wu et al., 2014). Molecularly defined subpopulations of neurons expressing a variety of neuropeptides and/or hormonal receptors in the MPOA are tightly associated with reproductive behaviors. MPOA neurons expressing *Gal* (galanin) or *Esr1* (estrogen receptor 1) are essential for parental behaviors, while MPOA neurons expressing *Esr1* or *Nts* (neurotensin) govern male-type mating behaviors and female socio-sexual behaviors, respectively (Fang et al., 2018; Karigo et al., 2020; Kohl et al., 2018; McHenry et al., 2017; Wei et al., 2018; Wu et al., 2014). Since the MPOA is enriched in the expression of steroid hormone receptors genes (e.g., *Esr1*, androgen receptor (*Ar*), progesterone receptor (*Pgr*)) (Brock et al., 2015; Moffitt et al., 2018), an increase in the levels of sex hormones during puberty may support the maturation of MPOA neural circuits needed for reproductive and other behaviors.

Differential effects of circulating steroids between sexes e.g., estrogen in females and testosterone in males, potentially underly sexually dimorphic features of the MPOA. Interestingly, whereas some sexually dimorphic features, including cell density and connectivity, are determined predominantly by the presence or absence of the perinatal testosterone surge (organizational effects), the maintenance of neuronal circuitry mediating sexually dimorphic reproductive behaviors requires intact levels of circulating sex hormones during adulthood (activational effects) (Gegenhuber and Tollkuhn, 2020; Hull and Dominguez, 2019; McCarthy et al., 2017; Nugent et al., 2015; Wu et al., 2009; Xu et al., 2012). As the levels of circulating sex steroids start to rise and incrementally approach those of adults during puberty, hormone-dependent alterations of neuronal circuitry gradually mature, as observed in cortical and subcortical areas (Afroz et al., 2016; Piekarski et al., 2017; Sigl-Glöckner et al., 2019). Since the past low-throughput studies of hormone and behaviors biasedly targeted single genes without single cell resolution, the comprehensive understanding of molecular underpinnings of brain pubertal maturation are still elusive. Recent advancement of scRNAseq technology allowed characterizations of transcriptionally defined cell-types in sexually dimorphic nuclei (Chen et al., 2019; Hrvatin et al., 2020; Kim et al., 2019; Moffitt et al., 2018; Welch et al., 2019; Wu et al., 2017). However, identification of pubertal transcriptional dynamics and their regulatory mechanisms at single cell resolution has not been achieved in the brain of any species.

We conducted scRNAseq of 58,921 MPOA cells at several developmental time points (pre-, mid, and post puberty) and in distinct hormonal states (intact vs. gonadectomized) in male and female mice. Our results indicate that hormonal regulation of transcriptional dynamics is predominantly observed in a subset of cell types expressing steroid hormone receptors genes, among which cell-types expressing *Slc32a1* and *Esr1* (Vgat^+^ Esr1^+^) are transcriptionally relevant populations for male and female sexual behaviors. Transcriptional trajectory analysis of scRNAseq and highly multiplex FISH data both revealed the emergence of an adult-like transcriptional state during puberty, which was bidirectionally delayed or facilitated by manipulating levels of sex hormones. Gene regulatory network analysis in conjunction with cell type specific deletion of *Esr1* in MPOA Vgat^+^ cells demonstrates that Esr1 regulates pubertal transcriptional dynamics via sexually monomorphic and dimorphic transcription factor (TF) networks and also governs the maturation of sexual behaviors. These data reveal the unique pubertal transcriptional trajectories that exist between sexes and cell types and will serve as an important resource for the continued study of cell type specific gene expression during adolescence brain development.

## Results

### Identification of transcriptionally dynamic cell types during puberty in the MPOA

To derive pubertal transcriptional dynamics in the MPOA, we collected MPOA tissues at pre- (P23), mid- (P35) and post- (P50) puberty timepoints of both sexes, as well as hormonally deprived mice during the entire puberty period (being gonadectomized at P23 and tissue harvested at P50) to utilize droplet-based scRNAseq (Chromium V2, 10x genomics) (**Figure 1A**) (Hashikawa et al., 2020; Macosko et al., 2015; Rossi et al., 2019). After performing quality control and doublet removal (see methods), we recovered the transcriptomes of 58,921 cells (In total 20,345 genes, median unique molecular identifiers (UMIs)/cell: 1,893, median genes/cell: 1,054, **Figure S1**), which were comparable to previously reported scRNAseq datasets targeting anterior hypothalamus (Hrvatin et al., 2020; Moffitt et al., 2018). To jointly cluster all the cells in the 8 experimental groups while minimizing the effects of experimental variations and transcriptional alternations by sex, age, and hormonal states on clustering, we performed Seurat V3 integrative analysis, which implemented canonical correlation analysis (CCA) and mutual nearest neighbors analysis (MNN) to compute a transformation matrix for batch correction between datasets (Butler et al., 2018; Haghverdi et al., 2018; Stuart et al., 2019). After Seurat integration, followed by dimensionality reduction using principal component analysis and graph-based clustering, we identified 32 clusters of transcriptionally distinct cell types. Differentially expressed gene (DEG) analysis identified 1,430 marker genes in total for the identified clusters. Thirteen neuronal clusters were defined by the expression of *Stmn2* and/or *Thy1*, while the remaining 19 clusters were enriched with distinct non-neuronal cell markers (**Figure S1**). To gain a high-resolution census of neuronal cell types and their pubertal transcriptional dynamics, we re-clustered 24,627 neuronal cells identified from the initial clustering. Thirty-five neuron specific clusters were identified with 434 marker genes in total (UMIs/cell: 5,395, median genes/cell: 2,592, **Figure 1B, C** **and S1**). Consistent with previous observations, 20 predominantly GABAergic clusters (Vgat^+^ clusters) were comprised of 13,334 cells, while 7,658 neurons contributed to 12 predominantly glutamatergic clusters (Vglut2^+^ clusters). An additional 3 clusters composed of 3,635 neurons expressed a mixture of glutamatergic and GABAergic markers. Importantly, all the neuronal cell types were represented with comparable proportions of cells from all the experimental groups and also expressed conserved marker genes, irrespective of their age, sex, and hormonal states (**Figure 1B, C** **and S1H**). These results suggest that terminal diversification and organization of neuronal cell types in the MPOA are largely completed prior to puberty, which is in agreement with a previous microarray or scRNAseq studies in the entire hypothalamus (Kim et al., 2020; Romanov et al., 2020; Shimogori et al., 2010).

**Figure 1.**
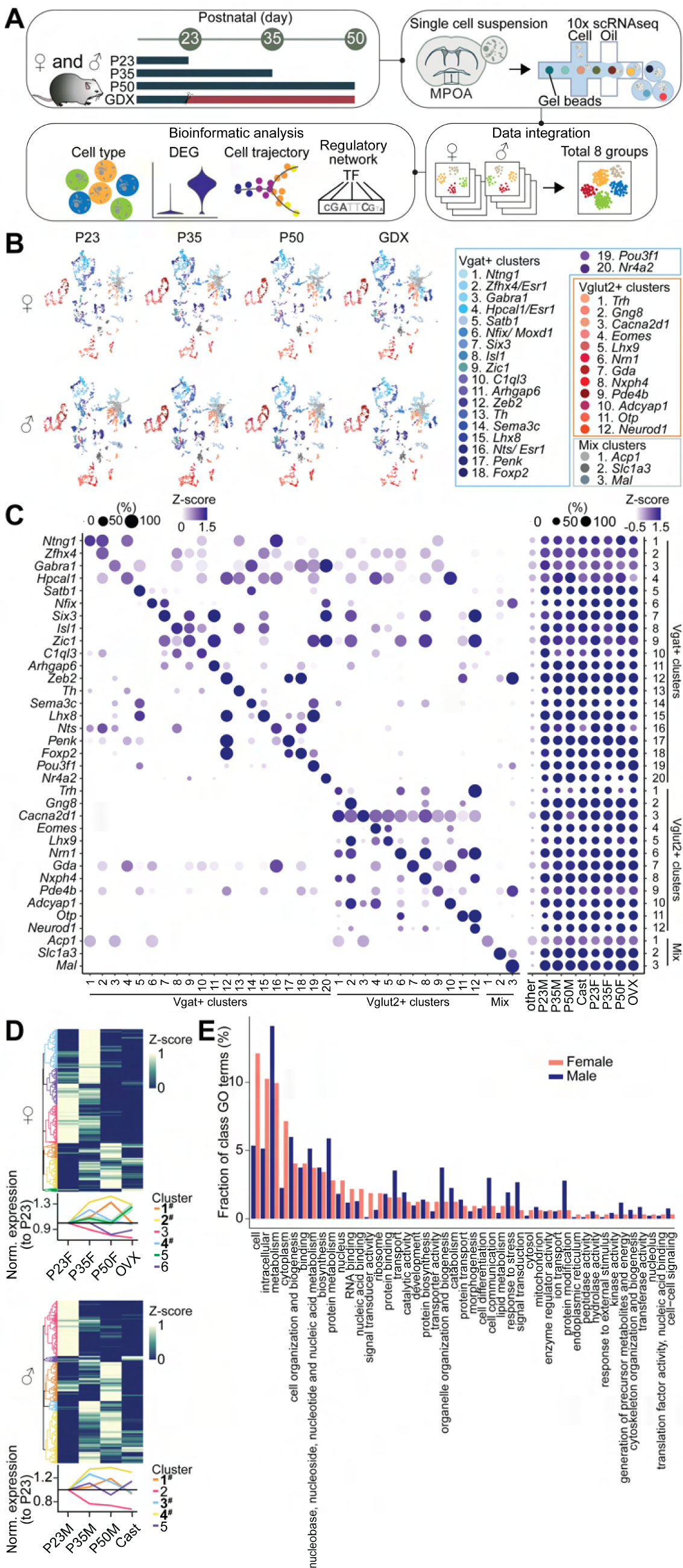
scRNAseq to jointly identify neuronal cell-types across puberty in the MPOA. (A) Schematic illustrating scRNAseq experiments. (B) Joint clustering of neuronal cells from all the groups identified 35 clusters. UMAP plot of each experimental group is color-coded by cell-type. (C) Dot plots illustrating scaled expression levels (color) and the proportions of expressing cells (dot size) of marker genes in each cluster from all the groups (left) or in each group (right). (D) Top: heatmaps showing hierarchical clustering of DEGs in all Vgat^+^ population in females (top) and males (bottom). Bottom: normalized expression of each gene cluster relative to the level at P23. Boldface clusters (#) show higher expression in P35 and P50 than P23 and GDX. One-way repeated measures ANOVA followed by multiple comparisons. Detailed stats are in Methods. (E) Enriched gene ontology terms in HA-DEGs in aggregated Vgat^+^ populations were categorized into GO classes. Fraction of each GO class term in female (salmon) and male (blue) were shown. Shades represent S.E.M. Detailed statistics are in Methods.

To examine transcriptional alterations associated with puberty, we conducted pseudo-bulk DEG analysis between all possible pairs of experimental groups within the same sex for aggregated Vgat^+^ and Vglut2^+^ cells. These pairwise comparisons identified DEGs in Vglut2+ and Vgat^+^ MPOA cells (Vglut2^+^ males:1626, female: 520; Vgat^+^ male:3201, female: 1501) (**Table S1**). To determine whether there are groups of DEGs enriched in particular combinations of experimental groups, we conducted hierarchical clustering on the expression of the DEGs across experimental groups in aggregated Vgat^+^ or Vglut2^+^ cells. In Vgat^+^ and Vglut2^+^ cells of both sexes, we found groups of genes selectively enriched in prepuberty (P23) samples. Additionally, groups of genes were identified where the highest level of expression was observed in hormonally intact adult (P50) and/or puberty (P35) samples with lower expression in hormonal deprivation and prepuberty samples (**Figure 1D** **and S1I; Table S1**). Gene ontology analysis on these hormonally associated DEGs (HA-DEGs) identified a number of GO terms (female:144, male:405) (Chen et al., 2013; Kuleshov et al., 2016). Further GO term categorization (Reecy JM, 2008) revealed that a large proportion of GO categories were associated with cell metabolism, gene expression and regulation, and processing of proteins while GO categories directly related to functional and electrophysiological properties of neurons (e.g. ion transport, morphogenesis) were also identified in both sexes (**Figure 1E**).

A large proportion (Vgat^+^: 51.7-62.2%; Vglut2^+^: 20.6-32.0%) of hormonally associated puberty/adult enriched genes identified in pseudo-bulk analysis (**Figure 1D, E** **and S1I; Table S1**) implies that circulating steroids regulate pubertal transcriptional states in a subset of cell types enriched with hormonal receptors in the MPOA. We next computed HA-DEGs for each cell type by comparing intact adult (P50) or puberty (P35) with gonadectomized scRNAseq data. The number of HA-DEGs was highly variable across cell types (0 - 88 HA-DEGs/cell type) (**Figure 2A, B** **and S1J; Table S2**). We next examined whether hormonal receptor gene expression correlated with the proportion of HA-DEGs in each neuronal cell type. Regression analysis between the fraction of hormone-receptor expressing cells and the number of HA-DEGs revealed that the abundance of *Esr1* and/or *Pgr* in specific cell types predicted the degree of HA-DEGs (**Figure 2A-D**, **S1J, K**). Collectively, although MPOA neuronal cell types are largely diversified by prepuberty (P23, **Figure 1B, C** **and S1H**), a subset of cell types defined by hormonal receptors (e.g., *Esr1*) in Vgat^+^ and Vglut2^+^ cells of both sexes display hormonally dependent, transcriptional refinement through puberty to adulthood (**Figure 2A-D** **and S1J, K**).

**Figure. 2.**
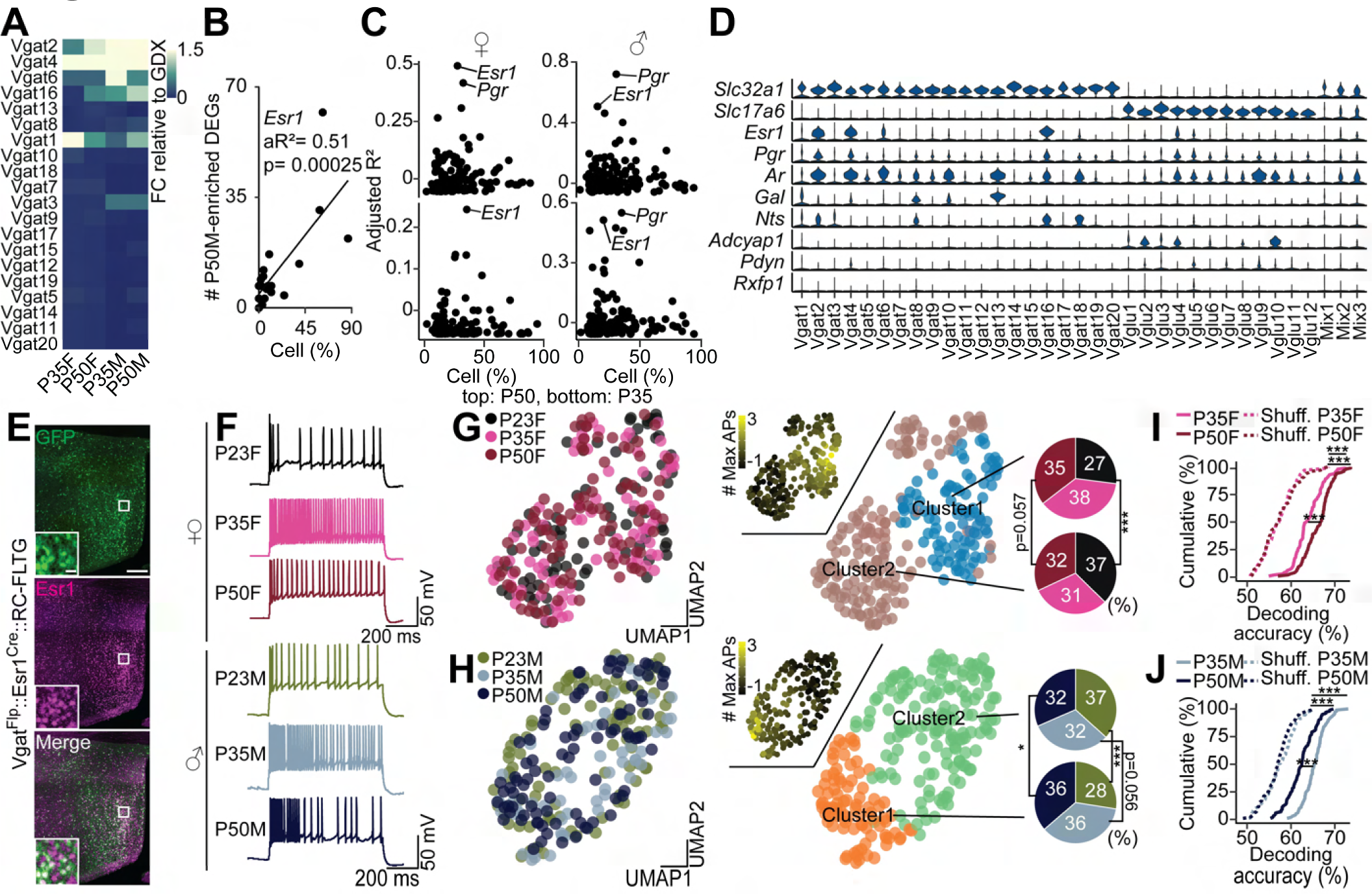
MPOA^Vgat+ Esr1+^ neurons are transcriptionally and electrophysiologically dynamic during puberty. (A) A heatmap showing scaled sum of logFC for each Vgat^+^ cluster in comparison with GDX samples. (B) Linear regression analysis between the percentage of Esr1 expressing cells and the number of P50M-enriched DEGs in comparison with GDX samples at each Vgat^+^ cluster (each dot). Linear regression analysis. aR^2^: adjusted R squared. (C) Scatter plots showing adjusted R^2^ values of hormone receptor genes (dots) in females (left) and males (right) as a result of linear regression analysis in comparison between P50 (top) or P35 (bottom) and GDX. (D) Violin plots showing the expressions of Slc32a1, Slc17a6, several steroid hormone receptor genes and canonical marker genes in the MPOA at each Vgat^+^ cluster. (E) Representative images showing GFP (left), Esr1 (middle) and merge (right) around the MPOA of a Vgat^Flp^::Esr1^Cre^::RC-FLTG mouse. (F) Representative whole-cell recordings following 220 pA current injections from cells at P23, P35 and P50 in females (left) and males (right). (G, H) UMAPs color coded by groups (left), # max firing (middle) and clusters (right) in females (G) and males (H). Pie charts showing the fractions of groups in each cluster. Fisher’s exact test. *p < 0.05, ***p < 0.001. (I, J) Cumulative distributions of decoding accuracy between P50 (female: dark red, male: dark blue) or P35 (female: pink, male: light blue) and P23 in females (I) and males (J). One-way ANOVA followed by multiple comparisons. Details in Methods. Error bars represent S.E.M. Detailed statistics are in Methods.

### Identification of transcriptionally defined cell types relevant for reproductive behaviors

The MPOA is comprised of heterogeneous cell types mediating various functions and behaviors, ranging from thermoregulation, regulation of endocrine system, social behaviors, hunting behaviors, and other motivated behaviors (Fang et al., 2018; Hrvatin et al., 2020; Kohl et al., 2018; McHenry et al., 2017; Park et al., 2018; Robison et al., 2018; Tan et al., 2016; Tobiansky et al., 2016, 2013; Wu et al., 2014)(**Figure 1B, C**). However, it is unclear which transcriptionally defined cell types in our dataset are relevant for reproductive behaviors, and whether those behaviorally relevant neural populations are transcriptionally dynamic during puberty. To address this, we utilized a publicly available spatial transcriptomic dataset of POA cells (acquired using the MERFISH platform)(Moffitt et al., 2018) and jointly analyzed the correspondence between MERFISH data and our scRNAseq data. First, we clustered neuronal cells in the MERFISH data and computed the enrichment of behaviorally induced neuronal activity marker (*Fos*) in MERFISH derived clusters. Consistent with previous observations, only a subset of *Gad1*^+^ inhibitory MERFISH clusters were enriched with *Fos* induced by social behaviors (mating, parenting and fighting behaviors) (**Figure S2A**). Second, to establish the correspondence between MERFISH clusters and our scRNAseq clusters, we utilized Seurat V3 integrative analysis to align cell-type specific MERFISH data associated with social behaviors onto our scRNAseq data (reference-based integration) (**Figure S2B**) (Stuart et al., 2019). In females, mating-relevant MERFISH clusters (mGad1-3 and mGad1-7) were related to a subset of scRNAseq clusters while parental-behavior related mGad1-9 and mGad1-10 corresponded to partially distinctive clusters **Figure S2C**). In males, mating-related mGad1-3 and mGad1-10 and aggression-related mGad1-9 respectively corresponded to a subset of scRNAseq clusters (**Figure S2C**). Next, we computed marker genes in the mating-relevant scRNAseq clusters. In both the male and female dataset, *Esr1* was the most enriched and selective gene in the Vgat clusters related to mating (**Figure S2D-E; Table S3**). This observation is consistent with a previous study demonstrating that optogenetic stimulation of Vgat^+^ Esr1^+^ neurons in the MPOA is sufficient to induce sexual behaviors in male mice (Karigo et al., 2020). Overall, a subset of Vgat^+^ clusters expressing *Esr1*, which are transcriptionally dynamic during puberty, are also relevant to male and female mating behaviors.

### Enhanced neuronal excitability of Vgat^+^ Esr1^+^ neurons during puberty

To examine the pubertal maturation of the electrophysiological properties of Vgat^+^ Esr1^+^ neurons, we conducted *ex vivo* slice electrophysiological experiments to characterize cellular excitability of Vgat^+^ Esr1^+^ neurons at pre-, during and post-puberty periods. To selectively mark Vgat^+^ Esr1^+^ cells, we bred Vgat^Flp^::Esr1^Cre^::RC-FLTG mice(Lee et al., 2014, p. 1; Plummer et al., 2015) where only Vgat^+^ and Esr1^+^ neurons were labeled with GFP (specificity: 91.3%. **Figure 2E**). We measured the intrinsic excitability of Vgat^+^ Esr1^+^ cells at prepuberty (P23), mid-puberty (P35) and post puberty (P50) time points using whole-cell, patch-clamp electrophysiological recordings (**Figure 2E-I**). In both males and females, current-evoked action potential firing rates were the highest in mid puberty groups followed by that of adult and prepuberty groups (**Figure 2F** **and Figure S2H**). K-means clustering using 14-20 distinct electrophysiological features separated recorded neurons into 2 clusters, one of which displayed higher excitability in both sexes (**Figure 2G, H** **and Figure S2F, G**). The proportions of P23 cells of both sexes in the low-firing clusters were smaller than those in high-firing clusters whereas those differences were not observed and opposite trends were present in P35 and adult groups (**Figure 2G, H**). Consistently, support vector machine (SVM) classification revealed that P23 cells were distinguishable from the other cell groups by their combination of electrophysiological features (**Figure 2I, J**). Collectively, although functionally heterogeneous, a subpopulation of Vgat^+^ Esr1^+^ neurons display enhanced neuronal excitability through puberty to adulthood.

### Pubertal transcriptional trajectories in Vgat**^+^** Esr1**^+^** neurons

During puberty, MPOA neurons transition from a juvenile to an adult transcriptional state as indexed by the expression of single genes (Blakemore et al., 2010; Perrin et al., 2008)(**Figure 1 and 2**). Although DEG analysis identified genes which were statistically enriched in aggregated cell-types (**Figure 1** **and** **Figure 2**), it did not provide how transcriptional states of single cells transition between distinct biological states. Pseudotime (trajectory) analysis has been widely utilized to compute transition of transcriptional states of single cells in various biological processes including development and disease (Cao et al., 2019; Morabito et al., 2021; Qiu et al., 2017; Rossi et al., 2019). To elucidate the cell-type specific progression of transcriptional dynamics of single cells during puberty, we performed pseudotime analysis, which measures transcriptional progression of single cells through a biological process (puberty) by learning a principal graph from the combinatorial gene expression in single cells in UMAP space (Cao et al., 2019; Qiu et al., 2017). In Vgat^+^ Esr1^+^ cells for males and females, transcriptional states of P35 and P50 groups were the most distinct from the prepuberty group within the transcriptional trajectory (**Figure 3A, E**). Strikingly, neurons from hormonally deprived groups showed arrested transcriptional progression with a high degree of similarity to cells in the prepuberty state (**Figure 3A, B, C, E, F, G**). This transcriptional trajectory pattern was also observed at the level of individual Vgat^+^ Esr1^+^ clusters (**Figure S3E, F**). Hormone-dependent transcriptional trajectories in Vgat^+^ cells that do not express steroid hormone receptor genes (Vgat^+^ hormone R^Low^) (**Figure S3A**), were minimal (**Figure 3D, H** **and S3B, C**), consistent with the idea that these cells are less transcriptionally dynamic during puberty.

**Figure 3.**
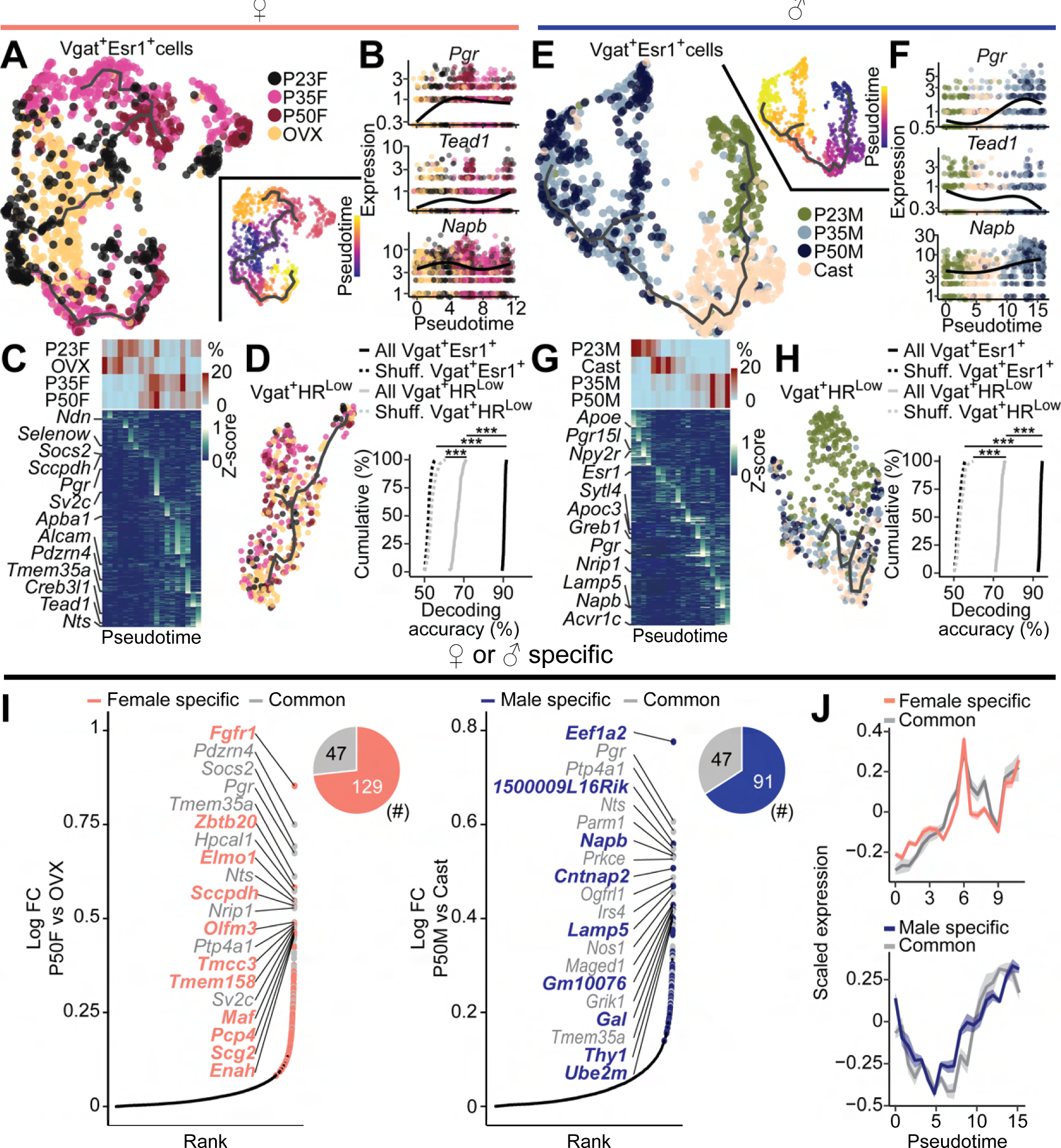
Identification of pubertal transcriptional trajectories in MPOA^Vgat+ Esr1+^ neurons. (A, E) UMAP visualization of transcriptional trajectory (black line) and cells (dots) color-coded by group (left) or pseudotime (right) in female (A) and male (E) Vgat^+^ Esr1^+^ populations. (B, F) Kinetics plots showing relative expression of hormonally associated DEGs in females (*Tead1*), males (*Napb*) or both (*Pgr*) across pubertal pseudotime in female (B) and male (F) Vgat^+^ Esr1^+^ population. (C, G) Heatmaps showing proportion of cells (top) and scaled expression of DEGs (bottom) across pseudotime in females (C) and males (G). (D, H) (left) UMAP visualization of transcriptional trajectory (black line) and cells (dots) color-coded by group in female (D) and male (H) Vgat^+^ HR^Low^ populations. (right) Cumulative distributions of decoding accuracy by SVM classification between mature groups (P50, P35) and immature group (P23, GDX) using expression data from Vgat^+^ Esr1^+^ (black, real line), Vgat^+^ HR^Low^ (grey) or shuffled data (dashed line) in females (D) and males (H). One-way ANOVA followed by multiple comparisons. Details in Methods. (I) Ranked logFC values of all genes in comparison between P50 and GDX in females (left) and males (right). Top 20 genes are highlighted, color-coded by being sex-specific or sex-common and their numbers are shown in pi-charts. (J) Averaged, scaled expression of HA-DEGs, which are sex-specific (female: salmon, top; male: blue, bottom) or sex-common (grey). Shades represent S.E.M. ***p < 0.001. Detailed statistics are in Methods.

Our DEG and pseudotime analyses together suggest that transcriptional states, which are defined by combinatorial expression of HA-DEGs (**Table S4**), transition from immature (prepubertal) state to mature (pubertal and adult) state during puberty. This assumption predicts that fate of single cells can be decoded solely from the gene expression in scRNAseq data (Weinreb et al., 2020). In support of this assumption, HA-DEGs in transcriptional trajectories were sufficient to decode via a support vector machine classifier (SVM) whether single Vgat^+^ Esr1^+^ cells belonged to transcriptionally mature groups (intact adult or puberty) or immature groups (prepuberty or hormonal deprivation) with high accuracy (median prediction accuracy 90.6 – 93.6 %) whereas decoding accuracy for Vgat^+^ hormone R^Low^ cells was much lower (median 66.7 - 73.0 %) (**Figure 3D, H**). Since all Vgat^+^ Esr1^+^ clusters co-expressed *Ar* to a high degree (68.3 – 88.3 %), we tested whether Vgat^+^ clusters that expressed *Ar*, but not *Esr1* (Vgat^+^ Esr1^-^/ AR^+^) (**Figure S3A**), showed hormonally dependent transcriptional progression. Consistent with higher levels of circulating testosterone in males, a subset of P35 and P50 cells of male Vgat^+^ Esr1^-^/ AR^+^ showed moderately progressive transcriptional states in comparison to prepuberty and hormonally deprived groups whereas this pattern was diminished in female Vgat^+^ Esr1^-^/ AR^+^(**Figure S3B, C**). Indeed, the decoding accuracy for male Vgat^+^ Esr1^-^/ AR^+^ was higher compared to females (**Figure S3D**). Importantly, to further computationally validate our trajectory analysis, which utilized Monocle V3, we conducted Manifold Enhancement Latent Dimension (MELD) analysis, which could measure continuous transcriptional progression by experimental perturbation as a likelihood of observing particular experimental conditions at single cells (Burkhardt et al., 2021; Moon et al., 2019). Consistent with our Monocle pseudotime analysis, MELD analysis revealed that intact adult and puberty groups of both sexes were spatially segregated and had higher likelihood of matured transcriptional states than prepuberty and hormone-deprivation groups in Potential of Heat-diffusion for Affinity-based Trajectory Embedding (PHATE) space (**Figure S3G, H**). Collectively, in both sexes, pubertal MPOA transcriptional progression of single cells was most prominently observed in the Vgat^+^ Esr1^+^ population, where the transcriptional state continuously transitioned from prepuberty to adulthood and required steroid synthesis from the gonads.

Although a clear transcriptional trajectory was observed in Vgat^+^ Esr1^+^ of both sexes, shared and distinct groups of genes were altered along the trajectory (**Figure 3I, J**; **Table S4**). Some genes (47) were highly expressed in intact adult (P50), compared to gonadectomized groups in both sexes. In contrast, 91-129 genes were significantly enriched only in one sex of intact adults e.g., *Tead1* in female, *Napb* in male. These shared and sex-specific DEGs showed similar progression patterns along the pseudotime transcriptional trajectory (**Figure 3J**). Thus, whereas continuous transcriptional progression towards the adult state is observed in both sexes in Vgat^+^ Esr1^+^ cells, sexually monomorphic and dimorphic gene expressions emerge within the pubertal transcriptional trajectory.

### Hormonal control of pubertal transcriptional trajectories

Delayed or immature transcriptional trajectories in gonadectomized mice (P50 at the time of harvest) (**Figure 3**) suggests that aging is not the sole determinant of pubertal transcriptional alterations, but that circulating hormones are critical in this process. To cross-validate our trajectory analysis from the scRNAseq data as well as to examine its hormonal regulation, we performed highly multiplexed hybridization chain reaction FISH (HM-HCR FISH, V3), which we used to detect transcripts of ∼12 genes in 41,549 cells at single molecule resolution *in situ* (**Figure 4A-C** **and S4**) (Choi et al., 2018; von Buchholtz et al., 2021) since the transcriptional states were determined by the combination of gene expressions. Besides the experimental groups used in scRNAseq experiments, we also prepared intact P28 male and female mice and hormonally supplemented groups receiving testosterone (male) or estrogen (female) from P23-P27 and harvested tissue at P28 to examine whether pubertal transcriptional trajectories were facilitated by early supplementation of circulating steroids. Our hormonal supplement protocol induced early puberty onset in 100% of tested mice including the subjects used in HM-HCR FISH experiments (Puberty onset age: Balanopreputial Separation (BPS) in males and Vaginal Opening (VO) in females). For intact males: puberty onset was P31.75 ± 1.03, hormonally supplemented males: P26.86 ± 0.14, intact females: P31.67 ± 1.20, hormonally supplemented females: P26 ± 0.37. To distinguish Vgat^+^ Esr1^+^ cells from Vgat^+^ hormone R^Low^, we measured *Slc32a1*, *Esr1* and *Ar* along with 8-9 HA-DEGs during puberty (**Figure 4D-E** **and S4**). Consistent with trajectory analysis using scRNAseq data, pseudotime analysis of HM-HCR data revealed that the transcriptional state of single cells associated with puberty (P35) and the intact adult (P50) groups were distinct from that of prepuberty (P23) and gonadectomized groups in Vgat^+^ Esr1^+^ cells (**Figure 4F-G**). In addition, hormone-supplemented groups of both sexes (P28) exhibited comparable transcriptional progression with the puberty and the adult groups, while age-matched control groups (P28) showed an immature transcriptional state similar to prepuberty groups (**Figure 4F-H** **and K**). To determine whether combinatorial DEG expressions in HM-HCR data are deterministic of cell fate, we conducted decoding analysis. SVM classification revealed that 10-11 genes in HM-HCR FISH data of Vgat^+^ Esr1^+^ cells were sufficient to decode whether single cells were from transcriptionally mature or immature groups with high accuracy (84.4 – 85.5 %) while the same set of genes underperformed in decoding for Vgat^+^ hormone R^Low^ cells (63.6 – 64.6 %) (**Figure 4I-L**). In addition, decoding capability was reduced by using a subset of genes (**Figure 4J-M**), further indicating that transcriptional states were defined by the combination of DEG expressions. In order to directly cross-reference multimodal scRNAseq and HM-HCR FISH datasets, we utilized Seurat V3 to find anchors between HM-HCR FISH and scRNAseq data and to impute gene expression profiles of scRNAseq by transferring the expression of HM-HCR FISH (**Figure 4N**) (Stuart et al., 2019). 90.9-100 % of imputed genes showed statistically significant correlation with real expression data (**Figure 4O, R**). Moreover, imputed scRNAseq datasets recapitulated pseudotime analysis and SVM classification results (**Figure 4P, Q, S, T**). These results strengthened and corroborated the pseudotime analysis of scRNAseq data by visualizing pubertal transcriptional trajectories in space, demonstrating its bidirectional control by circulating sex hormones, and the successful integrative analysis between the two distinct modalities.

**Figure 4.**
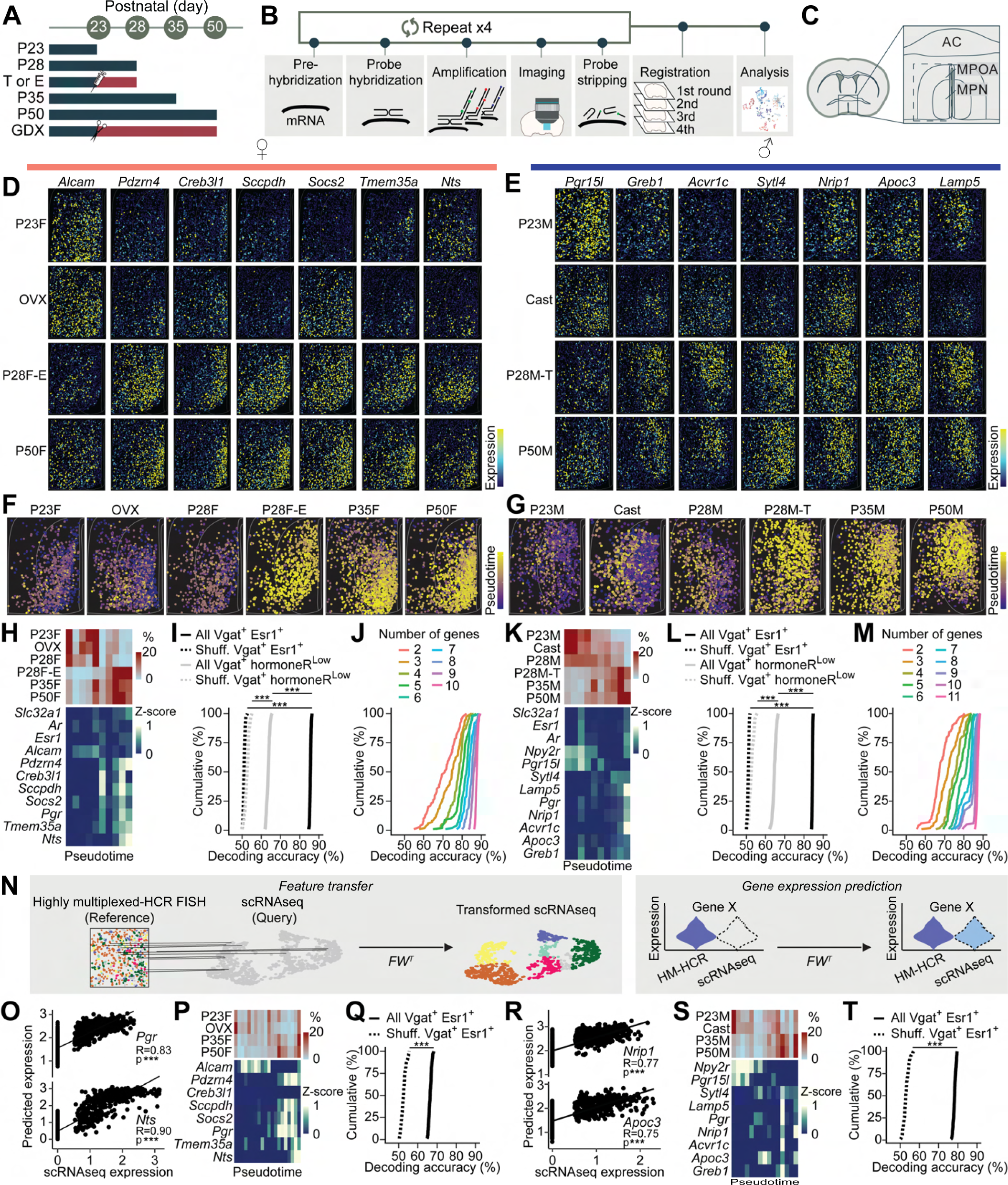
HM-HCR FISH to visualize pubertal transcriptional trajectories of MPOA^Vgat+ Esr1+^ neurons in space and their regulation by circulating sex hormones. (A) Experimental groups. (B) Schematic illustrating highly multiplexed HCR FISH procedure. (C) 41,549 cells from MPOA in total were analyzed. (D, E) Representative images of reconstructed cells, color-coded by scaled expression of genes in the MPOA from P23, GDX, P28 (hormone supplemented) and P50 in females (D) and males (E). (F, G) Pseudotime of Vgat^+^ Esr1^+^ population visualized in space across groups in females (F) and males (G). (H, K) Heatmaps showing proportion of cells (top) and scaled gene expression (bottom) across pseudotime in females (H) and males (K). (I, L) Cumulative distributions of decoding accuracy between mature groups (P50, P35, hormone-supplemented P28) and immature groups (P23, GDX, P28 control) using expression data from Vgat^+^ Esr1^+^ (black, real line), Vgat^+^ hormoneR^Low^ (grey) or shuffled data (dashed line) in females (I) and males (L). One-way ANOVA followed by multiple comparisons. Details in Methods. (J, M) Cumulative distributions of decoding accuracy between mature and immature groups using subsets of expression data from Vgat^+^ Esr1^+^, color-coded by the number of genes used for decoding in females (J) and males (M). (N) Schematic illustrating integrative analysis between HM-HCR FISH and scRNAseq to predict scRNAseq dataset. (O, R) Scatter plots showing the correlated expressions of real and predicted scRNAseq data in females (O) and males (R). R: Pearson correlation coefficient. (P, S) Heatmaps showing proportion of cells (top) and scaled predicted gene expression (bottom) across pseudotime in females (P) and males (S). (Q, T) Cumulative distributions of decoding accuracy between mature groups (P50, P35) and immature groups (P23, GDX) using predicted expression data in females (Q) and males (T). unpaired t-test. ***p < 0.001. Detailed stats are in Methods.

### Gene regulatory mechanisms for pubertal transcriptional dynamics

Hormone-dependent transcriptional dynamics during puberty suggests the possibility of unique regulatory mechanisms in males and females. To identify a gene regulatory network, first we utilized single-cell regulatory network inference and clustering (SCENIC) analysis to rank TFs by their co-expression values with HA-DEGs (Aibar et al., 2017). In both females and males, *Esr1* had the highest co-expression scores among 915-952 transcription factors (**Figure 5A**), implying that *Esr1* directly or indirectly regulates HA-DEGs. To test for causal roles of *Esr1*, we developed a viral based cell-type and region-specific deletion of *Esr1* because previous global Esr1 deletion/knockdown technologies lacked cell-type specificity, region specificity or temporal specificity (affecting the entire lifetime) (Ogawa et al., 1998, 1997; Ribeiro et al., 2012; Sano et al., 2013). Double transgenic mice (Vgat^Flp^::Esr1^lox/lox^) were injected with AAV-CAG-frtFlex-Cre:GFP (thereafter AAV-frtFlex-Cre) (Heymann et al., 2020) into the MPOA at P14-18 to delete *Esr1* during the entire puberty period (Hewitt et al., 2010). MPOA from these mice were then collected at P50 for subsequent scRNAseq (**Figure 5B**). Immunohistochemistry and FISH validated gene deletion efficiency and specificity of the virus (**Figure S5A, B and** **Figure 7B, J**). Many genes were downregulated by Esr1KO in Vgat^+^ Esr1^+^ neurons; 600 and 824 in males or females, respectively (Esr1-DEGs), while only 5.3 % and 3.3 % of Esr1-DEGs were reduced in Vgat^+^ hormone R^Low^ cells (**Figure 5C**). HA-DEGs overlapped with Esr1-DEGs suggesting that *Esr1* deletion recapitulates the effects of hormone deprivation on the transcriptomes of Vgat^+^ Esr1^+^ cells (males: 56.5%, 78 genes; females: 73.3 %, 124 genes; **Figure 5C**). Pseudotime analysis also indicated that Esr1KO near-completely prevented pubertal transcriptional maturation (**Figure 5D, E**). Thus, Esr1 plays the indispensable roles in the regulation of hormone dependent transcriptional dynamics during puberty.

**Figure 5.**
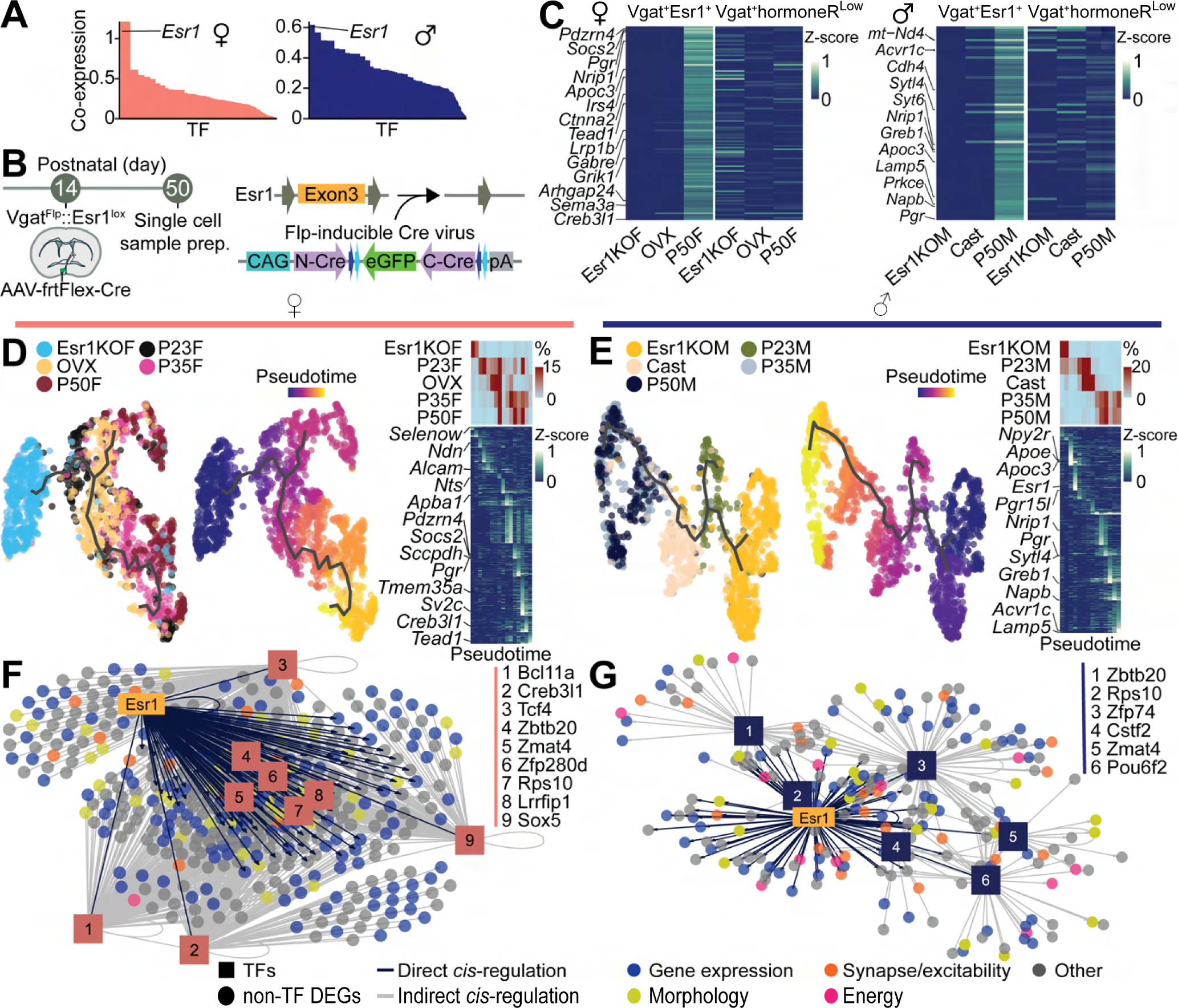
Deconstructing gene regulatory networks underlying pubertal transcriptional dynamics in MPOA^Vgat+ Esr1+^ neurons. (A) SCENIC analysis computed ranked sum of co-expression scores of TFs with HA-DEGs in females (left, 952 TFs) and males (right, 915 TFs). (B) Schematics illustrating cell-type, and site-specific deletion of Esr1 in the MPOA using AAV-frtFlex-Cre virus and Vgat^Flp^::Esr1^lox/lox^ mice. (C) Heatmaps showing scaled averaged expressions of HA-DEGs of Vgat^+^ Esr1^+^ or Vgat^+^ hormoneR^Low^ populations in Es1KO, GDX and P50 groups in females (left) or males (right). (D, E) UMAP visualization of transcriptional trajectory (black line) and cells (dots) color-coded by group (left) or pseudotime (middle) in female (D) and male (E) Vgat^+^ Esr1^+^ population. Right: Heatmaps showing proportion of cells (top) and scaled expressions of DEGs (bottom) across pseudotime. (F, G) Motif-enrichment analysis on Esr1-DEGs deconstructed GRNs where Esr1 and Esr1-regulated TFs (pink: females, dark blue: males) combinatorically cis-regulate Esr1-DEGs, color-coded by gene-categories identified through ontology analysis in females (F) and males (G).

**Figure 7.**
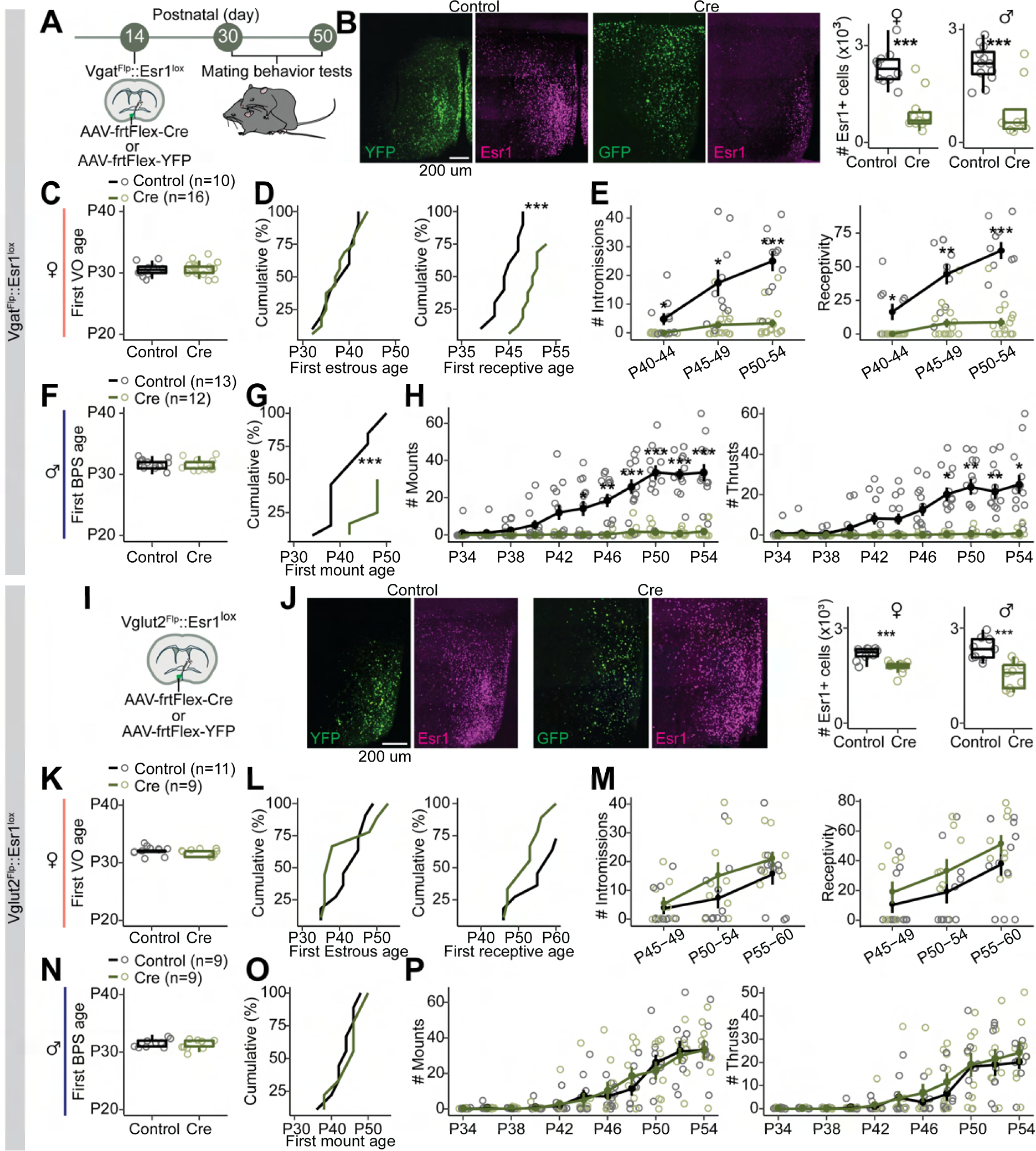
Esr1 in MPOA^Vgat+ Esr1+^ neurons governs pubertal maturation of sexual behaviors. (A) Schematics showing the timeline for behavioral experiments to test the role of Esr1 in MPOA^Vgat^ in pubertal maturation of sexual behaviors. (B) Left: representative images of viral-reporter or Esr1 expressions from control or Cre groups. Right: Quantification of Esr1 in the MPOA. Unpaired t-test. ***p < 0.001. (C-H) Quantitative comparisons of first VO age (C), estrous age (D, left), receptive age (D, right), BPS age (F) or mount age (G) and the number of intromissions (E, left), receptivity (E, right), the numbers of mounts (H, left) and thrusts (H, right). Line plots are shown in mean ± S.E.M. Box plots are shown with whiskers (min, max) and box (25%, median and 75%). C, D, F, G, Unpaired t-test. E, H, Two-way repeated measures ANOVA followed by multiple comparisons. ***p < 0.001. Statistical details in Methods. (I) Schematic showing the behavioral experiments to test the role of Esr1 in MPOA^Vglut2^ in pubertal maturation of sexual behaviors. (J) Left: representative images of viral-reporter or Esr1 expressions from control or Cre groups. Right: Quantification of Esr1 in the MPOA. Unpaired t-test. Details in Methods. (K-P) Quantitative comparisons of first VO age (K), estrous age (L, left), receptive age (L, right), the number of intromission (M, left), receptivity (M, right), BPS age (N) or mount age (O) and the numbers of mounts (P, left) and thrusts (P, right).

Upon estrogen binding, dimerized Esr1 can directly regulate its target genes by binding to their estrogen-response elements (ERE) (Carroll et al., 2006; Ikeda et al., 2015). To examine whether Esr1 cis-regulates Esr1-DEGs, we computed motif enrichments in Esr1-DEGs by utilizing RcisTarget analysis (Aibar et al., 2017). Fractions of Esr1-DEGs (123 (14.9 %, female) and 98 genes (16.33 %, male)) were significantly enriched with Esr1 motifs while 9 (female) or 6 (male) of these genes were TFs (causally regulated by Esr1. Esr1-TFs). We tested the possibility that Esr1-TFs further cis-regulate additional Esr1-DEGs. RcisTarget analysis inferred that 446 (54.126%, female) and 167 genes (28.3 %, male) of Esr1-DEGs were cis-regulated by Esr1-TFs. Collectively, 56.1 % (female) and 36.7 % (male) of Esr1-DEGs were potentially cis-regulated by Esr1 and/or Esr1-TFs. Based on these analyses, the gene regulatory network (GRN) for Esr1-DEGs was deconstructed (**Figure 5F, G**). Gene ontology analysis reveals that 4 classes of genes (related to gene expression and degradation, energy production, cell morphology, and synaptic transmission/cellular excitability) largely comprise Esr1-DEGs in both females and males (**Figure 5F, G** **and S5D**). Thus, the combination of Esr1 deletion, scRNAseq and motif-enrichment analysis revealed that GRN centering around Esr1 and Esr1-TFs (Esr1-GRN) orchestrates the expressions of functionally diverse genes to potentially support the maturation of MPOA functions in Vgat^+^ Esr1^+^ cells in both sexes.

### Development of transcriptional sexual dimorphism during puberty

Although GRNs in Vgat^+^ Esr1^+^ cells were centered around Esr1 in both sexes, 58.4% (female) and 42.8% (male) of Esr1-DEGs were sex specific. Consistently, 6 (female) or 3 (male) TFs in the identified GRN were unique in one sex (**Figure 5F, G**). Moreover, expression levels of most of Esr1-TFs increased during puberty and adulthood and they were reduced by gonadectomy or Esr1-KO (**Figure S6A, B**), suggesting that sexual dimorphism of cell type specific transcriptional states in the MPOA, at least in part, develops during puberty. Indeed, 71 out of 155 sexually dimorphic genes (comparison between P50 male and female; 45 female enriched genes, 110 male enriched genes) were in the Esr1-GRN (**Figure 6A** **and Table S5**). Next, we examined whether puberty-related Esr1-DEGs (intact P50 vs Esr1KO during puberty period) and sexually dimorphic genes (intact adult male vs female) were associated. Whereas a large proportion of sexually dimorphic genes were also in Esr1-DEGs in the dominant sex (34 out of 45 female dimorphic genes were Esr1-DEGs in females, 40 out of 110 male dimorphic genes are Esr1-DEGs in males), surprisingly, a subset of sexually dimorphic genes were also Esr1-DEGs in the opposite sex (13 out of 45 female dimorphic genes were Esr1-DEGs in males, 13 out of 110 male dimorphic genes are Esr1-DEGs in females) (**Figure 6B** **and Table S6**). These results suggest that a subset of Esr1-DEGs were commonly elevated during puberty in both sexes, but the degree of elevation was more pronounced in one sex. Interestingly, a subset of male dimorphic genes (15 out of 110) were elevated by Esr1KO in female, suggesting that Esr1 signaling in females suppresses several male dominant genes. To examine how sexual dimorphisms of the transcriptional states of single cells, which are defined by the combinatorial expression of sexually dimorphic Esr1-DEGs, develop during puberty, we performed trajectory analysis. We found that intact P50 females and males were transcriptionally the most distinct from each other while cells from P23 or gonadectomized male and female groups displayed a less dimorphic transcriptional state (**Figure 6C, D**, **and S6E, H**). To determine whether combinatorial expressions of sexually dimorphic Esr1-DEGs determined sex identities of single cells, we conducted decoding analysis. Dimorphic genes in the Esr1-GRN of Vgat^+^ Esr1^+^ cells were sufficient to decode the sexes of P50 groups with the highest accuracy (81.4 ± 0.11 %), followed by P35 and lowest in P23 groups, but not in Vgat^+^ hormone R^Low^ cells (**Figure 6E** **and S6F, G**). These results suggest that Esr1 orchestrates sex-specific transcriptional states via dimorphic sets of Esr1-TFs, which fully develops during puberty and may underlie the maturation of sexually dimorphic neuronal circuits in the MPOA.

**Figure 6.**
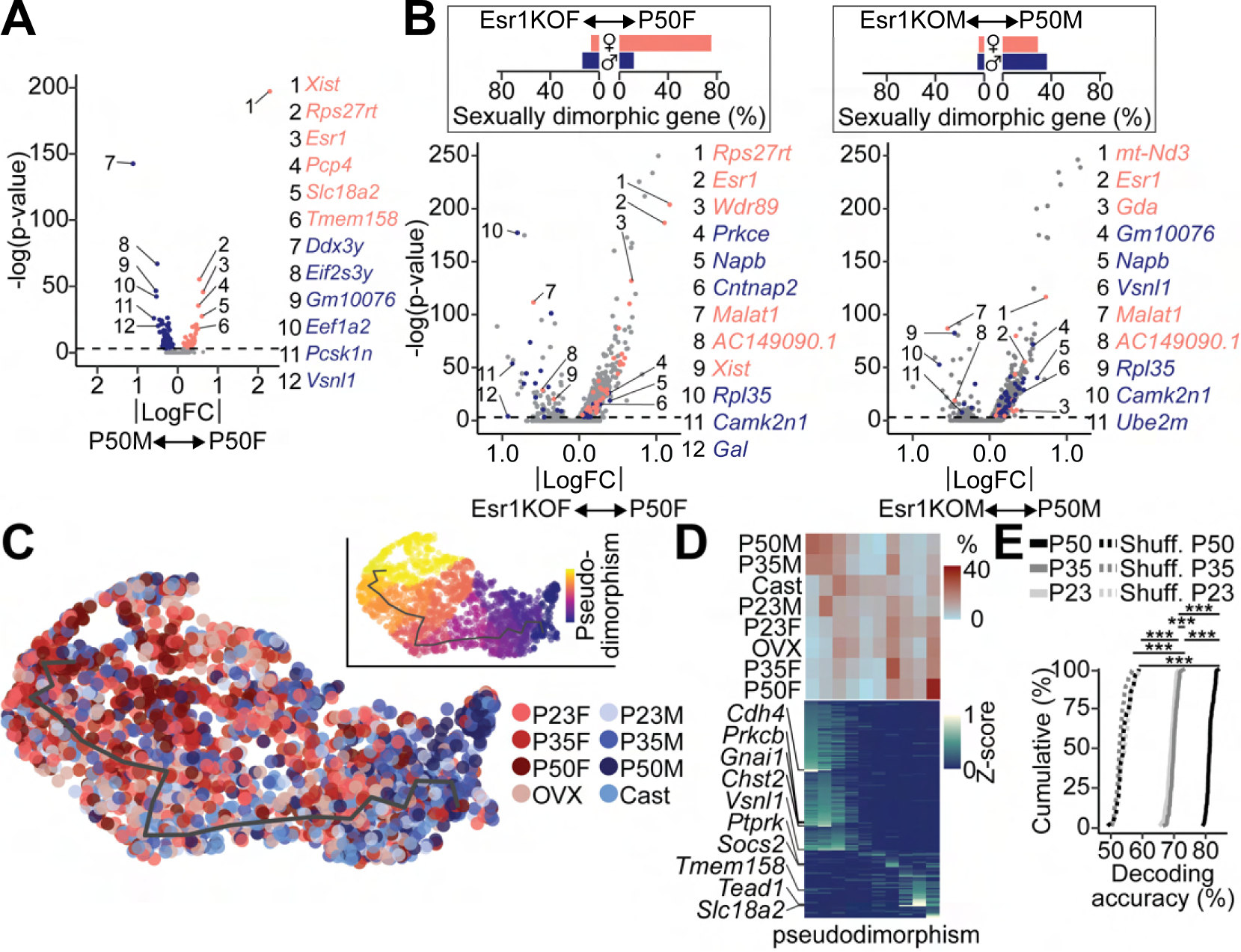
Development of transcriptional sexual dimorphism during puberty in MPOA^Vgat+ Esr1+^ neurons. (A) A volcano plot comparing P50 female and male gene expressions in Vgat^+^ Esr1^+^ populations. Dimorphic genes are highlighted (P50F rich: salmon, P50M rich: blue). (B) Left: A volcano plot comparing P50 female and Esr1KOF gene expressions in Vgat^+^ Esr1^+^ populations. The proportions of sexually dimorphic genes (P50F rich: salmon, P50M rich: blue), which are enriched in either intact P50F or Esr1KOF are quantified (top). Right: A volcano plot comparing P50 male and Esr1KOM gene expressions in Vgat^+^ Esr1^+^ populations. The proportions of sexually dimorphic genes (P50F rich: salmon, P50M rich: blue), which are enriched in either intact P50M or Esr1KOM are quantified (top). (C) UMAP visualization of transcriptional trajectory (black line) and cells (dots) color-coded by group (left) or pseudotime (right), computed using dimorphic genes in the Ers1-GRN. (D) Heatmaps showing proportion of cells (top) and scaled gene expression of sexually dimorphic genes (bottom) across dimorphic pseudotime. (E) Cumulative distributions of decoding accuracy between males and females (P50: black, P35: dark grey, P23: light grey, shuffled: dashed). One-way ANOVA followed by multiple comparisons. Details in Methods.

### The role of Esr1 in MPOA^Vgat^ neurons for the maturation of sexual behaviors

Because the MPOA is a critical circuit node for male and female sexual behaviors (Karigo et al., 2020; McHenry et al., 2017; Oomura et al., 1988; Wei et al., 2018), optogenetic stimulation of MPOA Vgat^+^ Esr1^+^ population is sufficient to elicit male sexual behaviors (Karigo et al., 2020), our transcriptional analysis showed that Vgat^+^ Esr1^+^ is a relevant population for sexual behaviors of both sexes (**Figure S2**), and Esr1 governs Esr1-GRNs in the Vgat^+^ Esr1^+^ in both sexes (**Figure5**), we assumed that hormone-associated pubertal transcriptional dynamics in the Vgat^+^ Esr1^+^ are crucial for pubertal maturation of mating behaviors. To directly test this hypothesis *in vivo*, we injected AAV-frtFlex-Cre into the MPOA of Vgat^Flp^ ::Esr1^lox/lox^ (Vgat-Esr1KO) or Vglut2^Flp^::Esr1^lox/lox^ (Vglut2-Esr1KO) mice of both sexes at P14-18 to delete *Esr1* in GABAergic or glutamatergic neurons for the entire puberty period and tested the maturation of sexual behaviors along with additional physiological and behavioral measures (**Figure 7A-P** **and S7A, B**). Neither the onset of puberty (BPS in males and VO in females) or the maturation of HPG axis (first day of estrous) were altered in either Vgat-Esr1KO and Vglut2-Esr1KO mice compared to controls (**Figure 7C, D, F, K, L, N**). However, maturation of sexual behaviors of both sexes was prevented in Vgat-Esr1KO mice (**Figure 7D, E, G, H**). In females, control group mice showed increased sexual receptivity as they approached adulthood while KO group showed significantly lower receptivity and delayed onset of the receptive age (**Figure 7D, E**). Similarly, in males, while control mice showed an increase in the number of mounting and intromitting behaviors towards adulthood, maturation of these sexual behaviors in the KO group were almost completely abolished (**Figure 7** **G, H**). In addition, there was a tendency for Vgat-Esr1KO mice to investigate female conspecific less than control mice (p=0.052) (**Figure S7A**). Interestingly, levels of aggression did not change by Esr1KO in the MPOA (number of attacks. KO: 11.6 ± 5.5, control:13 ± 7.5. unpaired t-test. p=0.8825). We did not see significant effects of investigative behaviors directed towards conspecifics in female Vgat-Esr1-KO mice. In both sexes of Vgat-Esr1KO mice, neither locomotion, anxiety-like behavior in the elevated plus maze test, or body weight were affected, suggesting that effects of Vgat-Esr1KO in the MPOA are specific to sexual behaviors. In contrast, consistent with the lack to activation in Vglut2 cells in the MPOA by sexual behaviors (**Figure S2**), Vglut2-Esr1KO mice had normal sexual behaviors, onset of puberty, locomotion, social preference, and elevated-plus maze activity by both sexes, except that Vglut2-Esr1KO mice had moderately increased body weight (**Figure 7L, M, O, P** **and Figure S7B**). Taken together, whereas previous studies showed the necessity of Esr1 in sexual behaviors by either global KO or non-cell type specific manipulations (Ogawa et al., 1998, 1997; Sano et al., 2013; Wu and Tollkuhn, 2017), our results show that Esr1 in the MPOA Vgat^+^ Esr1^+^ population is necessary for the pubertal maturation of sexual behaviors in both sexes.

## Discussion

In contrast to physiological functions and behaviors for survival including breathing, eating, drinking and other homeostatic functions needed throughout the lifespan, many brain functions required for emotion, cognition, and reproductive behaviors require secondary maturation during puberty for their full execution in adulthood (Blakemore et al., 2010, 2010; Piekarski et al., 2017; Romeo et al., 2002; Sigl-Glöckner et al., 2019). The pubertal maturation of behaviors is accompanied by brain-wide circuit modifications, which are likely governed by underlying molecular dynamics; however, previous studies did not profile pubertal transcriptional dynamics in the brain at single-cell resolution in any species. Our multimodal, transcriptional approaches revealed three important insights. First, in both males and females, neuronal cell types in the MPOA are already diversified before the onset of puberty. Cell types expressing both *Slc32a1* and *Esr1* (Vgat^+^ Esr1^+^) are relevant to sexual behaviors and transcriptionally active in both sexes. Second, pseudotime analysis on scRNAseq and HM-HCR FISH data consistently demonstrated that pubertal transcriptional dynamics in the single Vgat^+^ Esr1^+^ cells are a continuous progression towards the transcriptional states of adults, which could be delayed or facilitated by the levels of circulating sex hormones. Key HA-DEGs in the trajectory consist of both sexually monomorphic and dimorphic genes, which are sufficient for decoding whether each single cell is derived from mature (puberty, adult, hormonally supplemented) or immature (prepuberty, early puberty, hormonal deprivation) groups. Third, cis-regulatory analysis combined with *Esr1* KO showed that Esr1 causally regulates HA-DEGs and critical TFs, which compose a GRN centering around Esr1. A subset of Esr1-TFs are sexually dimorphic potentially orchestrating sexually dimorphic transcriptional states. Taken together, our multimodal analysis demonstrates that increasing levels of sex hormones during puberty govern pubertal transcriptional dynamics in the MPOA via activation of Esr1.

### Vgat-Esr1 neurons are transcriptionally dynamic during puberty and associated with mating behavior in adulthood

The MPOA is a highly heterogeneous structure with distinguishable cell types based on circuit connectivity, molecular markers, and their behavioral relevance. Indeed, previous scRNAseq studies identified 35 neuronal cell types in the avMPOA (Hrvatin et al., 2020) and 66 neuronal cell types in the anterior hypothalamus including MPOA (Moffitt et al., 2018). Consistent with the Hvatin et al, (2020) study, we found that 13,334 neurons reflecting of 20 Vgat-cell types and 7,658 neurons reflecting 12 Vglut2-cell types. Additionally, in agreement with microarray and scRNAseq studies on the development of the whole hypothalamus suggesting that cell-type-specific, marker genes are robustly expressed by P20 (Kim et al., 2020; Romanov et al., 2020; Shimogori et al., 2010), our study demonstrates that all the MPOA neuronal cell types expressing conserved marker genes, are consistently represented from prepuberty to adulthood, irrespective of their sex and hormonal states. As previous studies demonstrated that developmental diversification of cell lineages in the hypothalamus is determined by temporally dynamic expressions of a variety of unique TFs (Kim et al., 2020; Romanov et al., 2020), the establishment of diverse cell types in the MPOA is presumably governed cell-autonomously by TFs during prenatal and early postnatal development.

During puberty, our data imply that increasing levels of sex hormones induce secondary transcriptional dynamics to support pubertal maturation of brains. Indeed, we observed cell-type-specific induction of HA-DEGs in both Vgat^+^ and Vglut2^+^ cells from puberty to adulthood and an abundance of sex-hormone-related genes, such as *Esr1* and *Pgr*, in each cell type that predicts levels of transcriptional dynamics (**Figure2 and S1**), suggesting active roles of the sex hormone receptors in the regulation of pubertal transcriptome. Coincidently, our integrative analysis between published MERFISH (Moffitt et al., 2018) and our scRNAseq dataset revealed that a subset of cell types expressing both *Slc32a1* and *Esr1* are associated with sexual behaviors in both sexes. This is in line with the study by Karigo et al. (2020) showing that optogenetic stimulation of Vgat^+^ Esr1^+^ neurons in the MPOA is sufficient to elicit a variety of sexual behaviors in male mice. Compared to solid evidence of MPOA’s role in male sexual behaviors (Karigo et al., 2020; Wei et al., 2018), our analysis also shows that Vgat^+^ Esr1^+^ neurons in the MPOA are functionally relevant for sexual behaviors in both sexes.

### Hormonally linked transcriptional dynamics during puberty underlie a continuous progression towards an adult transcriptional state in the MPOA

During early developmental stages, progenitors first differentiate into precursors, which then develop into divergent terminally differentiated-cell types. Fates of progenitor and precursor cells are determined by cascades of several factors including cell signaling (Fukuchi-Shimogori and Grove, 2003, 2001; Lai, 2004; Lasky and Wu, 2005), neural activity (Luhmann et al., 2016) and spatiotemporally specific expression of TFs (Guillemot, 2007; Kim et al., 2020; Mayer et al., 2018; Romanov et al., 2020; Shimogori et al., 2010). Highly temporally specific expression of TFs causally regulates cell-type diversifications in the hypothalamus (Kim et al., 2020; Romanov et al., 2020). The majority of marker genes from diversified cell types are absent prior to late embryonic period and are only fully established by around P10-20 in the hypothalamus (Romanov et al., 2020; Shimogori et al., 2010), and cortical interneurons (Mayer et al., 2018). In contrast to early development, pubertal transcriptional alteration is minimal in the majority of MPOA neurons except for a subset of cell types enriched in *Esr1*, whose transcriptional state continuously progresses towards that of adulthood. In both the UMAP and PHATE space where global and local latent features of single cells are preserved (Becht et al., 2019; Burkhardt et al., 2021; Cao et al., 2019; Moon et al., 2019), the majority of pubertal and intact adult single Vgat^+^ Esr1^+^ neurons are spatially overlapping with similar transcriptional progression values (pseudotime or likelihood estimation) and highly segregated from cells from prepuberty and gonadectomized groups. SVM classification using the 30 top HA-DEGs in the scRNAseq data was sufficient to successfully decode whether neurons belonged to mature or immature groups while ∼11 genes in HM-HCR FISH were sufficient to reproduce pubertal transcriptional trajectory and decoding. Decoding accuracy incrementally increased as more DEGs were incorporated, suggesting that pubertal transcriptional states are globally determined by the combinatorial expression of HA-DEGs.

In humans, the most common mechanism for precocious puberty is the enhanced HPG activity (higher serum sex hormones and gonadotropins) (Carel and Léger, 2008) while around 40% of delayed puberty is associated with hypogonadotropic hypogonadism (Sedlmeyer and Palmert, 2002). Consistently, our pseudotime analysis on scRNAseq and HM-HCR data demonstrate that pubertal transcriptional progression in the MPOA is bidirectionally modulated by the levels of circulating sex hormones, estrogen in females and testosterone in males. Given that testosterone undergoes aromatization to estradiol, our data suggest that Esr1 acting alone or in conjunction with Ar serves as a master regulator of pubertal transcriptional progression. Indeed, stages of puberty are associated with enhanced development in emotion and cognition in humans (Goddings et al., 2012; Lawrence et al., 2015) and in mice (Anacker et al., 2020), suggesting similar mechanisms to what we describe in the MPOA occur in other brain regions.

### Sexually monomorphic and dimorphic TFs govern pubertal transcriptional dynamics

The MPOA exhibits sex differences in size, connectivity, and functions, which are likely supported by dimorphic transcriptional processes. Hormonal mechanisms at several developmental stages determine sexual dimorphism in the MPOA. First, an early postnatal surge of testosterone in males drives masculinization of the brain by regulating a variety of sexually dimorphic processes including apoptosis, gliogenesis and synaptogenesis (Gegenhuber and Tollkuhn, 2020; Wu et al., 2009). Indeed, testosterone administration at P0 in female mice masculinizes several sexually dimorphic brain regions in their sizes, cell numbers and neural connectivity (Gegenhuber and Tollkuhn, 2020; Wu et al., 2009). Increasing levels of circulating sex hormones during puberty are a second critical determinant for sex differences in the brain. A few lines of evidence suggest that changes in gene expression that begin during puberty may underlie the sexual dimorphism in the neural circuits and behaviors. Investigations of sexually dimorphic genes at prenatal, early childhood, puberty and adulthood stages in several brain regions from cortex to subcortical areas in humans, identified the largest number of dimorphic genes at puberty. Consistently, we found that HA-DEGs that start to increase during transition from puberty to adulthood are the majority of DEGs between all pair-wise comparisons in aggregated Vgat^+^ cells (**Figure1D**). Whereas male and female mice show some monomorphic patterns of pubertal transcriptional dynamics (**Figure 3** **and** **Figure 5**), Vgat^+^ Esr1^+^ cells are the most transcriptionally dynamic and dimorphic from puberty to adulthood. This process could be regulated by the levels of sex hormones in both sexes, and reflected in the substantial number of sexually dimorphic Esr1-DEGs (43-58 %).

Conventional gene regulation by Esr1 is mediated through direct DNA binding. However, Esr1 motif was enriched in only 15-16 % of Esr1-DEGs, among which 6 (female) and 3 (male) genes were dimorphic TFs. Further cis-regulatory analysis inferred that 37-56% of Esr1-DEGs were regulated by Esr1-TFs or Esr1 (**Figure 5F, G**). Thus, upon the elevation of circulating sex hormone levels during puberty, Esr1 facilitates the expression of critical TFs, which could then drive expression of diverse types of monomorphic and dimorphic genes. Importantly, expression levels of several sexually dimorphic Esr1-TFs e.g., *Sox5*, *Tcf4*, and *Pou6f2*, are elevated through puberty to adulthood, suggesting that sex differences in the transcriptome further extend beyond puberty. Indeed, trajectory analysis showed that transcriptional states of Vgat^+^ Esr1^+^ cells from adult males and females were the most distinct while prepuberty or hormonally deprived groups displayed intermediate states (**Figure 6C, D**). Sex differences in gene expression may also arise from sex chromosome linked genes, whose expression may also relate to steroid hormone levels. Further leveraging data from the current study may begin to unravel these interactions in a cell-type specific fashion. Collectively, in addition to sex differentiation regulated by the perinatal testosterone surge, puberty is an additional critical period where sexually dimorphic transcriptional states fully emerge.

### Maturation of neuronal circuitry for reproductive behaviors during puberty

The MPOA mediates a variety of homeostatic functions and behaviors, some of which require pubertal maturation, including sexual behaviors. Especially in males, there is extensive literature supporting its central role in the regulation of sexual behaviors across species (Karigo et al., 2020; Oomura et al., 1988; Wei et al., 2018). Recent advancements using optogenetics and imaging in mice have begun to identify mating-relevant neuronal populations in the MPOA. Wei et al. (2018) showed that optogenetic stimulation of *Esr1* expressing neurons in the MPOA is sufficient to elicit male-typical mounting behaviors. More recently, Karigo et al. (2020) advanced our understanding of male mating-relevant circuitry by elegantly demonstrating that MPOA Vgat^+^ Esr1^+^ cells regulate mating-related vocalization and mounting, but not aggression-related mounting. Though being investigated to a lesser extent in females, a population of neurons in the MPOA expressing *Nts*, which highly overlaps with Vgat^+^ Esr1^+^, drives socio-sexual behaviors in female mice (McHenry et al., 2017). Thus, Vgat^+^ Esr1^+^ cells are critical for sexual behaviors in both sexes. It is worth noting that *Nts*, *Gal*, or *Esr1* alone do not serve as reliable cell-type specific markers of hormonally responsive cell types (Karigo et al., 2020; McHenry et al., 2017; Wei et al., 2018; Wu et al., 2014). Future studies that manipulate neurocircuit function or measure activity will require an intersectional genetic approach to better understand the behavioral relevance of hormonally and molecularly defined cell types in the MPOA. It was unclear whether Esr1 signaling in Vgat^+^ neurons during puberty is critical for the maturation of sexual behaviors. Previous studies manipulating Esr1 in the MPOA did not distinguish cell-types, brain regions or developmental stages (Ogawa et al., 1998, 1997; Sano et al., 2016, 2013; Wu and Tollkuhn, 2017). Our data demonstrate that cell-type and region-specific deletion of *Esr1* in the MPOA severely disrupts the maturation of sexual behaviors in both sexes.

Although pubertal maturation of sexual behaviors was severely disrupted by Vgat-Esr1KO, other hormonally- and MPOA-dependent behaviors or physiological functions might also be regulated by pubertal transcriptional dynamics. Local application of estrogen in the MPOA activates mesolimbic reward circuitry via enhanced release of dopamine in the nucleus accumbens and can facilitate drug seeking behaviors in adult female rodents (Robison et al., 2018; Tobiansky et al., 2016, 2013), suggesting that motivated behaviors are broadly regulated by the levels of sex hormones. In addition, subpopulations of neurons expressing *Adcyap1*, *Slc17a6* or *Esr1* in the MPOA mediate thermoregulation and metabolism (Hrvatin et al., 2020; Tan et al., 2016; Veen et al., 2020). Consist with this, our Vglut2-Esr1KO moderately enhanced the gain of body weight during puberty (**Figure S7B**).

Dynamic adaptations in the circuit properties at several distinct levels, including cellular excitability, synaptic inputs and synaptic transmission to downstream targets are likely to synergistically contribute to pubertal maturation of mating circuitry. Consistent with electrophysiological studies showing changes in cellular excitability during puberty in several brain regions (Piekarski et al., 2017; Sabaliauskas et al., 2012; Srivastava et al., 2011; Yang et al., 2016), Vgat^+^ Esr1^+^ neurons in the MPOA gained excitability as a population through puberty (**Figure 2G**). Ontology analysis onto Esr1-DEGs revealed that genes supporting the axon structures and the machinery for synaptic transmission were highly enriched in both sexes (**Figure 1E** **and S5**). At network level, the MPOA receives dense monosynaptic inputs from brain nuclei enriched in sex-hormone receptors including the medial amygdala, the bed nucleus of the stria terminalis, and the ventromedial hypothalamus (Kohl et al., 2018), where circuit properties may alter during puberty via similar transcriptional dynamics as in the MPOA. Indeed, at least in adult mice, sex hormones regulate morphological (McEwen, 2002; McEwen and Woolley, 1994) or circuit properties at circuit nodes for social behaviors (Dey et al., 2015; Inoue et al., 2019; McHenry et al., 2017). Future studies are needed to better resolve brain-wide circuit connectivity potentially linking disparate groups of *Esr1*-expressing neurons.

## Conclusion

Our study is the first attempt to derive pubertal transcriptional dynamics in the brain at single cell resolution. Compared to embryonic and early postnatal development, the pubertal transcriptional progression is relatively minor except for in the cell types enriched with steroid-hormone receptors. Importantly, MPOA Vgat^+^ Esr1^+^ populations exhibit the most robust pubertal transcriptional dynamics, which is regulated by sexually monomorphic and dimorphic regulatory networks centered around Esr1. Similar pubertal transcriptional mechanisms at other circuit nodes may synergistically underlie the maturation of reproductive and other adult behaviors as well as contribute hormone-dependent neuropsychiatric disorders (Savell et al., 2020). Though the MPOA is molecularly and functionally heterogenous, transcriptional dynamics and its regulatory mechanisms identified in our study in conjunction with highly selective cell-type targeting technologies could facilitate comprehensive understanding of the relationships between genes, cell types and functions.

## Acknowledgments

We thank Stuber lab members for critical comments on the project and the manuscript. We thank C. Trapnell for helpful discussions. We thank Y. Tao and Z. Hu (University of North Carolina) for assistance with 10x genomics library generation. This work was supported by the Brain and Behavior Research Foundation (NARSAD Young Investigator Awards) (K.H. and M.A.R.), the National Institutes of Health DK121883 (M.A.R.), NS007431 (M.L.B.), DA038168 and DA032750 (G.D.S.), the Foundation of Hope and P30DA048736.

## Author Contributions

G.D.S. supervised the project. G.D.S. and K.H. conceived the project, designed experiments, analyzed and interpreted the data, and wrote the manuscript. K.H. wrote all the custom codes for the analysis of scRNAseq, MERFISH and HM-HCR data. K.H. and Y.H. conducted the majority of experiments. Y.H. analyzed HCR and slice-physiology experiments and interpreted the data. Y.L. conducted behavioral and histological experiments and analyzed the data. M.A.R. conducted the slice physiology experiments, assisted Y.H. with slice physiology analysis, assisted K.H. with decoding analysis and wrote the manuscript. M.L.B. analyzed the scRNAseq data. J.Y.C. conducted the slice-physiology experiments. O.R.A., R.V.M. and J.L.S. conducted image acquisition for histological experiments. J.A.M. and D.R.R. engaged in the conceptualization of the project. R.D.P. and L.S.Z. provided the split-Cre virus.

## STAR Methods

### KEY RESOURCES TABLE

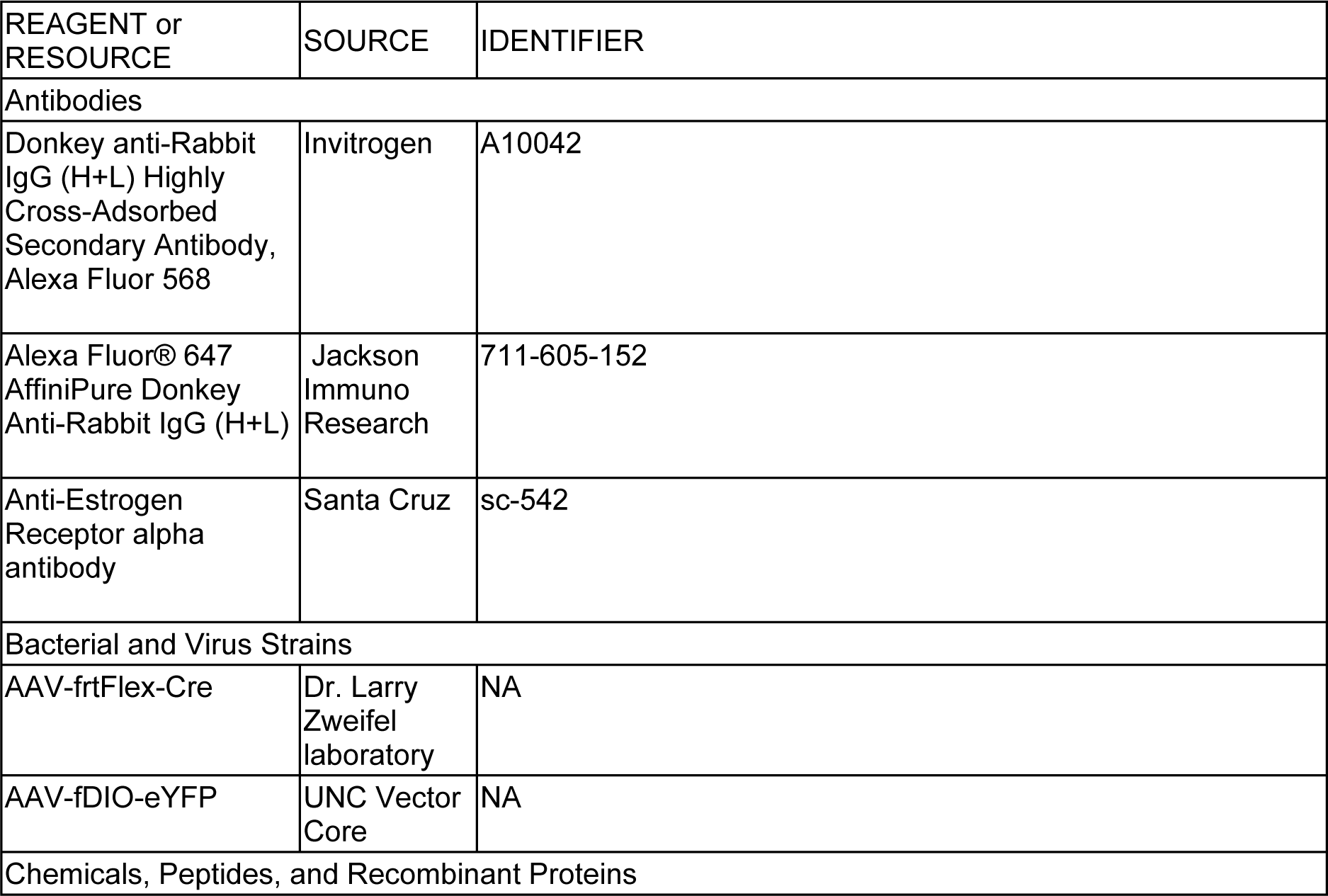
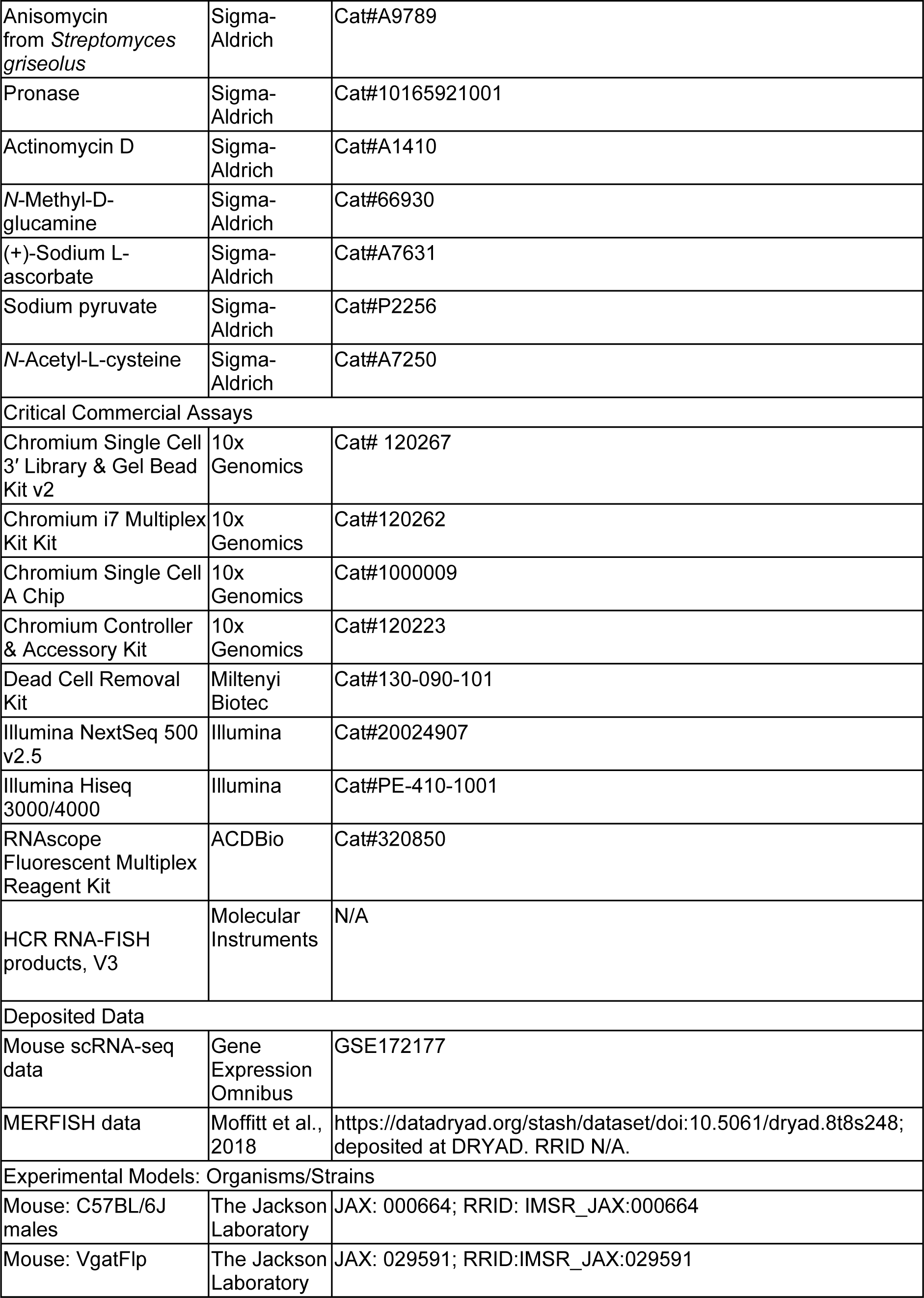
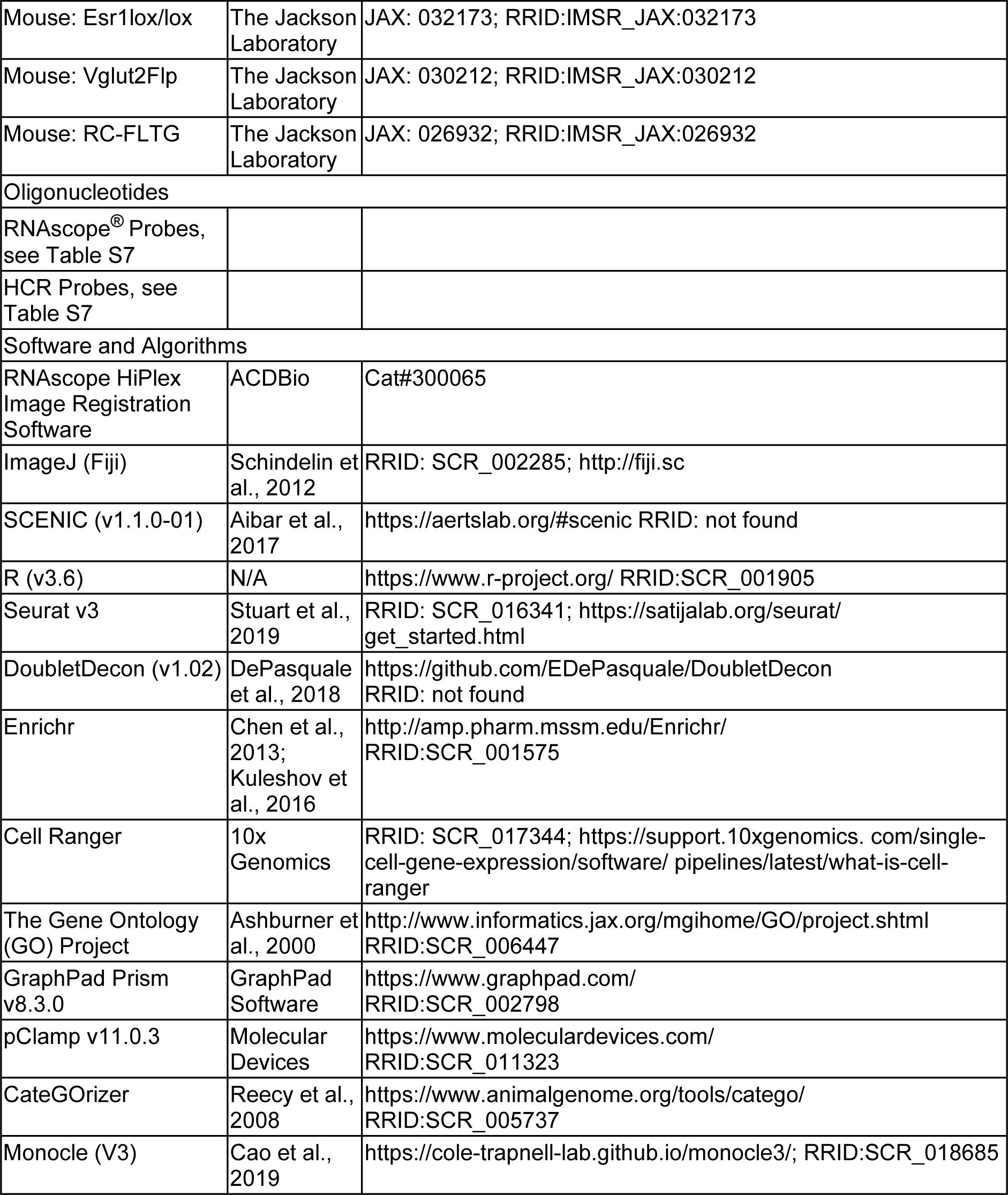

| REAGENT or RESOURCE | SOURCE | IDENTIFIER |
| --- | --- | --- |
| Antibodies |  |  |
| Donkey anti-Rabbit IgG (H+L) Highly Cross-Adsorbed Secondary Antibody, Alexa Fluor 568 | Invitrogen | A10042 |
| Alexa Fluor® 647 AffiniPure Donkey Anti-Rabbit IgG (H+L) | Jackson Immuno Research | 711-605-152 |
| Anti-Estrogen Receptor alpha antibody | Santa Cruz | sc-542 |
| Bacterial and Virus Strains |  |  |
| AAV-*frt*Flex-Cre | Dr. Larry Zweifel laboratory | NA |
| AAV-*fDIO*-eYFP | UNC Vector Core | NA |
| Chemicals, Peptides, and Recombinant Proteins |  |  |
| Anisomycin from *Streptomyces griseolus* | Sigma-Aldrich | Cat#A9789 |
| Pronase | Sigma-Aldrich | Cat#10165921001 |
| Actinomycin D | Sigma-Aldrich | Cat#A1410 |
| N-Methyl-D-glucamine | Sigma-Aldrich | Cat#66930 |
| (+)-Sodium L-ascorbate | Sigma-Aldrich | Cat#A7631 |
| Sodium pyruvate | Sigma-Aldrich | Cat#P2256 |
| N-Acetyl-L-cysteine | Sigma-Aldrich | Cat#A7250 |
| Critical Commercial Assays |  |  |
| Chromium Single Cell 3' Library & Gel Bead Kit v2 | 10x Genomics | Cat# 120267 |
| Chromium i7 Multiplex Kit Kit | 10x Genomics | Cat#120262 |
| Chromium Single Cell A Chip | 10x Genomics | Cat#1000009 |
| Chromium Controller & Accessory Kit | 10x Genomics | Cat#120223 |
| Dead Cell Removal Kit | Miltenyi Biotec | Cat#130-090-101 |
| Illumina NextSeq 500 v2.5 | Illumina | Cat#20024907 |
| Illumina Hiseq 3000/4000 | Illumina | Cat#PE-410-1001 |
| RNAscope Fluorescent Multiplex Reagent Kit | ACDBio | Cat#320850 |
| HCR RNA-FISH products, V3 | Molecular Instruments | N/A |
| Deposited Data |  |  |
| Mouse scRNA-seq data | Gene Expression Omnibus | GSE172177 |
| MERFISH data | Moffitt et al., 2018 | https://datadryad.org/stash/dataset/doi:10.5061/dryad.8t8s248; deposited at DRYAD. RRID N/A. |
| Experimental Models: Organisms/Strains |  |  |
| Mouse: C57BL/6J males | The Jackson Laboratory | JAX: 000664; RRID: IMSR_JAX:000664 |
| Mouse: VgatFlp | The Jackson Laboratory | JAX: 029591; RRID:IMSR_JAX:029591 |
| Mouse: Esr1lox/lox | The Jackson Laboratory | JAX: 032173; RRID:IMSR_JAX:032173 |
| Mouse: Vglut2Flp | The Jackson Laboratory | JAX: 030212; RRID:IMSR_JAX:030212 |
| Mouse: RC-FLTG | The Jackson Laboratory | JAX: 026932; RRID:IMSR_JAX:026932 |
| Oligonucleotides |  |  |
| RNAscope® Probes, see Table S7 |  |  |
| HCR Probes, see Table S7 |  |  |
| Software and Algorithms |  |  |
| RNAscope HiPlex Image Registration Software | ACDBio | Cat#300065 |
| ImageJ (Fiji) | Schindelin et al., 2012 | RRID: SCR_002285; http://fiji.sc |
| SCENIC (v1.1.0-01) | Aibar et al., 2017 | https://aertslab.org/#scenic RRID: not found |
| R (v3.6) | N/A | https://www.r-project.org/ RRID:SCR_001905 |
| Seurat v3 | Stuart et al., 2019 | RRID: SCR_016341; https://satijalab.org/seurat/get_started.html |
| DoubletDecon (v1.02) | DePasquale et al., 2018 | https://github.com/EDePasquale/DoubletDecon RRID: not found |
| Enrichr | Chen et al., 2013; Kuleshov et al., 2016 | http://amp.pharm.mssm.edu/Enrichr/ RRID:SCR_001575 |
| Cell Ranger | 10x Genomics | RRID: SCR_017344; https://support.10xgenomics.com/single-cell-gene-expression/software/pipelines/latest/what-is-cell-ranger |
| The Gene Ontology (GO) Project | Ashburner et al., 2000 | http://www.informatics.jax.org/mgihome/GO/project.shtml RRID:SCR_006447 |
| GraphPad Prism v8.3.0 | GraphPad Software | https://www.graphpad.com/ RRID:SCR_002798 |
| pClamp v11.0.3 | Molecular Devices | https://www.moleculardevices.com/ RRID:SCR_011323 |
| CateGORizer | Reecy et al., 2008 | https://www.animalgenome.org/tools/catego/ RRID:SCR_005737 |
| Monocle (V3) | Cao et al., 2019 | https://cole-trapnell-lab.github.io/monocle3/; RRID:SCR_018685 |

### RESOURCE AVAILABILITY

#### Lead Contact

Further information and requests for resources and reagents should be directed to and will be fulfilled by Garret Stuber.

#### Materials Availability

The plasmid generated in this study will be deposited to Addgene upon publication.

#### Data and Code Availability

The NCBI Gene Expression Omnibus accession number for the scRNAseq data reported in this paper will be available upon publication (https://www.ncbi.nlm.nih.gov/geo/query/acc.cgi?acc=GSE172177). All the codes used to analyze scRNAseq, and HM-HCR FISH are available at a Github repository affiliated with Stuber Laboratory group and this manuscript title (http://www.github.com/stuberlab/).

**Figure S1.**
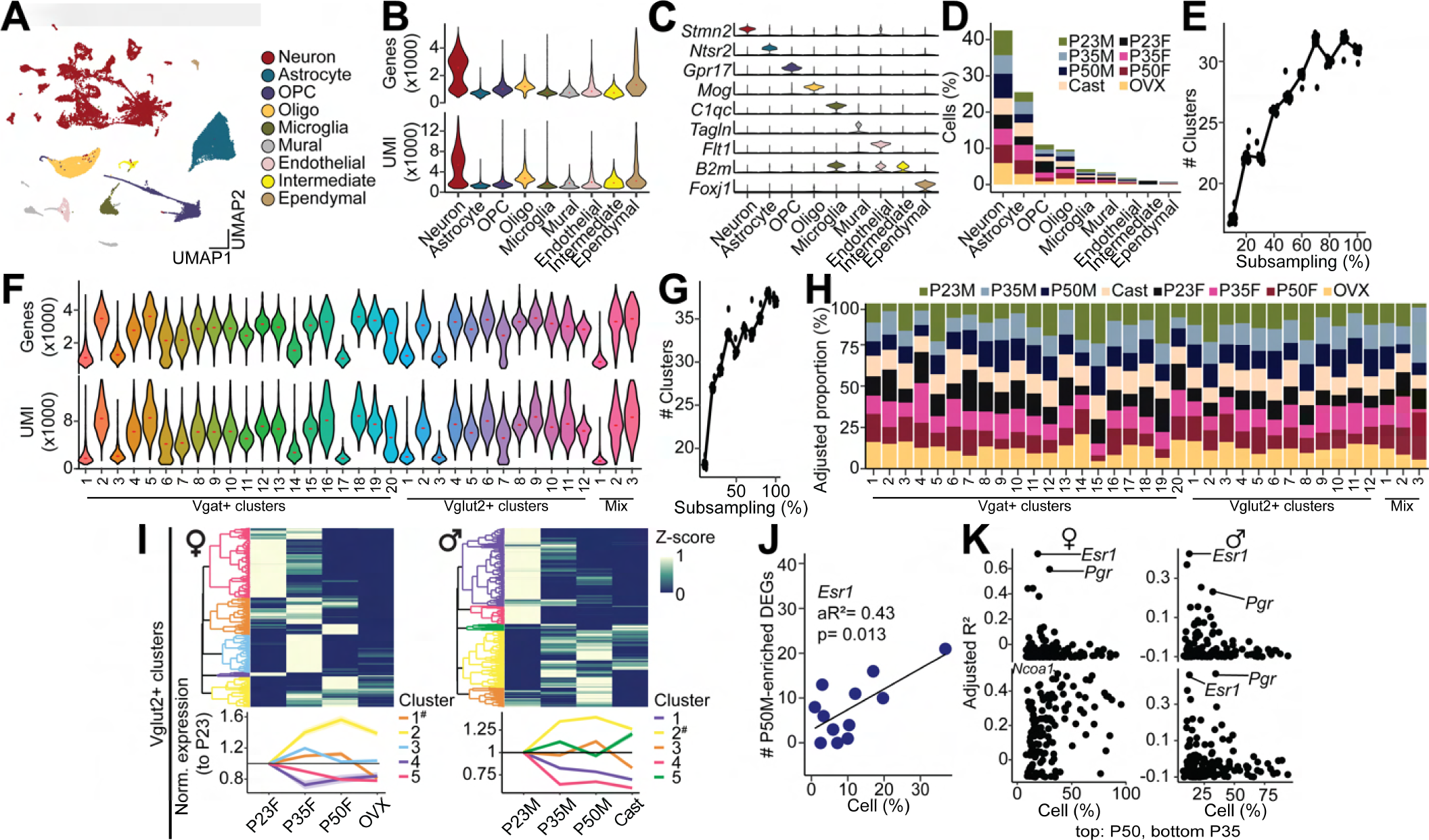
Basic supplementary information for scRNAseq experiments, related to Figure 1. (A) Joint clustering of 58,921 cells from all the groups identified 32 clusters. UMAP plot is color-coded by cell-category-type. (B) Violin plots showing the gene (top) and UMI (bottom) distributions in each cell-category-type. (C) Violin plots showing marker gene expression in each cell-category-type. (D) Proportion of cells from each group in each cell-category-type. (E) The number of clusters of sub-sampled MPOA cells. (F) Violin plots showing the gene (top) and UMI (bottom) distributions in each neuronal cell-type. (G) The number of clusters of sub-sampled MPOA neurons. (H) Fraction of cells from each group at each neuronal cell-type. (I) Top: heatmaps showing hierarchical clustering of DEGs in all Vglut2^+^ population in females (left) and males (right). Bottom: normalized expression of each gene cluster relative to the level of P23. Boldface clusters show higher expression in P35 and P50 than P23 and GDX. One-way repeated measures ANOVA followed by multiple comparisons. Detailed stats are in Methods. (J) An example for liner regression analysis between the percentage of Esr1 expressing cells and the number of P50M-enriched DEGs in comparison with GDX at each Vglut2^+^ cluster. Linear regression analysis. aR^2^: adjusted R squared. (K) Scatter plots showing adjusted R^2^ values of hormone receptors genes in females (left) and males (right) in comparison between P50 (top) or P35 (bottom) and GDX.

**Figure S2.**
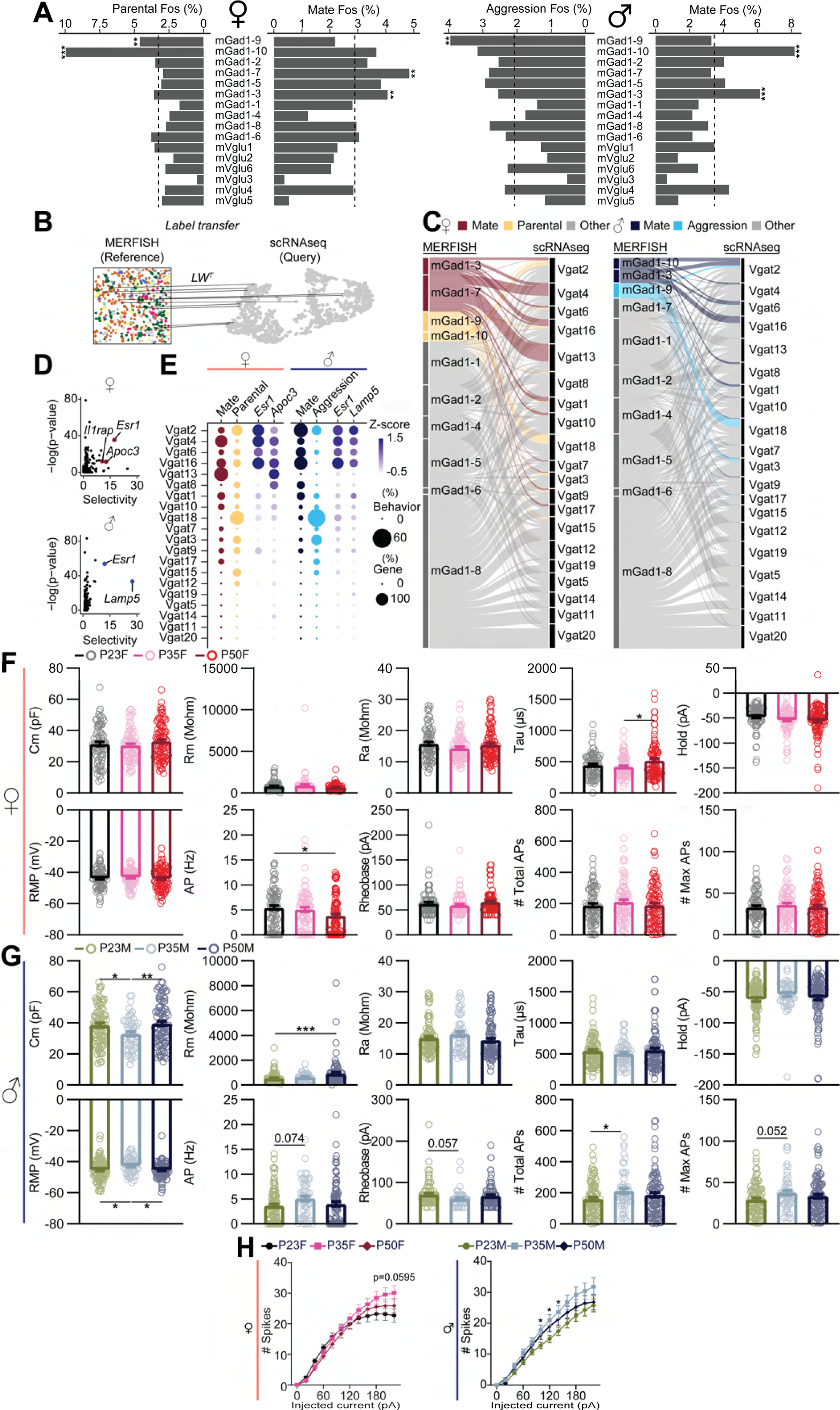
Basic supplementary information for scRNAseq and slice physiology experiments, related to Figure 2. (A) Fraction of *Fos* positive cells associated with behavioral types in each MERFISH cluster in female (top) and male (bottom). Dashed lines indicate mean percentages. Fisher’s exact test. **p < 0.01, ***p < 0.001. Statistical details in Methods. (B) Schematic illustrating integrative analysis to establish correspondence between MERFISH (Moffitt et al, 2018) and scRNAseq cell-types. (C) Alluvial plots showing the correspondence between MERFISH and scRNAseq clusters, color-coded by behavioral relevance. (D) Scatter plots showing enriched genes in mating-related scRNAseq clusters in females (top) and males (bottom). (E) Dot plots showing the fractions of socially relevant cells (indicated by the size of the dot) and two top marker genes in mating relevant clusters. Scaled gene expression is shown by color-coded intensity. Left: females, Right: males. (F, G) Upper left, membrane capacitance (Cm), membrane resistance (Rm), access Resistance (Ra), tau, holding current (Hold), resting membrane potential (RMP), base firing rate (AP), Rheobase, total number of firing during current injection, maximum number of firing during current injection in female (A) and male (B) Vgat^+^ Esr1^+^ cells from P23, P35 and P50. (H) Number of evoked spikes upon each current injection step in females (left) and males (right). Two-way repeated measures ANOVA followed by multiple comparisons. *p < 0.05. Details in Methods.

**Figure S3.**
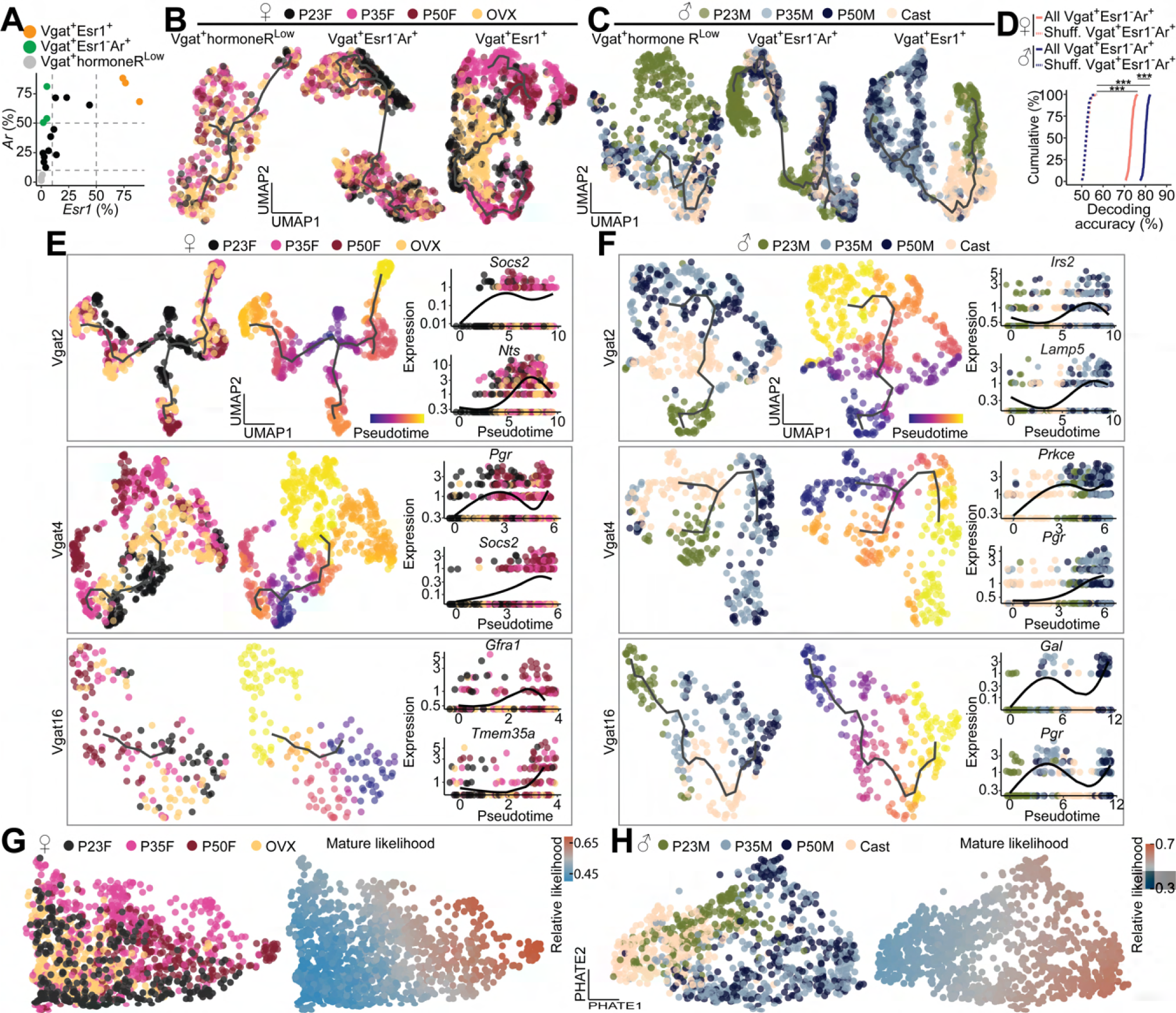
Supporting information for trajectory analysis on scRNAseq data, related to Figure 3. (A) A scatter plot showing the percentage of *Esr1* or *Ar* expressing cells in each Vgat cluster (dot). (B, C) UMAP visualization of transcriptional trajectories (black line) color-coded by experimental group in Vgat^+^ hormoneR^Low^ (left), Vgat^+^ Esr1^-^ Ar^+^(middle), Vgat^+^ Esr1^+^ (right) in female (B) and male (C). (D) Cumulative distributions of decoding accuracy by SVM classification between mature groups (P50, P35) and immature group (P23, GDX) using expression data from Vgat^+^ Esr1^-^ Ar^+^ (red: female, blue: male, real line), or shuffled data (dashed line). One-way ANOVA followed by multiple comparisons. Details in Methods. (E, F) UMAP visualization of transcriptional trajectory (black line) and cells (dots) color-coded by group (left), pseudotime (middle) in individual Vgat^+^ Esr1^+^ cluster of female (E) and male (F). (right) Kinetics plots showing relative expression of hormonally associated DEGs in females (E), males (F) across pubertal pseudotime in individual Vgat^+^ Esr1^+^ cluster. (G, H) PHATE visualization of transcriptional progression in Vgat^+^ Esr1^+^ cells (dot) color-coded by group (left) or relative mature likelihood (right) in female (G) or male (H). ***p < 0.001. Statistical details in Methods.

**Figure S4.**
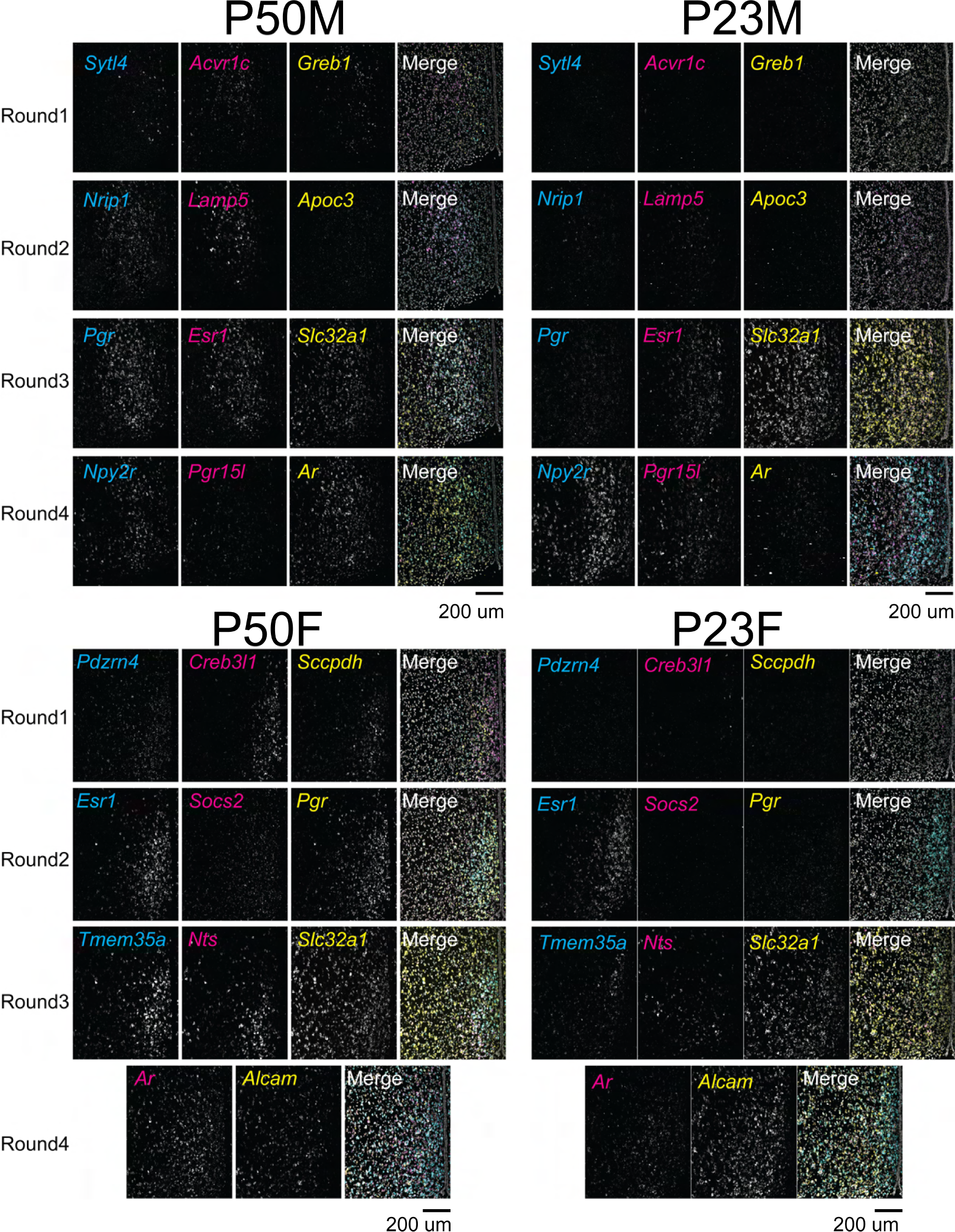
Supporting information for HM-HCR FISH, related to Figure 4. Representative images showing detected genes in 4 rounds of HCR FISH from P23 (left) and P50 (right) in female (bottom) and male (top). Scale bar: 200 µm.

**Figure S5.**
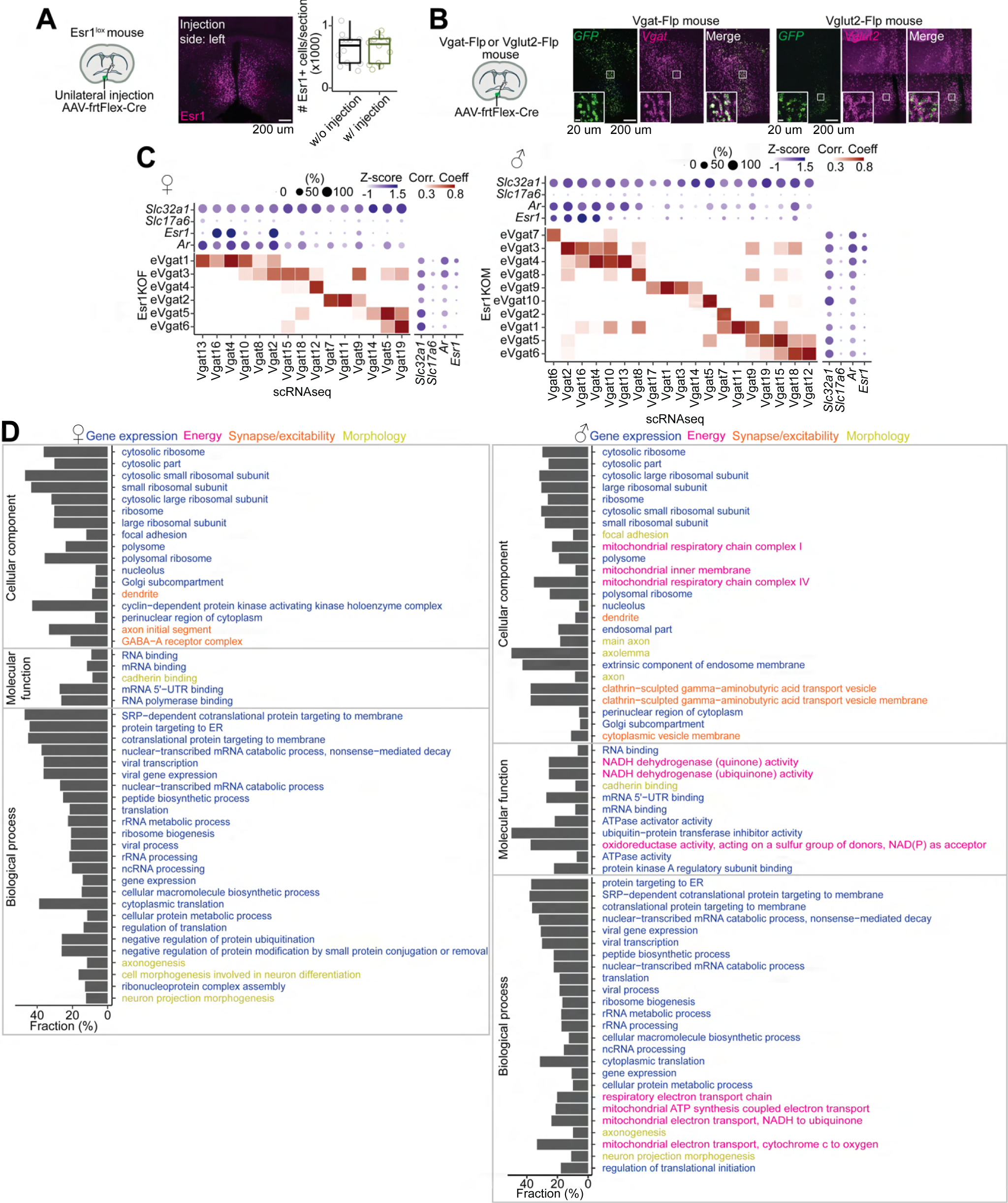
Supporting information for cell-type specific deletion of Esr1, related to Figure 5. (A) Left: schematic illustrating unilateral injection of AAV-frtFlex-cre virus in the MPOA of Esr1^lox/lox^ mice. Middle: a representative image showing the Esr1 expression in the MPOA. The virus was injected in the left. Right: quantification of Esr1 expression in the MPOA. Unpaired t-test. (B) Left: schematic illustrating AAV-frtFlex-Cre injection in the MPOA of Vgat^Flp^ or Vglut2^Flp^ mice. Representative images showing expression of *GFP* and *Slc32a1* (middle, Vgat^Flp^) or *Slc17a6* (right, Vglut2^Flp^) in the MPOA. Virally infected cells were largely overlapped with Slc32a1 (Vgat^Flp^, 96.3 %) or Slc17a6 (Vglut2^Flp^, 83.3 %). (C) Heatmaps illustrating Pearson correlation coefficient between Vgat clusters of Esr1KO and intact P50 groups. Dot plots are attached illustrating scaled expression levels (color) and the proportions of expressing cells (dot size) of *Slc32a1*, *Slc17a6*, *Esr1* and *Ar* in intact P50 (top) and Esr1KO (side) in female (left) and male (right). (D) Enriched gene ontology terms and fractions of all genes belonging to each term in Esr1-DEGs of female (left) and male (right). Up to 25 GO terms were shown and color-coded by gene category. GO Biological Process 2018, GO Molecular Function 2018, and GO Cellular Component 2018 were referenced. Statistical details in Methods.

**Figure S6.**
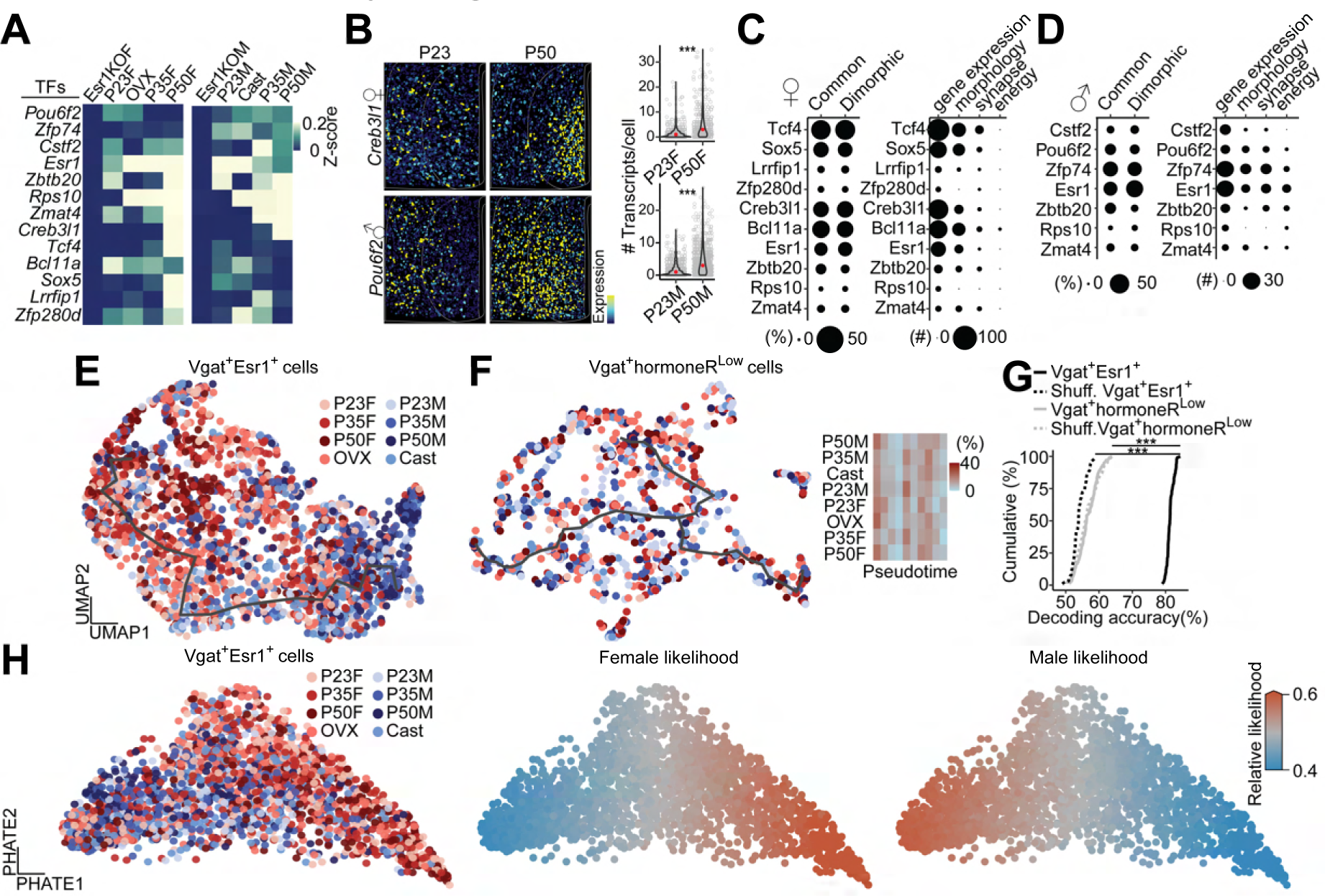
Supporting information for sexually dimorphic trajectory, related to Figure 6. (A) Heatmaps showing scaled expressions of Esr1 and Esr1-regulated TFs in female (left) or male (right) Vgat^+^ Esr1^+^ populations. (B) Representative images of reconstructed cells, color-coded by scaled expression of *Creb3l1* (female, top) or *Pou6f2* (male, bottom) in the MPOA from P23 (left) and P50 (middle). Right: violin plots showing the number of transcripts per cell. Wilcoxon rank-sum test. ***p < 0.001. (C, D) Dot plots illustrating the proportions of sexually monomorphic and dimorphic Esr1-DEGs regulated by each TF (left, dot size) and the number of regulated Esr1-DEGs in each gene category (right, dot size) in Vgat^+^ Esr1^+^ of female (**C**) and male (**D**). (E, F) UMAP visualization of transcriptional trajectory (black line) and cells (dots) color-coded by group in Vgat^+^ Esr1^+^ (**E**, also shown in Figure 5F) and Vgat^+^ hormoneR^Low^ (**F**), computed using dimorphic genes in the Ers1-GRN. (**F**, right) a heatmap showing proportion of cells across sexually dimorphic pseudotime. (G) Cumulative distributions of decoding accuracy by SVM classification between male and female P50 groups using expression data from Vgat^+^ Esr1^+^ (black real line), Vgat^+^ hormoneR^Low^ (grey, real line) or shuffled data (dashed line). One-way ANOVA followed by multiple comparisons. Details in Methods. (H) PHATE visualization of sexually dimorphic transcriptional transition in Vgat^+^ Esr1^+^ cells (dot) color-coded by group (left), relative female likelihood (middle) and relative male likelihood (right). ***p < 0.001. Statistical details in Methods.

**Figure S7.**
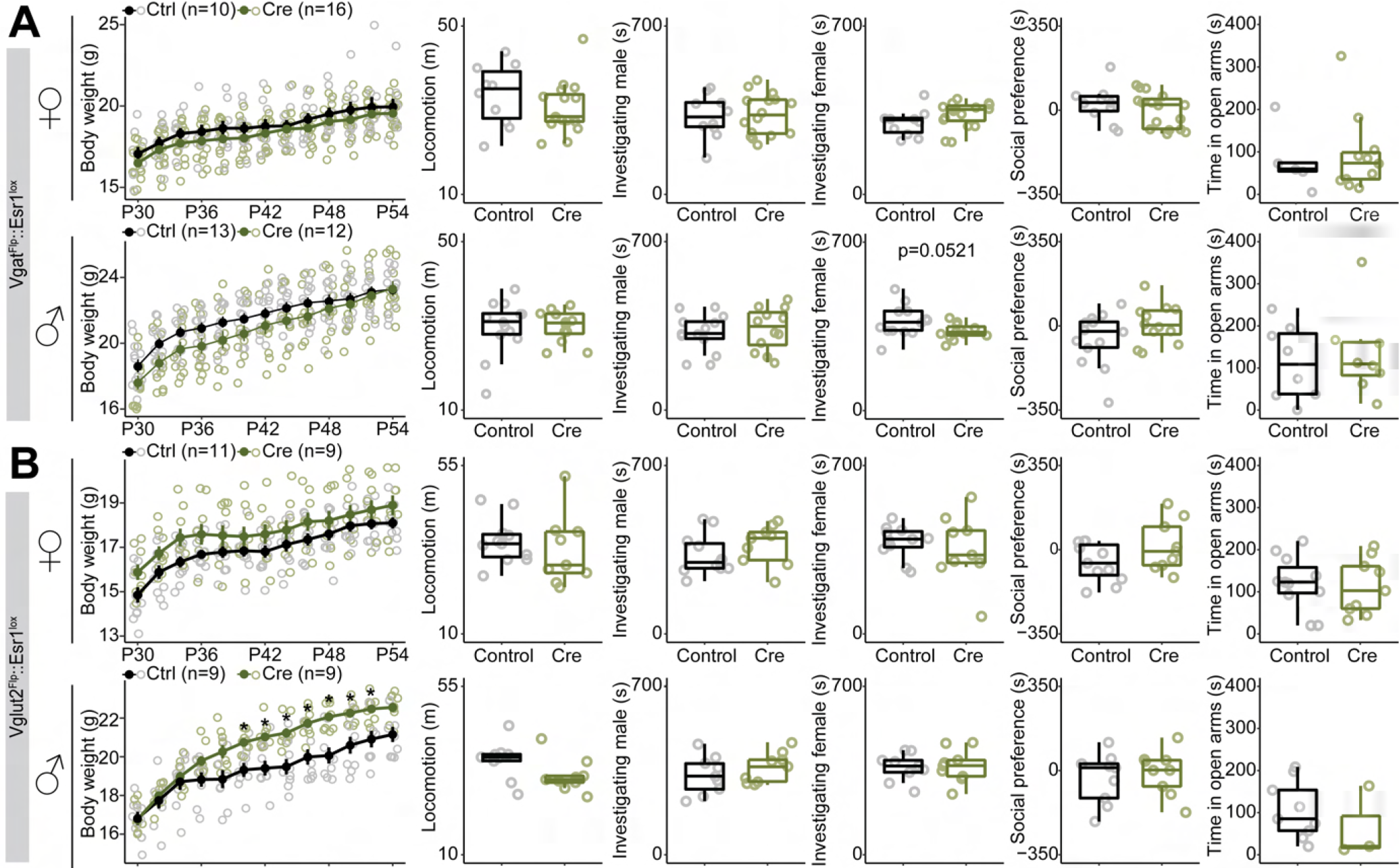
Supporting information for behavioral experiments, related to Figure 7. (A, B) From left to right: quantifications of body weight, locomotion, time investigating male, time investigating female, social preference and time spent in open arm of EPM in female (top) or male (bottom) of Vgat-Flp::Esr1^lox/lox^ (A) or Vglut2-Flp::Esr1^lox/lox^ mice (B). Line plots are shown in mean ± S.E.M. Box plots are shown with whiskers (min, max) and box (25%, median and 75%). ***p < 0.001. Statistical details in Methods.

Table S1. DEGs in aggregated Vgat^+^ or Vglut2^+^ clusters, related to Figure 1 and S1.

Table S2. DEGs in individual Vgat^+^ clusters, related to Figure 2.

Table S3. DEGs in mating-related clusters, related to Figure S2.

Table S4. DEGs in Vgat^+^ Esr1^+^ clusters comparing intact P50 or P35 and GDX, related to Figure 3.

Table S5. DEGs (sexually dimorphic genes) in Vgat^+^ Esr1^+^ clusters comparing intact P50M and P50F, related to Figure 5.

Table S6. DEGs in Vgat^+^ Esr1^+^ clusters comparing intact P50 and Esr1KO, related to Figure 5.

Table S7. Probe list for in situ hybridization.

### EXPERIMENTAL MODEL AND SUBJECT DETAILS

#### Mice

Several lines of male and female mice were used for scRNAseq, HM-HCR FISH, IHC, slice physiology and behavioral experiments (ages and detailed conditions will be provided in each section). All the mice used to generate data were on a C57BL/6 background. Wild-type C57BL/6J mice were originally obtained from Jackson Laboratory (JAX) and bred in-house. Vgat^Flp^ :: Esr1^lox/lox^ mice were generated by crossing Vgat^Flp^ (Jackson Laboratory stock no. 029591) and Esr1^lox/lox^ (kind gift from Kenneth Korach. Jackson Laboratory stock no. 032173) mice and then crossing them to Vgat^Flp^::Esr1^lox/+^ mice. Vglut2^Flp^::Esr1^lox/lox^ mice were generated by crossing Vglut2^Flp^ (Jackson Laboratory stock no. 030212) and Esr1^lox/lox^ mice and then crossing them with Vglut2^Flp^::Esr1^lox/+^ mice. For both Vgat^Flp^::Esr1^lox/lox^ and Vglut2^Flp^::Esr1^lox/lox^ mice, only the mice with heterozygous Flp and homozygous Esr1^lox/lox^ were used for experiments. Vgat^Flp^::Esr1^Cre^::RC-FLTG mice were generated by first generating heterozygous Vgat^Flp^ :: Esr1^Cre^ mice and then crossing female Vgat^Flp^::Esr1^Cre^ mice with a homozygous RC-FLTG male mouse (Jackson Laboratory stock no. 026932).For scRNAseq, HCR, and slice physiology experiments, mice were group housed except that for 1-2 days before tissue isolation, they were singly housed. Mice for behavioral experiments were singly housed during the experiments. Wildtype Swiss Webster (SW) mice were originally obtained from Taconic and bred in-house. Lactating SW mice were used for fostering pups that received brain surgery since original dams with C57BL/6 background frequently committed infanticide to their pups after the surgery. Mice had *ad libitum* access to food and water and were kept under a reverse 12 h light-dark cycle. All experiments were conducted in accordance with the National Institute of Health’s Guide for the Care and Use of Laboratory Animals and were approved by the Institutional Animal Care and Use Committee at the University of North Carolina and the University of Washington before the start of any experiments.

### METHOD DETAILS

#### Stereotaxic viral injection

Juvenile mice at P14-18 were separated from their dams, anaesthetized with isoflurane and placed in a stereotaxic frame (Kopf Instruments). The isoflurane anesthesia was maintained at <0.8%. After the craniotomy above target regions, viruses were injected using glass capillaries at a rate of 60 nl/min (Drummond, Nanoject). Injection was halted for 30 sec after every 60-nl infusion and capillaries were left in place for 10 min before retrieval. The target coordinate was 0.2 mm, 0.23 mm ML from Bregma and - 4.95 mm DV from brain surface. After the surgery and recovery from anesthesia, mice were transferred to a cage of lactating SW mice since lactating SW mice nursed juvenile mice that had received surgery while original dams with C57BL/6 background tended to commit infanticide after the surgery.

#### Gonadectomy

Gonadectomized groups were prepared for scRNAseq and HCR experiments. Mice at P22-23 were anaesthetized with isoflurane and placed in a stereotaxic frame (Kopf Instruments). The isoflurane anesthesia was maintained at <1.0%. After the hair was shaved around flank area (females) or scrotum (males), local anesthesia was administered, and a minimal incision was made on the skin and muscle above the target gonads. After the gonads were identified and exteriorized with forceps, they were dissected out using a heat cautery pen (Bovie Medical). Remaining tissues were placed back in the abdomen (females) or in the scrotum (males). The muscle and skin at the incision site were closed by suturing and an extra tissue adhesive (Vetbond) was applied onto the incision site to secure the closure of the skin. The brain tissues were collected at P50 ± 2 so that they were deprived of their sex hormones for the entire puberty period. Gonadectomized adult female mice were also prepared as stimulus mice for Resident Intruder (RI) and three chamber sociability assays. Identical procedure was conducted as above except that 7-8 weeks old C57BL/6J female mice were ovariectomized. After the 3 weeks recovery period, they were hormonally primed by the subcutaneous injection of β-Estradiol 3-Benzoate (Sigma, 10 µg at 48 hr and 5 µg at 24 hr before assay) and Progesterone (Sigma, 500 µg) at 6 prior to RI assays.

#### Generation of AAV-frtFlex-Cre virus

The split-Cre design was described in our previous studies (Chen et al., 2018; Heymann et al., 2020). In essence, an intron was introduced into the middle of Cre and the second exon was then flanked with frt sites; the action of FLP recombinase inverts the second exon and allows slicing to generate functional Cre. It was moved into a AAV vector for these studies.

The frtFlex-Cre-EGFP cassette was subcloned into pAAV-CAG-WPRE between AscI and XhoI. AAV1 virus for delivery of frtFlex-Cre-EGFP was generated as previously described (Heymann et al., 2020). Briefly, pAAV-CAG-frtFlex-Cre-EGFP-WPRE was transiently transfected into HEK293T/17 cells (ATCC) along with the pDG1 packaging plasmid. Twenty-four post-transfection cells were transferred to serum-free media and viral capsids were harvested 48 hrs later. AAV1 capsids were purified by gradient centrifugation and titre (∼2x10^12^ particles/mL) was estimated based on densitometry following gel electrophoresis relative to a known standard.

#### Validation of efficiency and specificity of AAV-frtFlex-split-Cre virus

To test the specificity of AAV-frtFlex-Cre, 300 nl of AAV-frtFlex-Cre was unilaterally injected into the MPOA of Vgat^Flp^ or Vglut2^Flp^ mice. Two weeks after the viral transduction, the brains were collected for FISH (detailed in RNAscope section) (**Figure S5B**). To test whether AAV-frtFlex-Cre would not express functional Cre without the presence of FLP, 300 nl of AAV-frtFlex-Cre was unilaterally injected into the MPOA of Esr1^lox/lox^ mice (n=3 each for male and female. Data from male and female subjects were combined). 6 weeks after the viral injection, the brain tissues were collected for IHC (detailed in Histology section) (**Figure S5A**). To test the efficiency, 300 nl of AAV-frtFlex-Cre was bilaterally injected into the MPOA of Vgat^Flp^::Esr1^lox/lox^ or Vglut2^Flp^::Esr1^lox/lox^ mice. All the mice for efficiency tests were first used for behavioral experiments (detailed in behavioral experiment section) and the quantitative results were reported for each mouse line and sex in **Figure 7** and **Figure S5**.

#### Single-cell preparation, cDNA library construction for scRNAseq

Male and female wild-type mice (n=3-4 per group) at P23 ± 1 (prepuberty group, hormonally intact), P35 ± 1 (mid-puberty group, hormonally intact), and P50 ± 2 (hormonally intact early adult or gonadectomy groups. Mice in gonadectomy groups were deprived of their gonads at P22-23) were singly housed 1-2 days prior to tissue dissection. We also prepared male and female Vgat-Flp :: Esr1^lox/lox^ mice, which were bilaterally injected with 300 nl of AAV-frtFlex-Cre at P14-18 in the MPOA, and their brain tissues were collected at P50 ± 2 (early adult) (n=3-4 per group). Subsequent steps largely followed previously published procedures (Hashikawa et al., 2020). Immediately after mice were removed from their home cage, they were deeply anesthetized with intraperitoneal injection of 0.2 mL of stock solution containing sodium pentobarbital (39 mg/mL) and phenytoin sodium (5mg/mL) and transcardially perfused with ice-old NMDG-aCSF containing inhibitor cocktails which was used throughout procedures unless noted. NMDG-aCSF: 96 mM NMDG, 2.5 mM KCl, 1.35 mM NaH2PO4, 30 mM NaHCO3, 20 mM HEPES, 25 mM glucose, 2 mM thiourea, 5 mM Na+ ascorbate, 3 mM Na+ pyruvate, 0.6 mM glutathione-ethyl-ester, 2 mM N-acetyl-cysteine, 0.5 mM CaCl2, 10 mM MgSO4; pH 7.35–7.40, 300-305 mOsm, oxygenated with 95% O2 and 5% CO2. Inhibitor cocktails: 500 nM TTX, 10 μM APV, 10 μM DNQX, 5 µM actinomycin, 37.7 µM anysomycin. The aCSF used for perfusion also contained transcription and translation inhibitors to minimize potential transcriptional evens introduced by pentobarbital injection, perfusion and brain extraction. Brains were extracted and coronal sections containing the MPOA were prepared at 300 µm using a vibratome (Leica, VT1200). Slices including the MPOA (3-4 slices/animal) were recovered in a chamber for 30 min on ice. MPOA tissues were then dissected using a micro scalpel (FEATHER) under the light microscope (LEICA, MZFL) from 3-4 animals/group (10-15 tissues/group) pooled and enzymatically digested with 1 mg/mL pronase (Sigma-Aldrich) for 30 min at room temperature, followed by mechanical trituration with fire-polished glass capillaries (tip diameter 200-300 µm) and filtered through strainers (pore size 40 µm) twice to remove cellular aggregates. Dead and dying cells in the cell suspension were removed using a dead cell removal kit (Miltenyi Biotec). After centrifugation, removal of supernatant and resuspension (20-30 μl), a fraction of cells (∼5 μl) were mixed with trypan blue to manually determine the proportion of live cells and cell concentrations using a hemocytometer. Only samples of high viability (>80%) were used for subsequent cDNA library preparation. Final cell concentration was adjusted to 800-1,000 cells/µL. We typically collected 30,000-80,000 cells in total from 3-4 mice, which well exceeded the number of cells required for cDNA library generation, so that biological variability between subjects was reduced.

cDNA libraries were constructed following manufacture’s instruction (Chromium Single Cell 3’ Reagents Kits V2 User Guide, 10x Genomics). Briefly, to recover up to 10,000 single cells, ∼17,000 dissociated cells were mixed with reverse transcription mix and loaded into the chip. The mRNAs of single cells were captured by barcoded beads in the droplets using a Chromium controller. Reverse transcribed cDNAs were then PCR amplified, fragmented, and ligated with adaptors followed by sample index PCR. cDNA libraries were sequenced on an Illumina Nextseq 500 (v2.5) or Hiseq and the alignment of raw sequencing reads to the mouse genome was conducted using the 10x Genomics Cell Ranger pipeline (V3) to obtain cell by gene-count matrices for subsequent downstream analysis.

#### Highly multiplexed HCR FISH

In addition to groups used in scRNAseq experiments including prepuberty (P23 ± 1), puberty (P35 ± 1), intact adult (P50 ± 2) and gonadectomy groups (P50 ± 2. They were deprived of their gonads at P22-23) from both sexes, we prepared a hormone-supplemented group that received a daily injection of sex hormones (Female: 5 µg β-Estradiol 3-Benzoate (Sigma) in 50 µl of sesame oil; male: 200 µg Testosterone Propionate (Sigma) in 50 µl of sesame oil) from P22-P27 and their tissues were collected at P28. A control group for hormone-supplemented group received a daily injection of sesame oil (50 µl) from P22 and P27 and their brains were harvested at P28. The physiological measures of puberty onset (BPS in male and VO in female) were examined daily by inspecting sexual organs for hormone-supplemented and control groups. It is important to note that all the subjects in hormone-supplemented groups showed BPS or VO by P28 while none of P28 control group were in puberty (detailed ages were provided in the main text). Mice in all 12 groups were singly housed for 1-2 days prior to tissue collection. Mice (n=2) were deeply anesthetized with isoflurane and brains were rapidly extracted and frozen on dry ice. Coronal sections were cut at 20 µm on a cryostat (Leica) and stored at -80°C until use. Sequential HCR fluorescent in situ hybridization (FISH) protocol was modified from the method described (von Buchholtz et al., 2021).

Tissue sample was fixed with pre-chilled 4% paraformaldehyde (PFA) for 30 min on ice, followed by quick rinse in 1x PBS twice at room temperature (RT). Tissue was dehydrated in various concentration of ethanol (50%, 70%, 100%, 100%, 5 min each) and air dried for 5 min at RT. After drawing contour with hydrophobic barrier pen, sections were permeabilized with protease IV (ACD, 322336) for 5 min at RT, then rinsed twice in 1x PBS and twice in 2x saline sodium citrate (2xSSC, Invitrogen, AM9763). For HCR FISH, all reagents (HCR version 3 probes, amplifiers and buffers) were purchased from Molecular Instruments unless noted (Choi et al., 2018). For hybridization, sample was first equilibrated in probe hybridization buffer for 10 min at 37°C inside the humidified chamber, then incubated in probe mixture, consisting of probes at a final concentration of 4 or 10 nM in probe hybridization buffer. The hybridization solution was covered with a coverslip and placed in the humidified chamber overnight at 37°C. The next day, coverslip was carefully removed in 100% wash buffer pre-warmed at 37°C. Sample was washed in pre-warmed wash buffer series [75%, 50%, 25% diluted in 5x SSCT (5x SSC, 0.1% Tween 20), 15 min each] and 5x SSCT (1x 15 min) at 37°C, then in 5x SSCT (1x 5 min) at RT. For HCR amplification, slices were equilibrated in amplification buffer for 30 min at RT and incubated in amplifier mixture consisting of hairpins conjugated with Alexa488, 546/594, and 647 in amplification buffer. Hairpins were prepared by snap-cooling; hairpin1 set and hairpin2 set were heated separately for 90 sec at 95°C and cooled down to RT for 30 min, then added to amplification buffer at final concentration of 60 nM. Samples were sealed with coverslips and incubated overnight at RT in a dark humidified chamber. The following day, coverslips were floated off in 5xSSCT at RT and washed in fresh 5x SSCT (2x 30 min), then rinsed twice in 2x SSC. Tissue autofluorescence was minimized using autofluorescence quenching kit (Vector laboratories, SP-8400-15) following manufacture’s user guide. Samples were treated with quenching solution (mixture of equal volume of reagent A, B and C) for 2 min at RT. After washing with 2x SSC for 5 min, nuclei were counterstained with Dapi solution (ACD, 320858) for 30 sec at RT, then mounted (Vector laboratories, H-1700-10). Dapi staining was performed only in the first round. Slides were stored at 4C until subsequent imaging. For optimal results, all images were acquired within 24 h of mounting.

##### Probe striping

After imaging, probes and amplifiers from the previous round were digested out before moving onto the next round of HM-HCR FISH. Samples were immersed in 2x SSC for >30 min at RT to float off coverslips and washed in 2x SSC (2x 5 min). Samples were incubated in DNase I (250 U/ml in 1x DNase I buffer, Roche, 04716728001) for 1.5 h at RT in a humidified chamber, then washed in 2x SSC (6x 5 min) and proceeded with the pre-hybridization step in the next round. Total 4 rounds of HCR-FISH allows detection of up to 12 different mRNAs.

##### Microscopy scanning

All images were acquired using epi-fluorescent microscopy AXIO Imager M2 and Zen software (Zeiss). 10x air objective was used. In addition to the three genes that were visualized in green (Alexa488), red (Alexa546/594) and far-red (Alexa647), brightfield images were acquired in the same field of view and used for image registration. Dapi signal in blue was imaged only in the first round and used to generate regions of interest (ROI) around nuclear. Individual channel was exported as tiff file.

##### Registration

Image registration was performed with HiPlex image registration software (ACD). First, all brightfield (BF) images were loaded and BF images from 2^nd^ - 4^th^ rounds were moved to that of round 1, generating round-specific transformation matrix. Rest of the images were transformed using the matrix file. Total up to17 images were overlayed and overlapping region was cropped to create a single 12-plex image.

##### Quantification

All Highly multiplexed HCR FISH images were analyzed with ImageJ. After registration, Dapi channel was used to generate regions of interest (ROI) containing nuclear in MPOA. Each ROI was then enlarged by few pixels to include cytoplasmic transcript. ROIs were transferred to the 11-12-plex image to measure number of transcripts for each gene.

#### RNAscope

AAV-frtFlex-Cre (300 nl) was unilaterally injected into the MPOA of Vgat^Flp^ or Vglut2^Flp^ mice (n=2 mice for each experiment). Two weeks after the viral incubation, subjects were deeply anesthetized with isoflurane and brains were rapidly extracted and frozen on dry ice. Coronal sections were cut at 20 µm on a cryostat (Leica) and stored at -80°C until use. The probe hybridization and signal detection procedures were performed following manufacture’s instruction (ACDbio). 2 genes (Vgat^Flp^: *Slc32a1*, *eGFP*; Vglut2^Flp^: *Slc17a6*, *eGFP*) were detected in each mouse line. The tiled Images containing MPOA were acquired with Zeiss ApoTome2 with 20x objective using Zen software (Zeiss). The settings for image acquisition were maintained for each experiment. The acquired CZI files were analyzed using Fiji for quantifying the number of *Scl32a1* or *Slc17a6*, and *eGFP* expressing cells and that of double-positive cells expressing *Scl32a1* or *Slc17a6*, and *eGFP* (**Figure S5B**).

#### Patch-clamp electrophysiology

Experimental methods were adapted from our previous study (Hashikawa et al., 2020; Rossi et al., 2019; Stamatakis et al., 2013). Male and female Vgat^Flp^::Esr1^Cre^::RC-FLTG mice (n=2-4 mice/group; at P23 ± 1, P35 ± 1 or P50 ± 2) were anesthetized with pentobarbital (0.39 mg/g body weight) and transcardially perfused with ice-cold sucrose cutting solution (in mM): NaCl 87, KCl 2.5, NaH2PO4 1.3, NaHCO3 25, MgCl2.6H2O 7, sucrose 75, CaCl2 0.5, 304-308 mOsm, saturated with 95% O2 and 5% CO2, pH 7.3-7.4. Brain was extracted rapidly and coronal section was prepared with vibratome (Leica VT1200) at 300 µm in the same cutting solution, then recovered in the holding chamber containing recording aCSF (in mM): NaCl 126, KCl 2.5, NaH2PO4 1.2, MgCl2.6H2O 1.2, NaHCO3 26, glucose 11, CaCl2 2.4, 304-308 mOsm, saturated with 95% O2 and 5% CO2, pH 7.3-7.4) for 1 h at 32 °C before recording. The slice was transferred to the recording chamber where the carbogenated recording aCSF was constantly perfused and warmed at 32°C. EGFP-expressing cells (putative Vgat^+^Esr1^+^ cells) in MPOA were identified under the rig microscope (Olympus, BX51WI) and targeted using borosilicate glass pipettes (Sutter Instrument, BF150-86-10, 3-5 MΩ) containing K-gluconate intracellular solution (in mM: K-gluconate 234.24, HEPES 238.31, KCl 74.56, mgATP 507.2, Na3GTP 523.2, k-gluconate and 1M KCl was added to adjust osmolarity and pH to 280-285 mOsm and pH7.3-7.4, respectively). Electrophysiological properties were measured using MultiClamp 700B and Clampex 11 (Molecular Devices).

Basal firing was recorded for 30-60 sec in I=0 mode, then switched to current clamp mode and cells were held at -70 mV. Rheobase was recorded by delivering 50-ms pulses in 10 pA steps (1 s/sweep). Current injection experiment was performed by injecting 800-ms current pulses stepwise from -100 pA to +220 pA with 20-pA increases (10 s/sweep). Cells that met following criteria were included in the analysis: giga-ohm seal was formed in cell-attach mode before break-in. Access resistance was lower than 30. Holding current was higher than -200 pA. Data analysis was done using pCLAMP11 (Molecular Devices). Cellular properties collected included: membrane capacitance (Cm), membrane resistance (Rm), access resistance (Ra), time constant (Tau), holding current, resting membrane potential (RMP), basal firing rate, rheobase, presence of burst firing during rheobase, number of action potentials during current injection experiment, presence of rebound firing during hyperpolarizing steps in current injection experiment.

#### Behavioral experiments

Male and female Vgat^Flp^::Esr1^lox/lox^ or Vglut2^Flp^::Esr1^lox/lox^ were bilaterally injected with 300 nl of AAV-frtFlex-Cre (Cre group. The virus was generated in-house. Detailed in Generation of AAV-frtFlex-Cre virus) or AAV-fDIO-eYFP (UNC Vector Core) at P14-18. From P25, the sexual organs were inspected daily to determine their first balanopreputial separation (male. BPS) or vaginal opening (female. VO) age. For female subjects, their vaginal smear was inspected daily after vaginal opening. All the behavioral experiments (Resident Intruder (RI) assays, locomotion, 3 chamber assays, and Elevated Plus maze) were conducted during the second half of the dark cycle in the room illuminated only with red light. The body weight was measured daily from P30-P54. Upon the completion of behavioral experiments, subjects were deeply anesthetized to extract brain tissues for histological experiments (detailed in Histology section). Subjects, which did not show increase in body weight or mistargeted viral injection were excluded from analysis.

#### Resident intruder assay to test pubertal maturation of sexual behaviors

Based on our pilot study on the maturation of sexual behaviors, we tested sexual behaviors of male subjects from P34 to P54 every other day. RI assay was adopted. Male subject mice were introduced with a hormonally primed adult female mouse to their home cage and freely interacted for 15 min (Vgat^Flp^::Esr1^lox/lox^ (cre), n=12; Vgat^Flp^::Esr1^lox/lox^ (control) n=13; Vglut^Flp^::Esr1^lox/lox^ (cre), n=9; Vglut^Flp^::Esr1^lox/lox^ (control), n=9). Importantly, to avoid ejaculation, male subjects and the female intruder were manually separated once the male subjects started thrusting. The RI test was recorded and the number of mounting and thrusting behaviors were manually counted. We also tested sexual receptivity of female subjects from P35 to P54 in Vgat^Flp^::Esr1^lox/lox^ (cre, n=16; control, n=10) and from P40 to P60 in Vglut2^Flp^::Esr1^lox/lox^ (cre, n=9; control, n=11), where the ages were determined by our pilot experiments. RI assay was conducted only when vaginal smear of female subjects indicated that they were in proestrus to estrous stages. Female subjects were introduced into the cage of singly housed and sexually experienced adult male mice and freely interacted for 10 min. Similar to RI assays in male subjects, to avoid ejaculation, female subjects were manually separated from the male residents once the male residents started intromission or about 5 s after the male started mounting without successful intromission. The RI test was video recorded with a head-mounted IR camera (IMAGINGSOUCE), which was controlled by Ethovision (Noldus) and the numbers of being mounted and being intromitted were manually counted. Receptivity was calculated as the fraction of intromitted bouts in the total mounted bouts. To minimize and equivalate the stress of female subjects across groups during RI test, the male residents were manually separated once they started aggression towards female subjects.

##### Sociability test

Sociability test was conducted after P45 on the days when RI test was not conducted. Subjects were placed in a standard three-chamber choice arena, where a caged social stimulus was present in one side of the chamber and an object was present in the other side of the chamber. Adult male or hormonally primed female mice were used for social stimuli. Subject mice were first habituated to the area without the presence of any stimuli for 5 min. After the habituation, a stimulus mouse and an object were introduced in the arena for 10 min. Then, after a stimulus mouse and an object were exchanged to a mouse of the opposite sex from the first test and a new object respectively, subject mice were freely exploring in the arena for another 10 min. Movements and locations of subjects during the habituation, first and second test sessions were tracked with a head-mounted IR camera (IMAGINGSOUCE) controlled by a live tracking software (Ethovision, Noldus). The total distance of movement during the habituation period was used to measure locomotion of subjects. The total time spent in the side of social stimulus chamber were used for statistical comparisons between groups.

##### Elevated plus maze

Elevated plus maze test (EPM) was conducted after P45 on the days when neither of RI test nor sociability test were conducted. Subjects were placed in a standard EPM arena (Rodriguez-Romaguera et al., 2020), where subject mice were allowed to freely explore the maze. Subject mice were first habituated to the maze for about 5 min. After the habituation, their locomotion was tracked with a head-mounted IR camera (IMAGINGSOUCE) controlled by Ethovision (Noldus). The total time spent in open or closed arms were used for statistical comparisons between groups.

#### Histology

Histological experiments were conducted 1) to examine the viral infection in the MPOA of subjects used in behavioral experiments (Vgat^Flp^:: Esr1^lox/lox^ or Vglut2^Flp^:: Esr1^lox/lox^ mice), 2) to validate specificity of AAV-frtFlex-Cre (Esr1^lox/lox^ mice) and 3) to validate the specificity of reporter gene expressions in Vgat^Flp^ :: Esr1^Cre^ :: RC-FLTG. Subjects were deeply anesthetized with pentobarbital and transcardially perfused with 40 ml of 4% PFA in PBS. The brains were extracted and post-fixed in 4 % PFA in PBS on ice for 5-7 hr. After the post-fixation, the brains were transferred to PBS with 0.05 % sodium azide (Sigma) and stored at 4 °C until sectioning. Free-floating coronal brain sections (50 µm) were obtained using a vibratome (Leica) and sections were stored in PBS with 0.05 % sodium azide until IHC. Sections were washed with PBS (3 x 5 min) and blocked in 15 % normal donkey serum (NDS) in PBST (0.3% triton) for 2 hrs at RT, followed by incubation with anti-Esr1 primary antibody (1:500, Santa Cruz, sc-542) in 15 % NDS in PBST for 72 hrs at 4 °C. Sections were washed with PBST (0.3 % triton, 3 x 30 min), incubated with secondary antibody (1:500, Life Technologies, donkey anti-rabbit 568 or Jackson ImmunoResearch, donkey anti-rabbit 647) in 15 % NDS in PBST for 2hrs at room temperature, washed with PBST (2 x 15 min), incubated with DAPI (2 min), washed with PBS (2x 15 min), mounted on slides and coverslipped with mounting medium. The tiled Images containing MPOA were acquired with Zeiss ApoTome2 with 20x objective using Zen software (Zeiss). The settings for image acquisition were maintained for each experiment. Acquired CZI files were analyzed using the HALO software with ISH-IF version 1.2 module (Indica Labs). In brief, the MPOA regions in both hemispheres were drawn as ROIs on three adjacent sections from each brain. Then, all GFP- or YFP-positive cells and Esr1-positive cells were automatically detected and counted using different threshold settings, and the double-positive cells were also counted by defining the cell phenotypes.

### QUANTIFICATION AND STATISTICAL ANALYSIS

#### Data Analysis for scRNAseq, MERFISH and HM-HCR data

Several R and Python analysis packages were adopted and modified for analyzing scRNAseq, MERFISH (previously published deposited data was acquired at DRYAD https://datadryad.org/stash/dataset/doi:10.5061/dryad.8t8s248) and HM-HCR FISH data. Clustering, differential gene expression analysis and integrative analysis of differential conditions and modalities were performed using the Seurat V3 package (Stuart et al., 2019). SCENIC package, consisting of GENIE3 (Aibar et al., 2017; Huynh-Thu et al., 2010) and RcisTarget (Aibar et al., 2017), was adopted to infer and deconstruct GRNs depicting cis-regulation of DEGs by transcription factors (Aibar et al., 2017; Davie et al., 2018). To learn progressions of transcriptional states by ages and hormonal states, we utilized Monocle V3 (Cao et al., 2019) and MELD (Burkhardt et al., 2021) packages. Ontology analysis was conducted using online software (Enrichr, https://maayanlab.cloud/Enrichr/)(Chen et al., 2013; Kuleshov et al., 2016). Data analysis consists of 4 major steps as described in detail each section below. 1) Clustering followed by identification of cell types which are transcriptionally dynamic during puberty and are relevant for reproductive behaviors. 2) Trajectory analysis to quantify pubertal transcriptional progression. 3) Transcriptional analysis in space. 4) Deconstruction of GRN of pubertal transcriptional dynamics.

#### Data preprocessing and doublet removal for scRNAseq data

Preprocessing procedure used in the previously published study was utilized (Hashikawa et al., 2020). Low expression genes (detected in less than 3 cells) and cells of low quality (total UMI <700 or total UMI>15,000, or percentage of mitochondrial genes >20%) were not included in downstream analysis. Suspected multiplet cells were computationally removed by utilizing DoubletDecon package (Version 1.1.5.) (DePasquale et al., 2019) with the default settings (doublet rate: 5.6 ± 1.0 %).

#### Integrative clustering and differential gene expression analysis

To minimize the effects of experimental variations (batch effects and effects of sex, ages and hormonal states) on clustering while retaining the global similarity of transcriptional states within the cell types, we used Seurat V3 integrative approach (Stuart et al., 2019), which utilized canonical correlation analysis (Butler et al., 2018) and mutual nearest neighbor analysis (Haghverdi et al., 2018). After preprocessing each data (58,921 cells in total), NormalizeData function was used to scale gene counts by the cellular sequencing depth (total UMI) with a constant scale factor (10,000) and then natural-log transformed (log1p). FindVariableFeatures function was used to select 2,000 highly variable genes in each sample based on a variance stabilizing transformation. For all pairwise datasets, FindIntegraiotnAnchors (CCA1-40) identified anchors with assigned scores and then IntegrationData computed integrated gene expression matrix by iteratively constructing and subtracting transformation matrix (*C*) from the original data matrix. Resulting integrated expression data was used for subsequent clustering. Integrated expression matrices were scaled and centered followed by principal component analysis (PCA) for dimensional reduction. Nearest neighbor graph was constructed in the PCA space (FindNeighbors, default setting) followed by Louvain clustering to identify clusters (resolution=0.8, FindClusters). For visualization of clusters, Uniform Manifold Approximation and Projection (UMAP) was independently generated using PCs (RunUMAP).

Initial clustering resulted in 32 clusters, among which 13 clusters were neuronal cells expressing *Thy1* or *Stmn2*. Based on the expression of canonical markers (Neuron: *Stmn2* or *Thy1;* Astrocyte: *Ntsr2;* OPC: *Gpr17;* Oligodendrocyte: *Mog,* Microglia: *C1qc,* Mural Cell: *Tagln,* Endothelial cell: *Flt1,* Intermediate cell: *B2m,* Ependymal cell: *Foxj1*) (Hashikawa et al., 2020; Tasic et al., 2018; Zeisel et al., 2018), neuronal cell-types and non-neuronal cell-types were determined and each cell types (neuron, astrocyte, oligodendrocyte, microglia, ependymal cells, OPC, intermediate cells and endothelial cells) were combined for the visualization in **Figure S1A-D**. To identify conserved markers (differentially expressed genes (DEGs) between clusters while being conserved across groups) for neuronal and non-neuronal clusters, FindConservedMarkers function was used to compare expression value of each gene in given clusters against the rest of cells with Wilcoxon rank sum test and p-values were adjusted with the number of genes tested. Genes with adjusted p-value <0.05 were considered to be significantly enriched. To examine the stability and robustness of clustering results, 10-100 % of cells were randomly sub-sampled and clustered by the identical procedure described above for 10 times at each sub-sampling rate. In order to obtain high-resolution census for neuronal cells, 24,831 cells in 13 neuronal clusters were extracted to be re-clustered using the identical integrative clustering analysis described above. Re-clustering identified 36 clusters, among which one cluster (204 cells) was excluded since that cluster expressed low levels of neuronal markers. Then, for the remaining 24,627 cells in 35 neuronal clusters, FindConservedMarkers function was used to identify marker genes in each cluster. 35 neuronal clusters were represented with all groups from different ages, sexes and hormonal states. To examine the DEGs between groups, first, FindMarkers was used to compute all the pairwise DEGs between groups in aggregate Vgat^+^ or Vglut2^+^ clusters in each sex (% of expression>10 %, logFC>0, adjusted p-value<0.05). Hierarchical clustering was conducted onto those DEGs to generate dendrogram trees where the same threshold (h=3.15) was used to identify clusters of DEGs. The hierarchical clustering identified groups of clusters which expressed higher in P50 and P35 groups than P23 and GDX in Vgat^+^ and Vglut2^+^ cells of both sexes. Similarly, we used FindMarkers function to further calculate various types of DEGs including hormonally associated DEGs (HA-DEGs) by comparing gene expressions between P50 or P35 and GDX groups in each cluster or various aggregated clusters (e.g. Vgat^+^ Esr1^+^, Vgat^+^ Esr1^-^ Ar^+^, Vgat^+^ hormone R^Low^. +: >50% positive cells, -: <10 % positive cells).

#### Integrative analysis of MERFISH and scRNAseq data

Seurat V3 was utilized to jointly analyze publicly available POA MERFISH data and our scRNAseq data. MERFISH dataset was acquired at DRYAD (Moffitt et al., 2018). Since we aimed to establish correspondence between MERFISH and scRNAseq clusters, related to reproductive behaviors in adult mice, cells from subjects, which did not undergo behaviors (Naïve, no *Fos* data) or were from lactating females or displaying aggression to a pup, were excluded. Then, cells categorized as “Inhibitory” or “Excitatory” in the metadata were extracted, and normalized with NormalizeData function. Highly variable genes were identified using FindVariableFeatures and top 60 variable genes (*Fos* was excluded) were used for subsequent clustering procedure described above. Seurat clustering identified 19 neuronal clusters in which 10 were GABAergic clusters enriched with *Gad1* and 6 were excitatory clusters expressing *Scl17a6*. Since MERFISH data does not have *Fos* data from behaviorally naïve animals, *Fos* enriched clusters were determined as the following. Per each behavioral category and sex, *Fos* enrichment threshold was calculated as the one-sided 95 percentile expression level. The proportion of cells above the threshold at each cell type was calculated and fisher’s exact test was performed to determine the *Fos* enriched clusters (p-values<0.05 after multicomparison correction). As the *Fos* enriched clusters were only present in a subset of GABAergic clusters, we only reported the correspondence of GABAergic clusters between modalities in subsequent analysis. In each sex, FindTransferAnchors (CCA1-30) and TransferData functions were used to identify anchors between reference (MERFISH) and query (scRNAseq, P50) data and to construct a weights matrix (*W*). Then, label (cell-type, classification) was transferred by *P = LW^T^*, where *L* is a binary classification matrix and *P* is a label prediction. Based on this imputed label of MERFISH clusters on cells in scRNAseq data, social-behavior related scRNAseq clusters were inferred. We used FindMarkers function to identify genes selectively enriched in scRNAseq clusters associated with sexual behaviors.

Integrative analysis of MERFISH and scRNAseq, and enrichment analysis of HA-DEGs consistently indicates that Vgat^+^ Esr1^+^ (proportion of Esr1^+^ cells>50%) clusters were transcriptionally dynamic and relevant for sexual behaviors. In the downstream analysis, we primarily focused on Vgat^+^ Esr1^+^ and, in case necessary, highlighted the differences with other Vgat^+^ populations (e.g. Vgat^+^ Esr1^-^ Ar^+^, Vgat^+^ hormone R^Low^).

#### Trajectory analysis of scRNAseq and HM-HCR FISH

Monocle 3 was utilized to quantify pubertal transcriptional progression for learning transcriptional trajectory in scRNAseq and HM-HCR FISH dataset (Cao et al., 2019). For scRNAseq data, Seurat clustering data was used to separately analyze Vgat^+^ Esr1^+^, Vgat^+^ Esr1^-^Ar^+^ and Vgat^+^ hormone R^Low^ in each sex. preprocess_cds function was used to normalize and scale data followed by PCA dimensional reduction using HA-DEGs (10-15 top PCs). reduce_dimension function was used to generate UMAP. principal graph in the UMAP space was learned using learn_graph function (default setting). Once the trajectory (principal graph) was learned, root node was determined by groups (either P23 or GDX). Pseudotime (measure of transcriptional progression from the root state) was computed by geodesic distance of each cell from the root node in the UMAP. Since we observed clear hormone-dependent pubertal transcriptional trajectory in the Vgat^+^ Esr1^+^, which consisted of 3 Vgat^+^ clusters, we repeated the same Monocle trajectory analysis on the individual Vgat^+^ Esr1^+^ clusters, Vgat2, 4, 16 (**Figure S3**). To test whether robust hormonally dependent transcriptional trajectory in Vgat^+^ Esr1^+^ cells is also observed in other cell types, similar monocle trajectory analysis was conducted to Vgat^+^ Esr1^-^ Ar^+^ and Vgat^+^ hormone _R_Low.

Similar to scRNAseq trajectory analysis, HM-HCR data was analyzed using Monocle 3. Trajectory was learned in Vgat^+^ Esr1^+^ cells with the identical procedure as the scRNAseq analysis except that log1p value, which was not normalized by total UMI, was used since the majority of detected genes were HA-DEGs and equivalent numbers of transcripts were not assumed in single cells.

To jointly learn transcriptional trajectories of scRNAseq and HM-HCR data, scRNAseq data was imputed from HM-HCR data using Seurat V3. In Vgat^+^ Esr1^+^ cells, *Slc32a1* and one of the genes in the HM-HCR data were removed from scRNAseq. Similar to the integrative analysis on MERFISH data, FindTransferAnchors (CCA) and TransferData functions were used to identify anchors between reference (HM-HCR) and query (scRNAseq) data and construct a weights matrix (*W*). Then, feature (gene expression) was transferred by *P = FW^T^*, where *F* is gene expression matrix and *P* is predicted gene expression. We iteratively repeated this procedure for all the HM-HCR genes to generate predicted scRNAseq data (11-12 genes). Predicted scRNAseq data was first validated with real data by Pearson correlation coefficient analysis. Then, transcriptional trajectory was learned on predicted data using the same Monocle trajectory analysis as above.

#### MELD analysis

To computationally cross validate monocle trajectory analysis of scRNAseq data, we utilized MELD python analysis package where transition of transcriptional state is learned as a relative likelihood of conditions in each cell in the PHATE space (Potential of Heat-diffusion for Affinity-based Trajectory Embedding) (Burkhardt et al., 2021; Moon et al., 2019). Preservation of local and distal relationships of transcriptional states of single cells in PHATE outperforms other embedding techniques (Moon et al., 2019). As input data, log normalized data and metadata (clustering) of Vgat^+^ Esr1^+^ cells generated by Seurat V3 analysis were used. First, to embed the data to low dimensional PHATE space, PCs were computed using scprep.reduce.pca (# components=8-15) and phate.PHATE and phate_op.fit_transform (default settings) were used to generate embedded data in PHATE. Second, kernel density of each condition (**Figure S3**: mature (P50, P35) or immature (P23, GDX). **Figure S6**: female (P50) or male (P50)) was estimated with meld.MELD (beta=67, knn=7) and meld_op.fit_transform (default setting) functions. The condition-associated relative likelihoods (relative likelihood that a cell would be observed in each condition) were computed using replicate_normalize_densities function (L1 normalization of the densities across samples). These MELD analyses were applied to measure hormonally associated pubertal trajectory (**Figure S3**) and sexually dimorphic trajectory (**Figure S6**).

#### Trajectory analysis of scRNAseq in Esr1KO experiments

Data of Esr1KO groups was first independently analyzed for preprocessing (removal of low quality cells and doublets) (DePasquale et al., 2019). Seurat V3 was utilized to perform clustering and identify neuronal clusters (detailed in **Integrative Clustering and differential gene expression analysis**). Neuronal cells were re-clustered to identify Vgat^+^ clusters. Then, correlation matrix was generated between datasets from Esr1KO and intact P50 groups in each sex by performing Person correlation coefficient analysis to determine Vgat^+^ clusters of Esr1KO group corresponding to Vgat2, 4, 16 of P50 group (Vgat^+^ Esr1^+^) (**Figure S5**). Vgat^+^ Esr1^+^ cells from all groups were combined for subsequent trajectory analysis using monocle 3. The same HA-DEGs were used to compute principal components, which will be used to construct UMAP. In the UMAP space, principal graph was learned, the root node was determined and pseudotime was computed (detailed in **Trajectory analysis of scRNAseq and HM-HCR FISH**).

#### Deconstruction of gene regulatory network

SCENIC is a powerful computational tool to infer the cis-regulation of genes by TFs (Aibar et al., 2017; Davie et al., 2018; Hashikawa et al., 2020). We adopted SCENIC and combined with cell-type specific gene manipulation to causally infer the GRN. First, using the P50 scRNAseq expression data of Vgat^+^ cells, GEINIE3 (runGenie3) was performed to compute co-expression scores between TFs and targeted genes (weights in the links). Then TFs were ranked by the sum of weights for HA-DEGs (logFC>0.25). In both female (952 TFs) and male (915 TFs), Esr1 had the highest ranked sum score among all the TFs in the database. This led us to perform scRNAseq on the subjects whose Esr1 was virally deleted in the Vgat^+^ cells in the MPOA (described in detailed above). After identifying Vgat^+^ Esr1^+^ clusters (described in

**Trajectory analysis of scRNAseq in Esr1KO experiments**), genes reduced by the Esr1 KO was computed using FindMarkers functions (Seurat V3, Esr1-DEGs). Next, to infer Esr1-DEGs which were cis-regulated by Esr1, RcisTarget package was utilized. RcisTarget database (mm9-tss-centered-10kb-7species.mc9nr.feather) was used for the scoring of the motif and annotation of TFs. Overrepresentation of each motif was calculated with calcAUC function. Based on the AUC distribution, significantly enriched motif was determined by the normalized enrichment score, which was calculated using addMotifAnnotation function (nesThreshold=2). 123 (female) and 98 (male) genes were found to be significantly enriched with Esr1 motif, among which 9 (female) and 6 (male) genes were TFs (Esr1-TFs). To test whether Esr1-TFs could cis-regulate Esr1-DEGs, we repeated RcisTarget analysis on the pair of Esr1-TFs and Esr1-DEGs. Ultimately, these analyses causally inferred genes cis-regulated by Esr1 or each Esr1-TF (regulons). Based on the regulon for each TF, igraph package was used to construct gene regulatory network (Esr1-GRN). In the Esr1-GRN, regulons were highlighted by the gene category (detailed in **Ontology analysis,** **Figure 5F, G**).

#### Ontology analysis

Enrichr was utilized to examine the enrichment of the functionally related genes in the HA-DEGs (**Figure1**) and Esr1-DEGs (**Figure S5**) (Chen et al., 2013; Kuleshov et al., 2016). GO Biological Process 2018, GO Molecular Function 2018, and Cellular Component 2018 were referenced to identify gene ontology terms (GO terms). Detailed GO terms were reported in **Figure S5**. GO terms were further grouped by 4 categories (Gene expression, Synapse/excitability, Morphology, Energy) and genes that were in multiple categories were manually assigned to one category. Since the number of GO terms in HA-DEGs of aggregated Vgat^+^ cells was extensive (female:144, male:405), The GO terms of HA-DEGs were further categorized into GO classes using CateGOrizer (55 in females; 61 males. 45 common GO classes were shown in **Figure 1**) (Reecy JM, 2008).

#### Clustering electrophysiological data

We conducted k-mean clustering to identify groups of cells in slice physiology data based on the electrophysiological properties, including membrane capacitance (Cm), membrane resistance (Rm), access resistance (Ra), time constant (Tau), holding current, resting membrane potential (RMP), basal firing rate, rheobase, presence of burst firing during rheobase, number of action potentials during current injection experiment, presence of rebound firing during hyperpolarizing steps in current injection experiment. 14-20 features of these features were used to perform PCA. fviz_nbclust function (factoextra package, method=”silhouette”) was used to determine optimal number of clusters for the k-mean clustering. Then, k-mean clustering was conducted using kmeans functions in R. The same PCs were used for the visualization of clusters in UMAP space.

#### Decoding conditions from scRNAseq, HM-HCR FISH or slice physiology data

To examine whether selected features in trajectory analysis for scRNAseq and HM-HCR FISH experiments or clustering analysis in slice physiology experiments were sufficient to decode the conditions of cells (age, sex, hormonal states, transcriptional maturity), we performed support vector machine (SVM) classification using scikit-learn with GridSearch cross validation as described in our previous studies (Otis et al., 2017; Rossi et al., 2019). SVM classifications were optimized using linear and rbf kernels across following parameters: γ: (10^-3^,10^-2^,10^-1^,10^0^,10^1^,10^2^,10^3^), *C*: (10^-3^,10^-2^,10^-1^,10^0^,10^1^,10^2^,10^3^). Input features for classifications were DEGs used in trajectory constructions in scRNAseq and HM-HCR FISH or electrophysiological features used for clustering in slice electrophysiology study. Decoding accuracies were compared with those calculated using randomized features (see Detailed Statical Procedures).

#### Detailed Statistical Procedures

Statistical details for each experiment are described in methods and below. Statistical analyses for electrophysiological data, behavioral data and SVM classification were conducted using GraphPad Prism v9.0.2. Non-parametric tests were performed to compare distributions of gene expression levels in scRNAseq and RNAscope data using R. No statistical tests were used to determine sample sizes.

**Figure 1D top:** one-way repeated measures ANOVA followed by multiple comparisons.

Cluster1: ANOVA, F (135, 405) = 2.9438259, p<0.0001. Holm-Sidak multi-comparison test: P23 vs OVX, t(135)= 0.073054964, p= 0.9419. P23 vs P35, t(135)= 6.6450382, p <0.0001. P23 vs P50, t(135)= 12.347378, p <0.0001. P35 vs OVX, t(135)= 4.0952302, p< 0.0001. P50 vs OVX, t(135)= 13.269141, p <0.0001. P35 vs P50, t(135)= 11.17 6780, p <0.0001.

Cluster2: ANOVA, F (282, 846) = 4.6794342, P<0.0001. Holm-Sidak multi-comparison test: P23 vs OVX, t(282)= 22.482676, p <0.0001. P23 vs P35, t(282)= 29.289408, p <0.0001. P23 vs P50, t(282)= 31.215903, p <0.0001. P35 vs OVX, t(282)= 5.2700226, p< 0.0001. P50 vs OVX, t(282)= 12.850838, p <0.0001. P35 vs P50, t(282)= 9.6808179, p <0.0001.

Cluster3: ANOVA, F (354, 1062) = 3.9834674, p<0.0001. Holm-Sidak multi-comparison test: P23 vs OVX, t(354)= 51.772642, p <0.0001. P23 vs P35, t(354)= 20.469338, p <0.0001. P23 vs P50, t(354)= 40.905873, p <0.0001. P35 vs OVX, t(354)= 28.730343, p< 0.0001. P50 vs OVX, t(354)= 11.831025, p<0.0001. P35 vs P50, t(354)= 20.702854, p <0.0001.

Cluster4: ANOVA, F (356, 1068) = 4.6642539, P<0.0001. Holm-Sidak multi-comparison test: P23 vs OVX, t(356)= 2.6050178, p <0.0096. P23 vs P35, t(356)= 23.110498, p <0.0001. P23 vs P50, t(356)= 6.1529826, p <0.0001. P35 vs OVX, t(356)= 33.395442, p< 0.0001. P50 vs OVX, t(356)= 11.259059, p <0.0001. P35 vs P50, t(356)= 30.387666, p <0.0001.

Cluster5: ANOVA, F (29, 87) = 6.4594382, p<0.0001. Holm-Sidak multi-comparison test: P23 vs OVX, t(29)= 3.9693469, p = 0.0017. P23 vs P35, t(29)= 1.4257659, p = 0.3021. P23 vs P50, t(29)= 0.040574193, p = 0.9679. P35 vs OVX, t(29)= 6.7801762, p< 0.0001. P50 vs OVX, t(29)= 7.3425963, p <0.0001. P35 vs P50, t(29)= 3.2891624, p = 0.0079.

Cluster6: ANOVA, F (339, 1017) = 3.7357024, p<0.0001. Holm-Sidak multi-comparison test: P23 vs OVX, t(339)= 27.925103, p < 0.0001. P23 vs P35, t(339)= 1.1969343, p = 0.2322. P23 vs P50, t(339)= 39.356609, p < 0.0001. P35 vs OVX, t(339)= 26.966204, p< 0.0001. P50 vs OVX, t(339)= 24.477635, p <0.0001. P35 vs P50, t(339)= 41.561976, p < 0.0001.

**Figure 1D bottom:** one-way repeated measures ANOVA followed by multiple comparisons.

Cluster1: ANOVA, F (770, 2310) = 4.8206939, p<0.0001. Holm-Sidak multi-comparison test: P23 vs Cast, t(770)= 12.113758, p < 0.0001. P23 vs P35, t(770)= 9.6124052, p < 0.0001. P23 vs P50, t(770)= 26.901991, p < 0.0001. P35 vs Cast, t(770)= 28.711090, p< 0.0001. P50 vs Cast, t(770)= 54.978801, p <0.0001. P35 vs P50, t(770)= 31.862981, p < 0.0001.

Cluster2: ANOVA, F (1089, 3267) = 5.6620007, p<0.0001. Holm-Sidak multi-comparison test: P23 vs Cast, t(1089)= 77.978449, p < 0.0001. P23 vs P35, t(1089)= 48.804692, p < 0.0001. P23 vs P50, t(1089)= 55.618808, p < 0.0001. P35 vs Cast, t(1089)= 36.331287, p< 0.0001. P50 vs Cast, t(1089)= 19.946091, p <0.0001. P35 vs P50, t(1089)= 10.943465, p < 0.0001.

Cluster3: ANOVA, F (189, 567) = 3.4899665, p<0.0001. Holm-Sidak multi-comparison test: P23 vs Cast, t(189)= 5.6579721, p < 0.0001. P23 vs P35, t(189)= 11.699347, p < 0.0001. P23 vs P50, t(189)= 8.0593044, p < 0.0001. P35 vs Cast, t(189)= 14.879007, p< 0.0001. P50 vs Cast, t(189)= 12.488563, p <0.0001. P35 vs P50, t(189)= 14.346162, p < 0.0001.

Cluster4: ANOVA, F (1030, 3090) = 6.0976581, p<0.0001. Holm-Sidak multi-comparison test: P23 vs Cast, t(1030)= 30.413779, p < 0.0001. P23 vs P35, t(1030)= 37.654898, p < 0.0001. P23 vs P50, t(1030)= 38.951160, p < 0.0001. P35 vs Cast, t(1030)= 11.231385, p< 0.0001. P50 vs Cast, t(1030)= 13.001623, p <0.0001. P35 vs P50, t(1030)= 5.5005486, p < 0.0001.

Cluster5: ANOVA, F (118, 354) = 3.3165807, p<0.0001. Holm-Sidak multi-comparison test: P23 vs Cast, t(118)= 6.3281474, p < 0.0001. P23 vs P35, t(118)= 7.1294177, p < 0.0001. P23 vs P50, t(118)= 6.5731275, p < 0.0001. P35 vs Cast, t(118)= 2.0573177, p= 0.0419. P50 vs Cast, t(118)= 11.363377, p <0.0001. P35 vs P50, t(118)= 14.575549, p < 0.0001.

**Figure 2B:** Linear regression analysis. Adjusted R-squared=0.51, p=0.00025.

**Figure 2C:** Linear regression analysis. Adjusted R-squared for each hormone receptor gene was reported.

**Figure 2I:** One-way ANOVA followed by multiple comparisons. ANOVA, F (3, 396) = 165.9, p<0.0001. Tukey’s multi-comparison test: real P50 vs shuffle P50, p< 0.0001. real P50 vs real P35, p< 0.0001. real P50 vs shuffle P35, p< 0.0001. shuffle P50 vs real P35, p< 0.0001. shuffle P50 vs shuffle P35, p= 0.9983. real P35 vs shuffle P35, p< 0.0001.

**Figure 2J:** One-way ANOVA followed by multiple comparisons. ANOVA, F (3, 396) = 165.9, P<0.0001. Tukey’s multi-comparison test: real P50 vs shuffle P50, p< 0.0001. real P50 vs real P35, p< 0.0001. real P50 vs shuffle P35, p< 0.0001. shuffle P50 vs real P35, p< 0.0001. shuffle P50 vs shuffle P35, p= 0.5252. real P35 vs shuffle P35, p< 0.0001.

**Figure 3D:** One-way ANOVA followed by multiple comparisons. ANOVA, F (3, 396) = 13016.7483, p<0.0001. Tukey’s multi-comparison test: real Vgat^+^ Esr1^+^ vs shuffle Vgat^+^ Esr1^+^ Vgat^+^, p< 0.0001. real Vgat^+^ Esr1^+^ vs real hormoneR^Low^, p< 0.0001. real Vgat^+^ Esr1^+^ vs shuffle hormoneR^Low^, p< 0.0001. shuffle Vgat^+^ Esr1^+^ vs real hormoneR^Low^, p< 0.0001. shuffle Vgat^+^ Esr1^+^ vs shuffle hormoneR^Low^, p< 0.0001. real hormoneR^Low^ vs shuffle hormoneR^Low^, p< 0.0001.

**Figure 3H:** One-way ANOVA followed by multiple comparisons. ANOVA, F (3, 396) = 21372.43634, p<0.0001. Tukey’s multi-comparison test: real Vgat^+^ Esr1^+^ vs shuffle Vgat^+^ Esr1^+^ Vgat^+^, p< 0.0001. real Vgat^+^ Esr1^+^ vs real hormoneR^Low^, p< 0.0001. real Vgat^+^ Esr1^+^ vs shuffle hormoneR^Low^, p< 0.0001. shuffle Vgat^+^ Esr1^+^ vs real hormoneR^Low^, p< 0.0001. shuffle Vgat^+^ Esr1^+^ vs shuffle hormoneR^Low^, p< 0.0001. real hormoneR^Low^ vs shuffle hormoneR^Low^, p< 0.0001.

**Figure 4I:** One-way ANOVA followed by multiple comparisons. ANOVA, F (3, 396) = 43666.7034, p<0.0001. Tukey’s multi-comparison test: real Vgat^+^ Esr1^+^ vs shuffle Vgat^+^ Esr1^+^ Vgat^+^, p< 0.0001. real Vgat^+^ Esr1^+^ vs real hormoneR^Low^, p< 0.0001. real Vgat^+^ Esr1^+^ vs shuffle hormoneR^Low^, p< 0.0001. shuffle Vgat^+^ Esr1^+^ vs real hormoneR^Low^, p< 0.0001. shuffle Vgat^+^ Esr1^+^ vs shuffle hormoneR^Low^, p< 0.0001. real hormoneR^Low^ vs shuffle hormoneR^Low^, p< 0.0001.

**Figure 4L:** One-way ANOVA followed by multiple comparisons. ANOVA, F (3, 396) = 36690.76438, p<0.0001. Tukey’s multi-comparison test: real Vgat^+^ Esr1^+^ vs shuffle Vgat^+^ Esr1^+^ Vgat^+^, p< 0.0001. real Vgat^+^ Esr1^+^ vs real hormoneR^Low^, p< 0.0001. real Vgat^+^ Esr1^+^ vs shuffle hormoneR^Low^, p< 0.0001. shuffle Vgat^+^ Esr1^+^ vs real hormoneR^Low^, p< 0.0001. shuffle Vgat^+^ Esr1^+^ vs shuffle hormoneR^Low^, p< 0.0001. real hormoneR^Low^ vs shuffle hormoneR^Low^, p< 0.0001.

**Figure 4Q:** unpaired t-test. T(198)=91.01, p<0.0001.

**Figure 4T:** unpaired t-test. T(198)=157.2, p<0.0001.

**Figure 6E:** One-way ANOVA followed by multiple comparisons. ANOVA, F (5, 594) = 5120.211650, p<0.0001. Tukey’s multi-comparison test: real P50 vs shuffle P50, p< 0.0001. real P50 vs real P35, p< 0.0001. real P50 vs shuffle P35, p< 0.0001. real P50 vs real P23, p< 0.0001. real P50 vs shuffle P23, p< 0.0001. shuffle P50 vs real P35, p< 0.0001. shuffle P50 vs shuffle P35, p< 0.0001. shuffle P50 vs real P23, p< 0.0001. shuffle P50 vs shuffle P23, p= 0.9968. real P35 vs shuffle P35, p< 0.0001. real P35 vs real P23, p= 0.0004. real P35 vs shuffle P23, p< 0.0001. real P23 vs shuffle P35, p< 0.0001. shuffle P35 vs shuffle P23, p< 0.0001. real P23 vs shuffle P23, p< 0.0001.

**Figure 7B:** Female: unpaired t-test. t(19)=5.599, p < 0.001., Male: unpaired t-test. t(18)=4.361, p=0.0004.

**Figure 7C:** unpaired t-test. t(24)=0.7317, p=0.4714.

**Figure 7D left:** unpaired t-test. t(24)=0.1302, p=0.8975.

**Figure 7D right:** unpaired t-test. Subjects, which did not become receptive by P55, were given the value 60. T(24)=4.138, p=0.0004.

**Figure 7E left:** Two-way repeated measures ANOVA followed by multiple comparisons. ANOVA revealed main effect of age (F (1.92778, 46.2667) = 21.8041, p<0.0001), main effect of group (F (1, 24) = 36.9300, p<0.0001) and interaction between group and age (F (2, 48) = 10.8824, p=0.0010). Holm-Sidak multi-comparison test was conducted. *p < 0.05, ***p < 0.001.

**Figure 7E right:** Two-way repeated measures ANOVA followed by multiple comparisons. ANOVA revealed main effect of age (F (1.88961, 45.3507) = 23.1245, p<0.0001), main effect of group (F (1, 24) = 75.2342, p<0.0001) and interaction between group and age (F (2, 48) = 10.3659, p=0.0002). Holm-Sidak multi-comparison test was conducted. *p < 0.05, **p < 0.01, ***p < 0.001.

**Figure 7F:** unpaired t-test. t(23)=0.09601, p=0.9243.

**Figure 7G:** unpaired t-test. t(23)=4.534, p=0.0001.

**Figure 7H left:** Two-way repeated measures ANOVA followed by multiple comparisons. ANOVA revealed main effect of age (F (3.94868, 90.8196) = 22.7095, p<0.0001), main effect of group (F (1, 23) = 66.9792, p<0.0001) and interaction between group and age (F (10, 230) = 18.7448, p<0.0001). Holm-Sidak multi-comparison test was conducted. *p < 0.05, **p < 0.01, ***p < 0.001.

**Figure 7H right:** Two-way repeated measures ANOVA followed by multiple comparisons. ANOVA revealed main effect of age (F (4.27919, 98.4214) = 13.4311, p<0.0001), main effect of group (F (1, 23) = 19.1087, p=0.0002) and interaction between group and age (F (10, 230) = 8.95218, p<0.0001). Holm-Sidak multi-comparison test was conducted. *p < 0.05, **p < 0.01.

**Figure 7J:** Female: unpaired t-test. t(18)=4.773, p=0.0002. Male: unpaired t-test. t(16)=4.531, p=0.0003.

**Figure 7K:** First VO age: unpaired t-test. t(18)=1.285, p=0.2152.

**Figure 7L left:** First estrous age: unpaired t-test. t(18)=0.5142, p=0.6133.

**Figure 7L right:** First receptive age: unpaired t-test. Subjects, which did not become receptive by P55, were given the value 65. t(18)=1.664, p=0.1134.

**Figure 7M left:** The number of being intromitted: Two-way repeated measures ANOVA. ANOVA revealed main effect of age (F (1.933, 34.79) = 10.92, p=0.0002), no effect of group (F (1, 18) = 2.471, p=0.1334) and no interaction between group and age (F (2, 36) = 0.5656, p=0.5730).

**Figure 7M right:** Receptivity: Two-way repeated measures ANOVA. ANOVA revealed main effect of age (F (1.990, 35.82) = 13.80, p<0.0001), no effect of group (F (1, 18) = 2.221, p=0.1534) and no interaction between group and age (F (2, 36) = 0.1445, p=0.8660).

**Figure 7N:** First BPS age: unpaired t-test. t(16)=1.000, p=0.3322.

**Figure 7O:** First mount age: unpaired t-test. t(16)=0.6030, p=0.5549.

**Figure 7P left:** The number of mounts: Two-way repeated measures ANOVA. ANOVA revealed main effect of age (F (3.494, 55.90) = 39.72, p<0.0001), no effect of group (F (1, 16) = 0.001556, p=0.9690) and no interaction between group and age (F (10, 160) = 0.5442, p=0.8566).

**Figure 7P right:** The number of thrusts: Two-way repeated measures ANOVA. ANOVA revealed main effect of age (F (3.831, 61.29) = 24.70, p<0.0001), no effect of group (F (1, 16) = 0.4016, p=0.5352) and no interaction between group and age (F (10, 160) = 0.3325, p=0.9713).

**Figure S1I left:** one-way repeated measures ANOVA followed by multiple comparisons.

Cluster1: ANOVA, F (106, 318) = 1.784, p<0.0001. Holm-Sidak multi-comparison test: P23 vs OVX, t(106)= 17.25, p<0.0001. P23 vs P35, t(106)= 5.938, p<0.0001. P23 vs P50, t(106)= 3.823, p =0.0004. P35 vs OVX, t(106)= 17.54, p< 0.0001. P50 vs OVX, t(106)= 9.249, p <0.0001. P35 vs P50, t(106)= 0.9242, p =0.3575.

Cluster2: ANOVA, F (87, 261) = 6.329, p<0.0001. Holm-Sidak multi-comparison test: P23 vs OVX, t(87)= 10.19, p<0.0001. P23 vs P35, t(87)= 8.772, p<0.0001. P23 vs P50, t(87)= 10.93, p<0.0001. P35 vs OVX, t(87)= 0.6321, p=0.5291. P50 vs OVX, t(87)= 7.370, p <0.0001. P35 vs P50, t(87)= 6.112, p<0.0001.

Cluster3: ANOVA, F (110, 330) = 2.172, p<0.0001. Holm-Sidak multi-comparison test: P23 vs OVX, t(110)= 4.682, p<0.0001. P23 vs P35, t(110)= 16.20, p<0.0001. P23 vs P50, t(110)= 3.059, p=0.0056. P35 vs OVX, t(110)= 11.90, p <0.0001. P50 vs OVX, t(110)= 1.708, p =0.0904. P35 vs P50, t(110)= 15.13, p<0.0001.

Cluster4: ANOVA, F (11, 33) = 6.390, p<0.0001. Holm-Sidak multi-comparison test: P23 vs OVX, t(11)= 3.374, p=0.0185. P23 vs P35, t(11)= 4.954, p=0.0026. P23 vs P50, t(11)= 3.743, p=0.0129. P35 vs OVX, t(11)= 4.328, p =0.0060. P50 vs OVX, t(11)= 1.324, p =0.2122. P35 vs P50, t(11)= 2.866, p=0.0305.

Cluster5: ANOVA, F (201, 603) = 4.559, p<0.0001. Holm-Sidak multi-comparison test: P23 vs OVX, t(201)= 29.20, p<0.0001. P23 vs P35, t(201)= 14.46, p<0.0001. P23 vs P50, t(201)= 32.37, p<0.0001. P35 vs OVX, t(201)= 26.25, p <0.0001. P50 vs OVX, t(201)= 2.931, p =0.0038. P35 vs P50, t(201)= 17.71, p<0.0001.

**Figure S1I right:** one-way repeated measures ANOVA followed by multiple comparisons.

Cluster1: F (706, 2118) = 5.635, p<0.0001. Holm-Sidak multi-comparison test: P23 vs Cast, t(706)= 52.31, p < 0.0001. P23 vs P35, t(706)= 28.25, p < 0.0001. P23 vs P50, t(706)= 33.39, p < 0.0001. P35 vs Cast, t(706)= 39.25, p< 0.0001. P50 vs Cast, t(706)= 19.27, p <0.0001. P35 vs P50, t(706)= 10.93, p < 0.0001.

Cluster2: ANOVA, F (520, 1560) = 5.850, p<0.0001. Holm-Sidak multi-comparison test: P23 vs Cast, t(520)= 13.78, p < 0.0001. P23 vs P35, t(520)= 21.64, p < 0.0001. P23 vs P50, t(520)= 22.69, p < 0.0001. P35 vs Cast, t(520)= 9.028, p< 0.0001. P50 vs Cast, t(520)= 10.51, p <0.0001. P35 vs P50, t(520)= 5.259, p < 0.0001.

Cluster3: ANOVA, F (176, 528) = 3.522, p<0.0001. Holm-Sidak multi-comparison test: P23 vs Cast, t(176)= 15.68, p < 0.0001. P23 vs P35, t(176)= 4.169, p < 0.0001. P23 vs P50, t(176)= 8.638, p < 0.0001. P35 vs Cast, t(176)= 17.51, p< 0.0001. P50 vs Cast, t(176)= 26.15, p <0.0001. P35 vs P50, t(176)= 13.70, p < 0.0001.

Cluster4: ANOVA, F (164, 492) = 5.971, p<0.0001. Holm-Sidak multi-comparison test: P23 vs Cast, t(164)= 29.20, p < 0.0001. P23 vs P35, t(164)= 33.47, p < 0.0001. P23 vs P50, t(164)= 31.91, p < 0.0001. P35 vs Cast, t(164)= 5.824, p< 0.0001. P50 vs Cast, t(164)= 9.851, p <0.0001. P35 vs P50, t(164)= 4.902, p < 0.0001.

Cluster5: ANOVAF (55, 165) = 3.797, p<0.0001. Holm-Sidak multi-comparison test: P23 vs Cast, t(55)= 6.306, p < 0.0001. P23 vs P35, t(55)= 6.165, p < 0.0001. P23 vs P50, t(55)= 1.518, p = 0.1348. P35 vs Cast, t(55)= 3.378, p= 0.0027. P50 vs Cast, t(55)= 11.23, p <0.0001. P35 vs P50, t(55)= 7.715, p < 0.0001.

**Figure S1J:** Linear regression analysis. Adjusted R-squared=0.43, p=0.013.

**Figure S1K:** Linear regression analysis. Adjusted R-squared for each hormone receptor gene was reported.

**Figure S2A:** Fisher’s exact test was conducted to compare each cluster with all clusters. P-values were Bonferroni corrected. **p < 0.01, ***p < 0.001.

**Figure S2F:** Top: One-way ANOVA followed by multiple comparisons. Cm: ANOVA, F (2, 252) = 1.191, p=0.3055. Rm: ANOVA, F (2, 252) = 1.859, p=0.1580. Ra: ANOVA, F (2, 252) = 1.814, p=0.1652. Tau: ANOVA, F (2, 252) = 4.066, p=0.0183. Tukey’s multi-comparison test was conducted. *p < 0.05. Hold: ANOVA, F (2, 252) = 2.464, p=0.0872. Bottom: One-way ANOVA followed by multiple comparisons. RMP: ANOVA, F (2, 252) = 0.4253, p=0.6541. AP: ANOVA, F (2, 252) = 4.094, p=0.0178. Tukey’s multi-comparison test was conducted. *p < 0.05. Rheobase: ANOVA, F (2, 252) = 1.681, p=0.1882. Current Inj Total AP No.: ANOVA, F (2, 252) = 0.9260, p=0.3975. Current Inj Max AP No.: ANOVA, F (2, 252) = 0.5893, p=0.5555.

**Figure S2G:** Top: One-way ANOVA followed by multiple comparisons. Cm: ANOVA, F (2, 236) = 4.990, p=0.0075. Tukey’s multi-comparison test was conducted. *p < 0.05, **p < 0.01. Rm: ANOVA, F (2, 236) = 6.969, p=0.0011. Tukey’s multi-comparison test was conducted. ***p < 0.001. Ra: ANOVA, F (2, 235) = 2.096, p=0.1253. Tau: ANOVA, F (2, 236) = 0.9338, p=0.3945. Hold: ANOVA, F (2, 236) = 1.237, p=0.2920. Bottom: One-way ANOVA followed by multiple comparisons. RMP: ANOVA, F (2, 236) = 4.934, p=0.0080. Tukey’s multi-comparison test was conducted. *p < 0.05. AP: ANOVA, F (2, 236) = 2.472, p=0.0866. Rheobase: ANOVA, F (2, 236) = 2.674, p=0.0711. Current Inj Total AP No.: ANOVA, F (2, 236) = 3.071, p=0.0482. Tukey’s multi-comparison test was conducted. *p < 0.05. Current Inj Max AP No.: ANOVA, F (2, 236) = 2.820, p=0.0616.

**Figure S2H left:** Two-way repeated measures ANOVA followed by multiple comparisons. ANOVA revealed main effect of time (F (1.776, 447.6) = 237.0, p<0.0001), no effect of group (F (2, 252) = 0.9118, p=0.4031) and interaction between group and time (F (22, 2772) = 2.270, p=0.0006). Tukey’s multi-comparison test was conducted.

**Figure S2H right:** Two-way repeated measures ANOVA followed by multiple comparisons. ANOVA revealed main effect of time (F (1.640, 387.1) = 248.4, p<0.0001), main effect of group (F (2, 236) = 3.071, p=0.0482) and interaction between group and time (F (22, 2596) = 1.282, p=0.1701). Tukey’s multi-comparison test was conducted. *p < 0.05.

**Figure S3D:** One-way ANOVA followed by multiple comparisons. ANOVA, F (3, 396) = 15075, p<0.0001. Tukey’s multi-comparison test: real male vs shuffle male, p< 0.0001. real male vs real female, p< 0.0001. real male vs shuffle female, p< 0.0001. shuffle male vs real female, p< 0.0001. shuffle male vs shuffle female, p= 0.6720. real female vs shuffle female, p< 0.0001.

**Figure S5A:** paired t-test. t(11)=0.08131, p=0.9367.

**Figure S6B:** Female: unpaired Wilcoxon test. W=197164, p<0.0001. Male: unpaired Wilcoxon test. W=506476, p<0.0001.

**Figure S6G:** One-way ANOVA followed by multiple comparisons. ANOVA, F (3, 396) = 2632, p<0.0001. Tukey’s multi-comparison test: real Vgat^+^ Esr1^+^ vs shuffle Vgat^+^ Esr1^+^ Vgat^+^, p< 0.0001. real Vgat^+^ Esr1^+^ vs real hormoneR^Low^, p< 0.0001. real Vgat^+^ Esr1^+^ vs shuffle hormoneR^Low^, p< 0.0001. shuffle Vgat^+^ Esr1^+^ vs real hormoneR^Low^, p< 0.0001. shuffle Vgat^+^ Esr1^+^ vs shuffle hormoneR^Low^, p< 0.0001. real hormoneR^Low^ vs shuffle hormoneR^Low^, p= 0.9331.

**Figure S7A top:** Body weight: Two-way repeated measures ANOVA. ANOVA revealed main effect of age (F (3.062, 73.49) = 51.24, p<0.0001), no effect of group (F (1, 24) = 1.366, p=0.2539) and no interaction between group and age (F (12, 288) = 0.2786, p=0.9922). Locomotion: unpaired t-test. t(24)=1.134, p=0.2682. Time investigating male: unpaired t-test. t(24)=0.2027, p=0.8411. Time investigating female: unpaired t-test. t(24)=1.499, p=0.1469. Social Preference: unpaired t-test. t(24)=0.8857, p=0.3846. Time in open arm: unpaired t-test. t(14)=0.2707, p=0.7906.

**Figure S7A bottom:** Body weight: Two-way repeated measures ANOVA followed by multiple comparisons. ANOVA revealed main effect of age (F (3.182, 73.19) = 202.5, p<0.0001), no effect of group (F (1, 23) = 1.788, p=0.1942) and interaction between group and age (F (12, 276) = 3.375, p=0.0001). Holm-Sidak multi-comparison test was conducted. No significance was observed at any age between groups. Locomotion: unpaired t-test. t(23)=0.06150, p=0.9515. Time investigating male: unpaired t-test. t(23)=0.5738, p=0.5717. Time investigating female: unpaired t-test. t(23)=2.048, p=0.0521. Social Preference: unpaired t-test. t(23)=1.563, p=0.1317. Time in open arm: unpaired t-test. t(14)=0.4188, p=0.6817.

**Figure S7B top:** Body weight: Two-way repeated measures ANOVA. ANOVA revealed main effect of age (F (4.942, 88.95) = 104.5, p<0.0001), no effect of group (F (1, 18) = 2.849, p=0.1087) and no interaction between group and age (F (12, 216) = 1.006, p=0.4442). Locomotion: unpaired t-test. t(18)=0.7283, p=0.4758.Time investigating male: unpaired t-test. t(18)=1.060, p=0.3030. Time investigating female: unpaired t-test. t(18)=0.5965, p=0.5583. Social Preference: unpaired t-test. t(18)=1.684, p=0.1094. Time in open arm: unpaired t-test. t(18)=0.2933, p=0.7727.

**Figure S7B bottom:** Body weight: Two-way repeated measures ANOVA followed by multiple comparisons. ANOVA revealed main effect of age (F (4.126, 66.02) = 197.4, p<0.0001), main effect of group (F (1, 16) = 8.571, p=0.0099) and interaction between group and age (F (12, 192) = 9.574, p<0.0001). Holm-Sidak multi-comparison test was conducted. *p < 0.05. Locomotion: unpaired t-test. t(16)=1.826, p=0.0865. Time investigating male: unpaired t-test. t(16)=0.9004, p=0.3813. Time investigating female: unpaired t-test. t(16)=0.1816, p=0.8582. Social Preference: unpaired t-test. t(16)=0.5261, p=0.6061. Time in open arm: unpaired t-test. t(10)=0.8410, p=0.4200.

